# Enzymatic formation of a conserved isoaspartate in ribosomal protein uS11

**DOI:** 10.64898/2026.08.26.747336

**Authors:** Yanqing Xue, Chandrima Majumdar, Salimat O. Sofela, Yekaterina Shulgina, Roma Nagle, Tianqi Wu, Hassane Mchaourab, Markus Voehler, Jamie H. D. Cate, Douglas A. Mitchell

## Abstract

Isoaspartate (isoAsp) formation is typically viewed as a “molecular clock” through nonenzymatic degradation of aspartate or asparagine during protein aging. Here we report a nearly universal enzymatic pathway for the formation of a conserved isoAsp in the bacterial ribosomal protein uS11. Proteome-wide protein-protein interaction scans using AlphaFold3 identified YbeY as a candidate enzyme from *Escherichia coli*. NMR spectroscopy supported a stable YbeY–uS11 complex from *Thermotoga maritima*. Biochemical assays indicated that *Ec*YbeY catalysis is zinc-dependent and prefers the conserved Asn-Gly motif for isoAsp formation. A high-resolution cryo-electron microscopy structure of the 70S ribosome from *E. coli* Δ*ybeY* revealed that loss of isoAsp alters contacts with the 16S rRNA groove and bS21. Phylogenetic analysis indicated that YbeY is present in almost all bacteria, and its absence is correlated to changes in the Asn-Gly motif of uS11. Additionally, our structural analyses implicate Fap7 as the functional counterpart in archaea and eukaryotes.

**Table of Contents:** 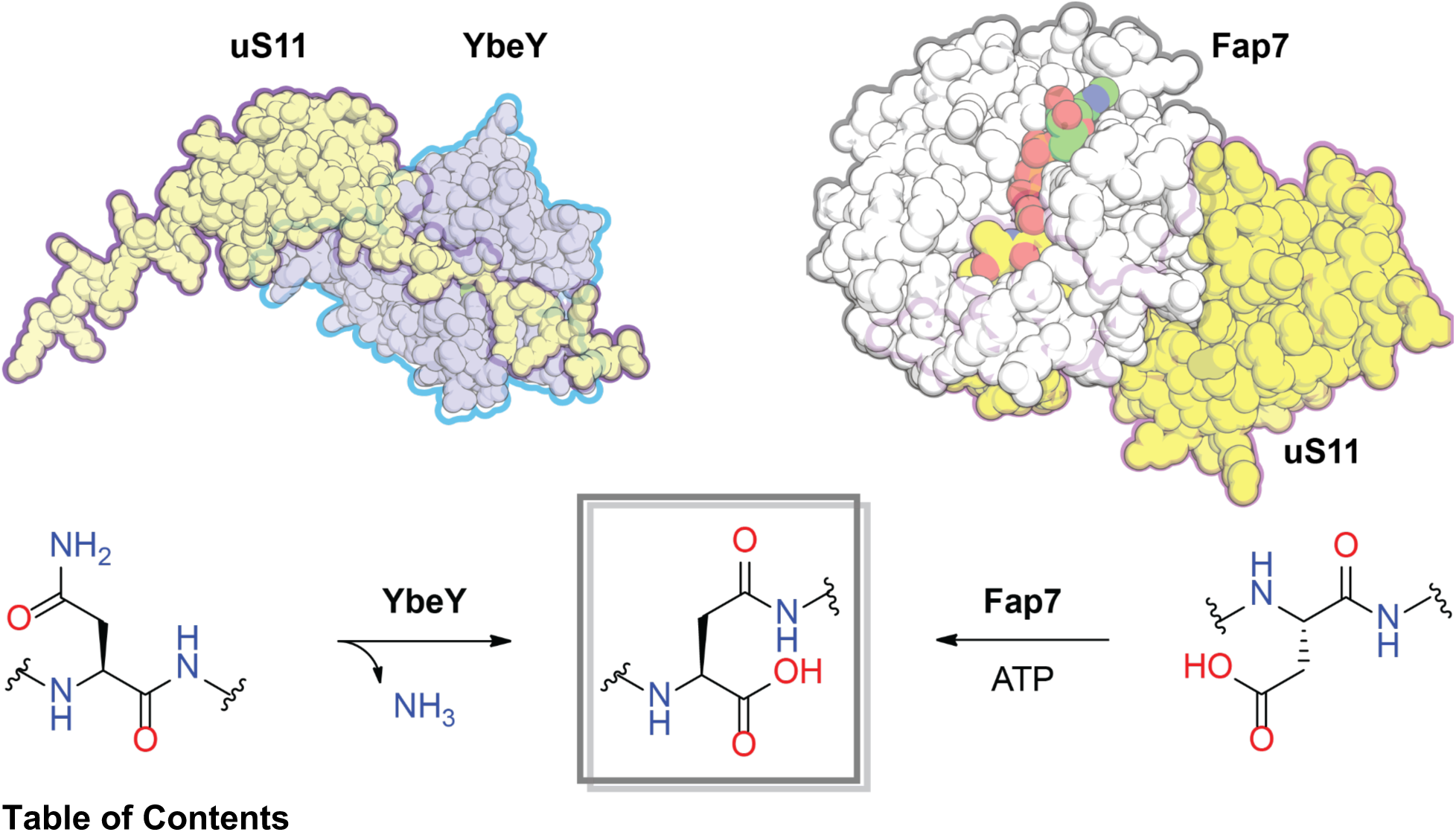

## Main

Formation of isoaspartate (isoAsp) within a polypeptide shifts the backbone from the α- to the β-carboxyl group of aspartate (Asp), thereby inserting an extra methylene group into the peptide backbone (Fig. 1a).^1^ IsoAsp is thought to form spontaneously at asparagine (Asn) or Asp residues in long-lived proteins, such as crystallins and Aβ,^2,3^ and has been linked to age-associated degenerative diseases.^4^ These spontaneous reactions are usually slow, occurring over days to years in model peptides,^5^ although a rapid example (t_1/2_ = 4.7 h) has been reported for MurA, a bacterial enzyme essential in peptidoglycan biosynthesis.^6^ IsoAsp can be reverted to Asp by protein L-isoaspartyl methyltransferases (PIMTs, InterPro: IPR000682), which act as intracellular repair enzymes.^7^ Enzymatic routes to isoAsp formation have also been reported at Asp residues through activation of the β-carboxyl group (Fig. 1a).^8–12^ For example, mammalian *O*-linked β-*N*-acetylglucosamine transferase (OGT) glycosylates an Asp residue in a non-native peptide and subsequently generates isoAsp via an aspartimide intermediate.^8^ Many ribosomally synthesized and post-translationally modified peptides (RiPPs) encode peptide/protein L-aspartyl *O*-methyltransferases (PAMTs), which are homologous to PIMTs, to methylate Asp residues and generate isoAsp products through the same aspartimide intermediate.^9,10^ In some cases, the aspartimide moiety is stable and constitutes the final product.^13–15^ The C-terminal Asn of cereblon substrates can also form a stable aspartimide product, catalyzed by a protein carboxymethyltransferase (PCMT1).^16^

**Figure 1.**
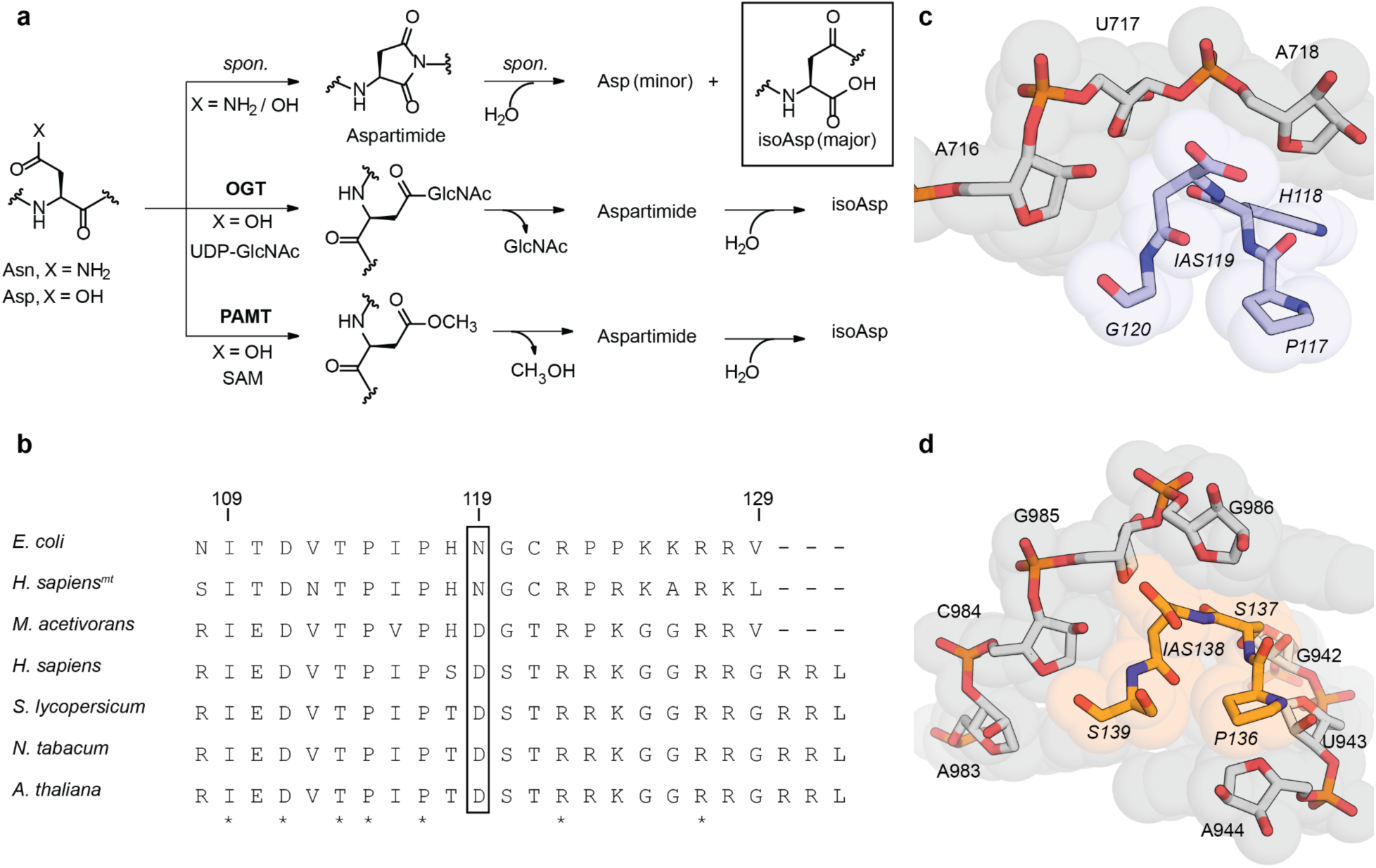
IsoAsp formation in ribosomal protein uS11. **a**, Biosynthetic pathways for isoAsp formation in the peptide backbone. OGT, *O*-linked β-*N*-acetylglucosamine transferase. PAMT, peptide/protein L-aspartyl *O*-methyltransferase. UDP, uridine diphosphate. GlcNAc, *N*-acetylglucosamine. SAM, *S-*adenosyl methionine. **b,** Multiple sequence alignment (MSA) of uS11 encoded in *E. coli* (PDB: 7K00), *Methanosarcina acetivorans* (PDB: 9ZNF), human *(Homo sapiens*) mitochondria (PDB: 7QI4) and nucleus (PDB: 8QOI), tomato (*Solanum lycopersicum*) (PDB: 7QIW), tobacco (*Nicotiana tabacum*) (PDB: 8B2L), and thale cress (*Arabidopsis thaliana*) (PDB: 9H3G). The C-terminal MSA is shown with numbering according to *E. coli* uS11. Columns containing an identical residue across all sequences are marked with asterisks. **c,** Space-filling model showing the shape complementarity between uS11 and 16S rRNA nucleotides near IAS119 in the *E. coli* 70S ribosome (PDB: 7K00). 16S rRNA is shown in gray and uS11 in blue, with residues labeled and uS11 residues italicized. IAS, isoAsp. **d,** Space-filling model showing the shape complementarity between uS11 and 18S rRNA nucleotides near IAS138 in the human 80S ribosome (PDB: 8QOI). 18S rRNA is shown in gray and uS11 in orange, with residues labeled and uS11 residues italicized.

Recent high-resolution cryo-electron microscopy (cryo-EM) analyses of ribosomes from our work and others have unambiguously revealed a stoichiometric isoAsp modification at a conserved Asn/Asp in uS11 across bacteria, archaea, and eukaryotes (Fig. 1b–d).^17–20^ In these structures, isoAsp consistently increases shape complementarity and stabilizes the local region at the interface with rRNA (Fig. 1c,d and Extended Fig. 1), but the mechanism of isoAsp formation was not examined. This presents a paradox in *Escherichia coli*, where uS11 (UniProt: P0A7R9) is near-stoichiometrically modified in early log phase despite a doubling time of ∼20 minutes,^21^ far shorter than the 1.2-day median half-life for spontaneous deamidation at an Asn-Gly motif.^5^ In contrast to MurA, where the isoAsp lies within a structurally constrained β-hairpin,^6^ uS11 isoAsp formation occurs at N119 that resides in the flexible C-terminal tail, which likely lacks the structural constraints needed to promote rapid spontaneous isomerization. Together, these observations suggest the involvement of an unknown enzyme. Previously, we used a co-folding method (AlphaFold3)^22^ to identify YcaO as the enzyme responsible for uL16 thioamidation.^23^ We reasoned that *bona fide* enzyme–substrate complexes can be distinguished from nonspecific predictions by higher confidence scores and by assessing whether the predicted complex is in a catalytically competent orientation.^23^ Here, we identified YbeY as the only promising candidate for enzyme-uS11 interaction using the proteome-wide scanning method. Phylogenetic analysis revealed that YbeY is present in most bacterial genomes, and that loss of YbeY co-occurs with a change to the isoAsp residue in uS11. Subsequent biophysical and biochemical assays validated YbeY binding to uS11 and YbeY catalytic activity. Mutagenesis assays further indicated that YbeY preferentially catalyzes isoAsp formation from Asn rather than Asp. In contrast, Asp-to-isoAsp conversion in uS11 likely proceeds via a distinct pathway involving Fap7, an ATPase essential in archaeal and eukaryotic ribosome biogenesis.^24,25^ Cryo-EM analysis shows the structural impact of the modified isoAsp or unmodified Asn in uS11 within the *E. coli* ribosome.

## Results

### *E. coli* proteome-wide AlphaFold3 scanning using *E. coli* uS11 as a query

Using locally installed AlphaFold3, *E. coli* uS11 (*Ec*uS11) was individually predicted in dimeric complexes with each of the 4,403 proteins in the *E. coli* K-12 proteome. Cofactors and ternary complexes were omitted from the analysis. During this screen, only 3 of 4,403 proteins (0.068%) were predicted to form a “confident” complex with *Ec*uS11, as defined by an interface predicted template modeling (ipTM) score > 0.8. These three proteins were: bS21 (UniProt: P68679), YchA (synonym SirB1; UniProt: P0AGM5), and YbeY (UniProt: P0A898) (Fig. 2a). To evaluate whether these predicted structures were catalytically competent for conversion of *Ec*uS11 N119 to isoAsp, we manually examined the structural models, focusing on the predicted local distance difference test (pLDDT) score at N119, the predicted aligned error (PAE) for N119 relative to the other protein, and the interactions of N119 within the predicted binding site. bS21 is a basic protein with an isoelectric point (pI) > 12,^26^ whereas YchA and YbeY are acidic, with predicted pI values of 4.5 and 4.2, respectively. Because *Ec*uS11 is highly basic (pI > 12),^26^ attractive electrostatic interactions should substantially contribute to complexation with YchA and YbeY. bS21 physically interacts with *Ec*uS11 in the 30S ribosomal subunit,^17^ and the predicted models likely recapitulate these interactions (Fig. S1). YchA is annotated as a transglutaminase-like/TPR repeat-containing protein and is upregulated during acid exposure.^27^ However, the relative position and orientation of *Ec*uS11 N119 with respect to YchA were not confidently predicted, based on low pLDDT value (73.5) and large PAE value (minimum PAE = 12.8 Å with YchA) (Fig. S2).

**Figure 2.**
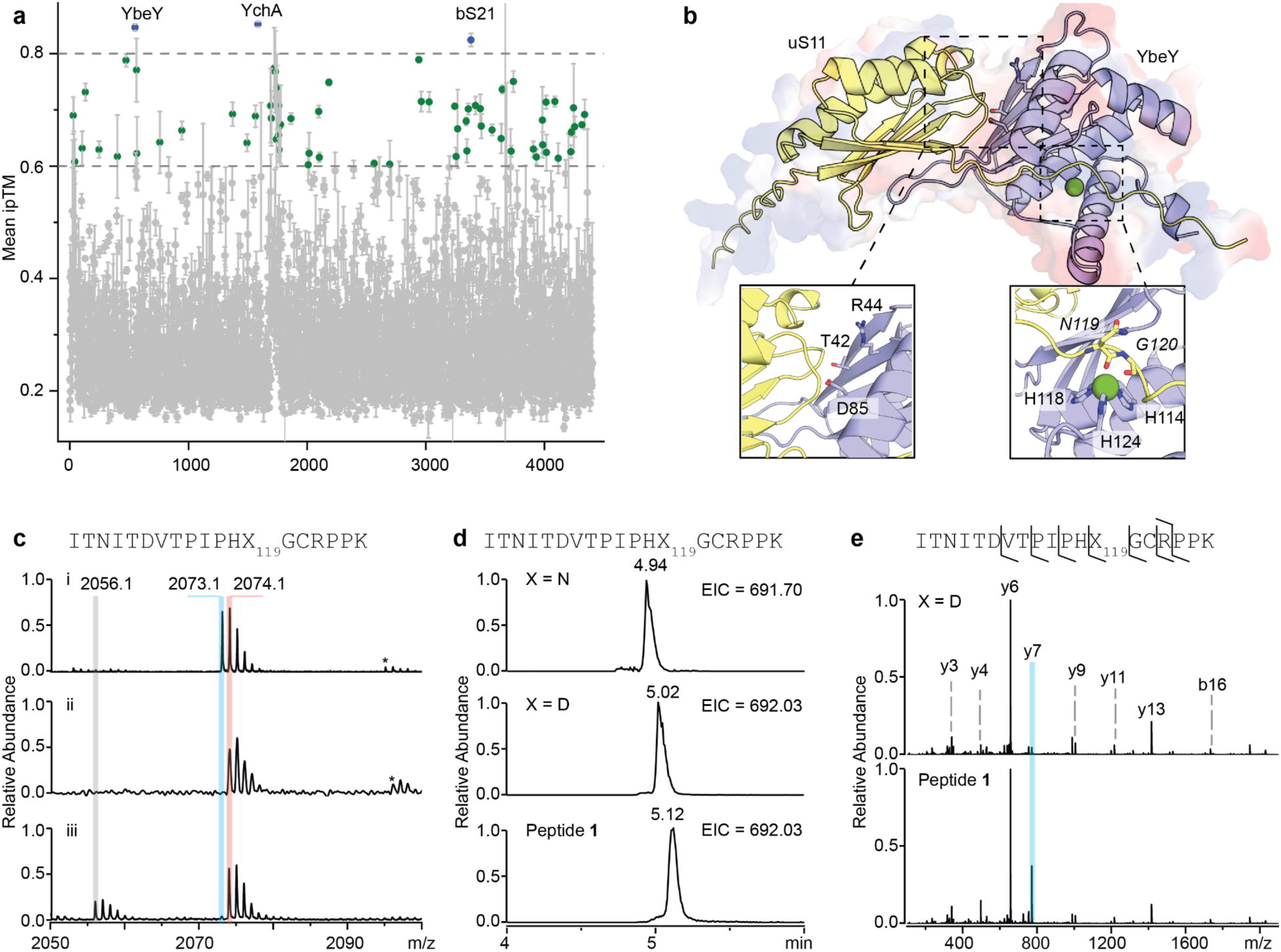
Bioinformatic and biochemical studies of isoAsp formation in *Ec*uS11. **a**, Table of mean ipTM score calculated from five models generated by AlphaFold3. Confident ipTM scores (>0.8) are shown as blue dots. Plausible iPTM values (>0.6) are shown as green dots. **b,** AlphaFold3 model of YbeY (blue) and uS11 (yellow) from *E. coli*. The electrostatic potential surface is shown in red (negative) and blue (positive). *Ec*YbeY residues important for *Ec*uS11 interaction (T42, R44, and D85) and zinc binding (H114, H118, and H124) are labeled. Zinc is shown as a green sphere. **c,** MALDI–TOF mass spectra of tryptic peptides derived from (i) *Ec*uS11 substrate, (ii) *Ec*uS11 + *Ec*YbeY, and (iii) *Ec*uS11 + *Ec*YbeY + EDTA. The sequence of peptide of interest is indicated. X = N, calculated [M+H]^+^ = 2073.0909; X = D or isoAsp (peptide **1**), calc. [M+H]^+^ = 2074.0749; and X = aspartimide (peptide **2**), calc. [M+H]^+^ = 2056.0644. Na^+^ adducts are labeled with asterisks. **d,** Extracted ion chromatograms of N119- or D119-containing peptide standards and peptide **1**. The sequence of peptide of interest is indicated. X = N, calc. [M+3H]^3+^ = 691.7018; X = D or isoAsp (peptide **1**), [M+3H]^3+^ = 692.0298. The retention time of each peptide is labeled on the corresponding peak. **e,** MALDI–TOF/TOF spectra of D119-containing peptide standard (upper) and peptide **1** (bottom). b/y ions are assigned, and the diagnostic y7 ion is highlighted.

By contrast, the *Ec*uS11 C-terminal tail was predicted to thread into a cavity of YbeY with higher confidence scores (pLDDT = 86.3; minimum PAE = 2.6 Å with YbeY) (Extended Fig. 2). YbeY homologs are essential for 16S rRNA maturation in *E. coli*, *Vibrio cholerae,* and *Sinorhizobium meliloti*.^28–30^ *E. coli* YbeY (*Ec*YbeY) has also been reported to interact with *Ec*uS11 and several ribosome-associated GTPases.^31^ The *Ec*YbeY residues previously implicated in *Ec*uS11 interaction (T42, R44, and D85) are located in a β-sheet predicted to participate in *Ec*uS11 binding.^31^ Although the *Ec*YbeY crystal structure (PDB: 1XM5) contained a nickel ion introduced during purification,^32^ subsequent studies have suggested that YbeY homologs are zinc-binding enzymes.^30,33^ We regenerated a three-component model by including a zinc ion, coordinated by three conserved histidines (H114, H118, and H124) in *Ec*YbeY and the N119 α-amide carbonyl of *Ec*uS11 (Fig. 2b). This geometry would likely facilitate deprotonation of the N119 α-amide nitrogen and subsequent nucleophilic attack on the N119 β-amide, making *Ec*YbeY a promising candidate enzyme for isoAsp formation. Although many other models showed plausible overall complex accuracy (ipTM 0.6–0.8) (Fig. 2a and Table S1), none provided confident predictions for the local interface around N119 in *Ec*uS11, and we therefore tentatively excluded these hits from further characterization.

### *Ec*YbeY catalyzes isoAsp formation in *Ec*uS11

To assess whether *Ec*YbeY directly catalyzes *Ec*uS11 isomerization, we produced *Ec*YbeY with an N-terminal maltose-binding protein (MBP) tag and *Ec*uS11 with an N-terminal His_6_ tag in *E. coli* BL21(DE3) and purified both proteins to homogeneity for *in vitro* studies (Fig. S3). After purification, *Ec*uS11 showed partial isoAsp formation at N119, as assessed by peptide mass fingerprinting after trypsin digestion and matrix-assisted laser desorption/ionization time-of-flight (MALDI–TOF) mass spectrometry (Fig. S4). We attribute this to endogenous *Ec*YbeY activity or spontaneous isomerization during protein expression. To evaluate the involvement of *Ec*YbeY, we generated an in-frame deletion of *ybeY* in *E. coli* K-12 (hereafter *E. coli* Δ*ybeY*) and used it as the host for *Ec*uS11 expression. *Ec*uS11 isolated from *E. coli* Δ*ybeY* lacked isoAsp modification (Fig. S5), suggesting a critical *in vivo* role for *Ec*YbeY. Of note, the *E. coli* Δ*ybeY* strain in the Keio collection^34^ is mislabeled: PCR indicates it retains wild-type (WT) *ybeY*.

In a 1 h *in vitro* endpoint assay, nearly all *Ec*uS11 substrate was converted to a +1 Da product (peptide **1**), while inclusion of EDTA yielded both peptide **1** and a -17 Da presumed intermediate (peptide **2**) (Fig. 2c). Peptide **1** was further analyzed by liquid chromatography–high-resolution MS (LC–HRMS) and HRMS/MS, which localized the modification to N119 (Fig. S6). The retention time (t*_R_*) of peptide **1** was distinct from those of N119- and D119-containing standards (Fig. 2d). To unequivocally identify the sequence of peptide **1**, we acquired MALDI–TOF/TOF spectra for the D119-containing standard and peptide **1**. Relative to the standard, fragmentation of peptide **1** yielded a diagnostic and intense y7 ion (Fig. 2e), consistent with preferential dissociation of the peptide bond immediately N-terminal to isoAsp (Scheme S1).^35,36^ In addition, peptide **1** resisted digestion by endoprotease AspN (Extended Fig. 3), which selectively cleaves the peptide bond immediately N-terminal to Asp but not isoAsp.^6,37^ Collectively, these data confirmed that peptide **1** contained IAS119. Peptide **2** was then analyzed by HRMS/MS and showed a 17 Da loss (-NH_3_) at N119 (Fig. S7), consistent with formation of an aspartimide intermediate commonly observed during isoAsp formation (Fig. 1a).^8–10^ Given that EDTA partially inhibited isoAsp formation, we proposed that *Ec*YbeY promotes the reaction in a metal-dependent manner.

### Interactions in the *Thermotoga maritima* YbeY–uS11 complex in solution

A solution NMR structure is available for the YbeY homolog from *Thermotoga maritima* (*Tm*YbeY; 27% identical to *Ec*YbeY; PDB: 1TVI),^38^ so we used the *T. maritima* proteins to probe YbeY–uS11 interactions. Genes encoding *Tm*YbeY and *T. maritima* uS11 (*Tm*uS11; UniProt: Q9X1I4; 55% identical to *Ec*uS11) were codon-optimized and cloned for individual expression in BL21(DE3). After purification, *Tm*uS11 showed partial modification (Fig. S8), likely from endogenous *Ec*YbeY processing in the host strain. *In vitro* reconstitution confirmed that *Tm*YbeY catalyzes isoAsp formation on *Tm*uS11. To investigate the binding mode, we first generated a structural model of the *Tm*YbeY–*Tm*uS11–zinc complex (Fig. 3a–c). *Tm*YbeY retains the HX_3_HX_5_H zinc-binding motif but differs from *Ec*YbeY at several positions implicated in substrate recognition (Fig. S8). For example, T42, R44, and D85 in *Ec*YbeY correspond to N35, I37, and E73 in *Tm*YbeY, respectively. The model further suggested contacts between *Tm*YbeY and the *Tm*uS11 C terminus: T58, D59, and N84 engage the N120-G121 motif, whereas E87, E95, E98, and N90 contact other C-terminal basic residues.

**Figure 3.**
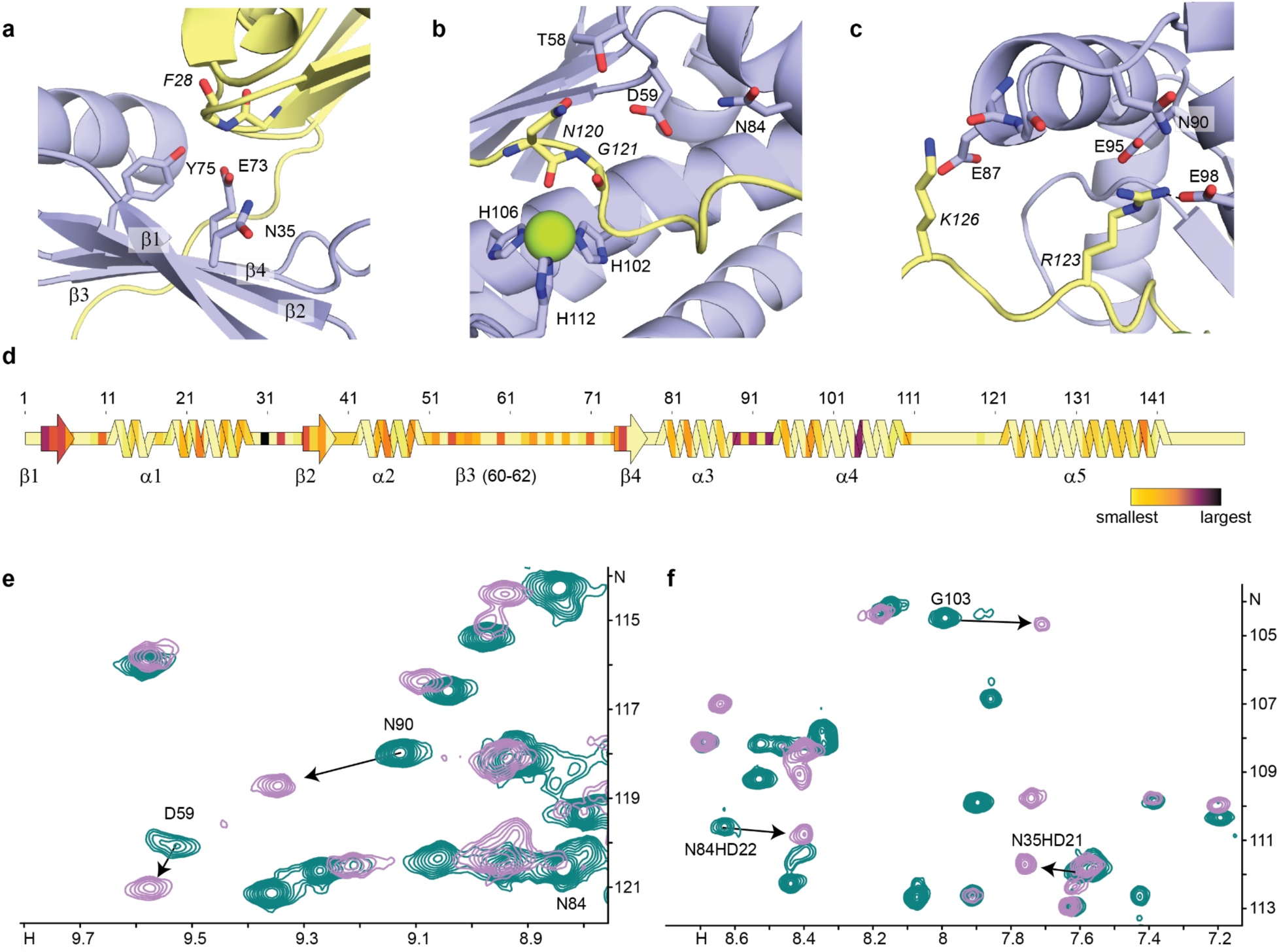
Structural analysis of the YbeY-uS11 complex in *T. maritima*. **a**–**c,** Zoomed regions of the AlphaFold3 model for *Tm*YbeY-*Tm*uS11. *Tm*YbeY and *Tm*uS11 are shown in blue and yellow, respectively. Residues in *Tm*uS11 are italicized. Zinc is shown as a green sphere. **d,** Secondary structure diagram of *Tm*YbeY colored by chemical shift perturbation (CSP) values. Residues not observed in the HSQC spectra were shown a CSP value of 0. Residue numbering (top) and secondary structure element (bottom) are indicated. The short β3 strand is indicated the same as the original report.^38^ **e** and **f,** Zoomed overlays of ^1^H–^15^N HSQC spectra of *Tm*YbeY in the apo form (teal) and in the presence of one equivalent of unlabeled *Tm*uS11 (purple). Apo peaks of interest are assigned, while the corresponding homo peaks are indicated by arrows.

We then used ^1^H–^15^N heteronuclear single quantum coherence (HSQC) nuclear magnetic resonance (NMR) spectroscopy to monitor backbone amide chemical shift perturbations (CSPs) upon binding. Uniformly ^15^N-labeled *Tm*YbeY and unlabeled *Tm*uS11 were prepared, and ^1^H-^15^N HSQC spectra were recorded during titration of *Tm*uS11 at 55 °C to alleviate precipitation. Excluding five prolines (undetectable in ^1^H–^15^N HSQC), 118 of 145 backbone amides (81%) were observed in apo *Tm*YbeY (Fig. S9), along with several well-resolved Asn β- and Gln γ-amides. Unresolved peaks mainly clustered in the loop connecting α4–α5 and the zinc-binding pocket, implying rapid exchange with bulk water under the conditions used. During titration with *Tm*uS11, many amide peaks decreased in intensity with the concomitant appearance of new peaks, while few additional peaks changed beyond a 1:1 *Tm*YbeY:*Tm*uS11 molar ratio (Extended Fig. 4), indicating formation of a stable complex that exchanges slowly with free subunits on the NMR timescale.^39^

We assigned the bound spectra using a minimum chemical-shift approach.^39^ The C-terminal segment of *Tm*YbeY was least perturbed (Extended Fig. 4), consistent with its distance from the predicted interface. In line with the structural model, the *Tm*YbeY β-sheet likely mediates multiple contacts with *Tm*uS11, supported by widespread perturbations across all four β-strands (Fig. 3d). Although the E73 (in β4) backbone amide was not observed, the N35 (in β2) backbone amide exhibited a large CSP, also clearly reflected in the N35HD21 peak (Fig. 3f). Residues surrounding the zinc-binding pocket were likewise sensitive to binding: D59, N84, and G103 exhibited pronounced CSPs, potentially reflecting accommodation of the *Tm*uS11 N120-G121 motif (Fig. 3e and 3f). Large CSPs were also detected in several loops, including the loop containing N90, suggesting engagement of the *Tm*uS11 C-terminus for substrate recognition and/or additional conformational changes upon binding.

### Mutational and mechanistic analysis of *Ec*YbeY

To probe biological function, ten residues of *Ec*YbeY were selected for mutational analysis based on the *Ec*YbeY–*Ec*uS11 model, NMR data of *Tm*YbeY–*Tm*uS11, and previous functional studies of *Ec*YbeY.^28,31,40^ Among ten variants tested for isoAsp formation (Table 1 and Fig. S10), conversion rates in endpoint assays ranged from ∼25% to >95%. Of the three residues (H114, H118, and H124) in *Ec*YbeY that coordinate the metal ion,^32^ only the *Ec*YbeY H114A variant caused a pronounced activity defect (∼50%). To estimate zinc-binding affinities for *Ec*YbeY WT and variants, we performed isothermal titration calorimetry (ITC) experiments using freshly prepared zinc solution and EDTA-treated proteins. *Ec*YbeY WT bound zinc with a dissociation constant (K*_D_*) of 2.3 μM (Fig. S11), similar to the YbeY homolog from *Staphylococcus aureus* (*Sa*YbeY, K*_D_* = 1.7 μM).^33^ The *Ec*YbeY H118A and *Ec*YbeY H124A variants showed modestly weaker affinities (K*_D_* = 5.4 μM and 5.8 μM, respectively), whereas the *Ec*YbeY H114A showed substantially reduced affinity (K*_D_* = 16.8 μM). Taken together with the activity results, these data support a zinc-dependent mechanism for *Ec*YbeY catalysis.

**Table 1.** Activities of *Ec*YbeY variants and modification on *Ec*uS11 variants.

| *EcYbeY* variant | Conversion ratio<sup>a</sup> | *EcuS11* variant | Conversion ratio<sup>a</sup> |
| --- | --- | --- | --- |
| WT | >95% | WT | >95% |
| T42F | ~85% | P117A | ~80% |
| R44D | ~75% | H118A | ~90% |
| N55A | ~95% | N119D | N.D. |
| R59A | >95% | N119E | N.D. |
| T65A | ~75% | N119Q | ~15% |
| N66A | ~70% | N119A | N.A. |
| D85R | ~25% | G120A | <5% |
| H114A | ~50% | C121A | >95% |
| H118A | ~90% | R122A | ~90% |
| H124A | ~90% | C-ter peptide<sup>b</sup> | N.D. |

<sup>a</sup>The ratio was estimated based on areas in LC–HRMS spectra (Fig. S10 and S15). Values are approximated to the nearest 5-unit interval. <sup>b</sup>The C-terminal peptide (residues 107-125) was analyzed by MALDI–TOF. N.D., not detected. N.A., not available.

Residues T42, R44, and D85 in *Ec*YbeY have been implicated in *Ec*uS11 binding *in vivo*,^31^ suggesting that the reduced activity of these variants reflects weakened substrate binding. *Ec*YbeY D85R showed substantially diminished activity (∼25%). The R59 residue has been proposed to be important for the RNase activity of YbeY,^28,30^ yet the *Ec*YbeY R59A and *Ec*YbeY N55A variants retained WT-level activity (≥ 95%), implying that isoAsp formation uses a distinct set of residues from those required for the putative RNA hydrolysis activity. In the structural model, T65 and N66 are predicted to form hydrogen bonds with the *Ec*uS11 N119 α- and β-amides (Fig. S12). The modest activity decreases observed for *Ec*YbeY T65A (∼75%) and N66A (∼70%) variants are likely due to disruption of these local interactions with the substrate.

### Fap7 is predicted for isoAsp formation in eukaryotes and archaea

The YbeY protein family (InterPro: IPR002036; ∼20,000 proteins as of July 2026) comprises sequences from bacteria (∼90%) and eukaryotic organelles (∼10%).^41^ The two eukaryotic YbeY homologs from thale cress (*At*YbeY) and human (*Hs*YbeY) were reported to localize to chloroplasts and mitochondria, respectively.^42–44^ We generated structural models for *At*YbeY (UniProt: Q8L5Z4) with *At*uS11c (UniProt: P56802) and *Hs*YbeY (UniProt: P58557) with *Hs*uS11m (UniProt: P82912), with zinc included in each model (Extended Fig. 5). Both models resemble the *E. coli* YbeY-uS11 complex, particularly in zinc coordination and recognition of the uS11 Asn substrate, supporting a conserved catalytic role for eukaryotic YbeY homologs. *At*YbeY also has an additional haloacid dehalogenase hydrolase-like domain conserved among plant-derived YbeY homologs,^42^ although it does not participate in *At*uS11 binding in the model.

We then investigated how isoAsp forms on uS11 in the eukaryotic nucleus and in archaea. Using a predicted high-accuracy human interactome,^45^ we evaluated all putative binders for human cytoplasmic uS11 (*Hs*uS11, UniProt: P62263) and identified a single strong candidate, adenylate kinase isoenzyme 6 (*Hs*AK6, UniProt: Q9Y3D8), that formed a catalytically plausible complex. *Hs*AK6 is a cell nuclear-localized ATPase whose activity is activated by *Hs*uS11 binding.^46,47^ We generated a structural model of *Hs*AK6–*Hs*uS11–ATP in high confidence (ipTM = 0.9) (Extended Fig. 6). In the model, the C-terminal extension of *Hs*uS11 extended into the ATP-binding pocket of *Hs*AK6. In particular, the ATP γ-phosphate was positioned very close to the *Hs*uS11 D138 β-carboxylate. This structural arrangement is reminiscent of the catalysis of ATP-grasp ligase, in which ATP is used to activate the acceptor Asp or Glu sidechain carboxylate.^48^ The AK6 homolog in yeast and archaea is factor activating pos9 (Fap7), whose ATPase activity also depends on uS11 binding.^24,49^ A co-crystal structure of Fap7–uS11 was determined for the archaeon *Pyrococcus abyssi*, in which *Pa*uS11 was reported with an unmodified D124.^24^ By carefully fitting the electron density, we refined the structure to reveal that *Pa*uS11 in this structure actually contains isoAsp at residue 124 (IAS124, Extended Fig. 7). The transformation likely occurred prior to crystallization, as the protein complex was co-expressed using a polycistronic construct. We therefore propose that Fap7 is the enzyme responsible for isoAsp formation in archaea and eukaryotes by activating the β-carboxylate of Asp.

### Phylogenetic distribution of YbeY is correlated with the NG motif in uS11

To investigate the phylogenetic distribution of YbeY and motifs in uS11, we mined the representative genomes of all 189,801 bacterial species in the Genome Taxonomy Database (GTDB) for homologs of uS11, YbeY, and Fap7 (Supplementary Files 1–4). uS11 and YbeY are nearly ubiquitous, with homologs found in 89% and 92% of bacterial genomes, respectively. Genes may be missing from a genome due to the incomplete assembly or bona fide absence. We examined whether the presence of a YbeY homolog is associated with the canonical amino acid at modified position (N119, using *E. coli* numbering to describe motif positions) and also at surrounding residues 117, 118, and 120. Across bacteria, the presence of a YbeY homolog co-occurs with N119 and G120, while deviations from the canonical P117 or H118 are not strongly associated with YbeY absence (Fig. 4a). uS11 motifs lacking the canonical N119 and G120 are accompanied by a sharp drop in YbeY occurrence (down to 15.5% and 33.3%, respectively), suggesting that bacterial lineages that have lost YbeY, and by extension, formation of isoAsp, have evolved alternative sequences in this region of uS11. Focusing on the most common alternative motifs at positions 119 and 120 reveals that the effect is amino acid specific: some alternative motifs, such as D119–G120, retain high presence of YbeY, suggesting that D119 may serve as a substrate for YbeY in some species, while others, most notably G119–G120, predominantly occur in genomes that lack a detectable YbeY homolog (Fig. 4b).

**Figure 4.**
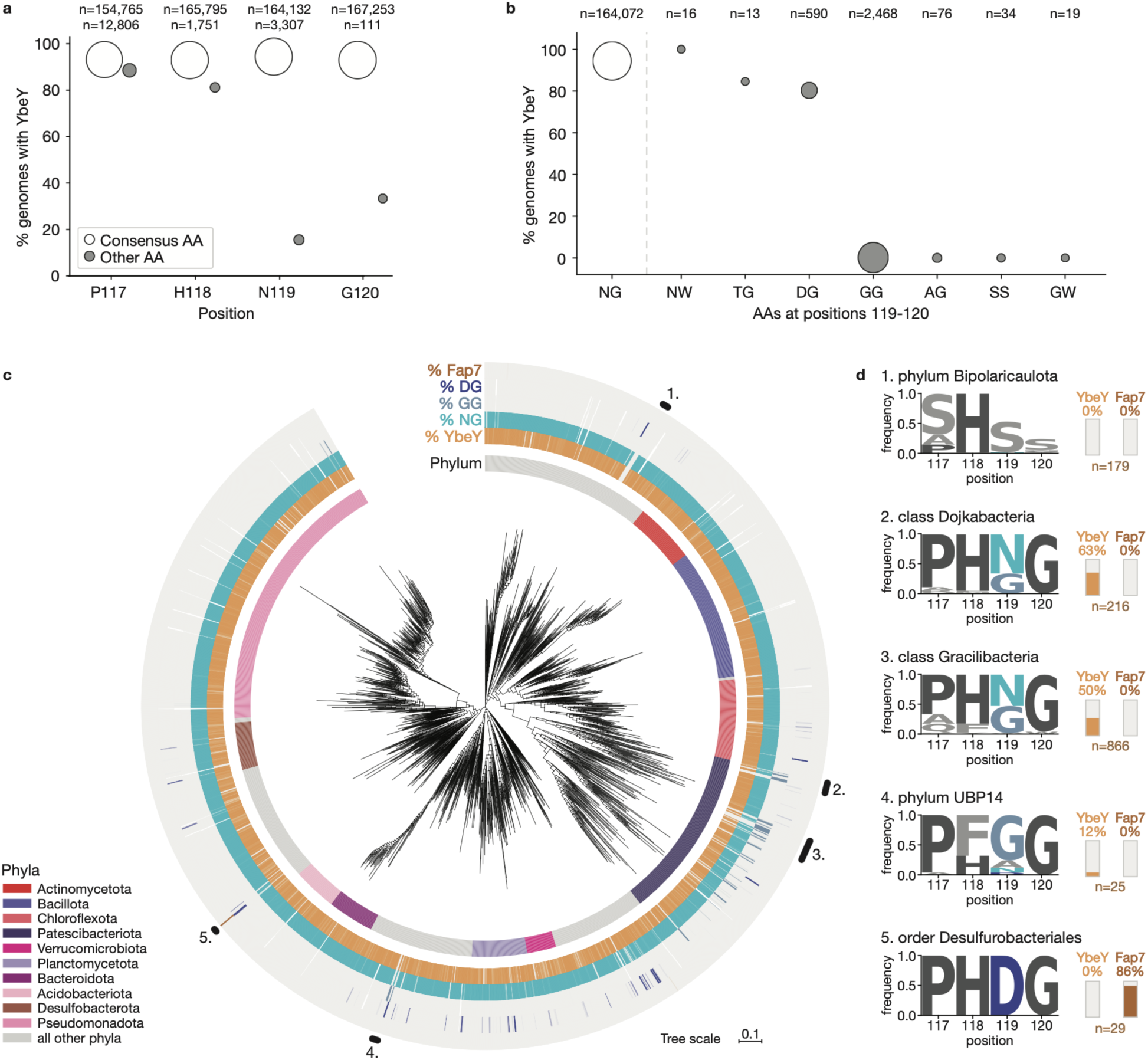
Phylogenetic distribution and co-occurrence of uS11 motifs with YbeY and Fap7 gene presence. **a**, For each position in the uS11 motif (*E. coli* positions 117-120), the percentage of bacterial genomes encoding a YbeY homolog is reported, separately for the canonical amino acid (open circles) or any other amino acid (filled circles). Circle area is proportional to the log of the number of genomes (n, shown above each pair of circles). Genomes with a gap or ambiguous residue at the corresponding alignment position were excluded from the counts. **b,** For combinations of amino acids at positions 119-120 seen in at least 10 genomes, the percentage of bacterial genomes encoding a YbeY homolog, ordered by descending YbeY occurrence. Circle area is proportional to the log of the number of genomes (shown above each circle). **c,** Phylogenetic distribution of YbeY and Fap7 homologs and uS11 motifs at positions 119-120 across GTDB bacteria. The GTDB bacterial phylogenetic tree is shown with each node collapsed to the order level, with the ten largest phyla indicated in the inner ring. Outer rings show presence of YbeY and Fap7 homologs as percent of species per order, and presence of uS11 motifs at position 119-120 NG, GG, and DG as percent of uS11 homologs from species in the order (white indicates orders with no annotated uS11 homologs). Clades of interest that are missing YbeY are marked on the tree and in **d,** show further information, such as residue frequency (including gaps) at uS11 motif positions 117-120 and percent of species in the clade that have YbeY and Fap7 homologs.

Visualizing the presence of YbeY homologs on the GTDB bacterial phylogenetic tree alongside usage of uS11 motifs reveals scattered clades that have lost YbeY (Fig. 4c). Multiple clades have evolved a concerted change from N119 to G119 (Supplementary File 1), including members of the classes Dojkabacteria and Gracilibacteria (phylum Patescibacteriota) and the phylum UBP14 (Fig. 4d). We also observed an unusual change in the phylum Bipolarcaulota (Fig. 4d), where loss of YbeY occurs alongside a truncation of uS11 to end after position 118. To confirm if this reflects a genuine coding change rather than an annotation artifact or frameshifting event, we translated the genomic uS11 window of Bipolaricaulota in all three reading frames (Fig. S13). Frame 0 retains H118 but carries divergent residues at positions 117 and 119 and a stop codon at the position 120, and other reading frames lack conserved sequence after the stop codon. Thus, the disrupted motif in Bipolaricaulota likely represents a truncated uS11 that lacks N119, consistent with loss of YbeY. While Fap7 was absent from almost all bacterial genome (Fig. 4c), homologs were found in 86% of genomes in the order Desulfurobacteriales. These may have been acquired by horizontal gene transfer, accompanied by a concurrent change of the uS11 motif to D119-G120, the typical motif for Fap7 (Fig. 4d, Extended Fig. 7d).

### *Ec*YbeY prefers the NG motif in *Ec*uS11

To experimentally probe the correlation of YbeY and the NG motif of uS11, we conducted a mutagenesis study on *Ec*uS11. We first tested if *Ec*YbeY can process the C-ter peptide (residues 107–125) alone instead of the intact *Ec*uS11 protein. Under the same conditions, modification (+1 Da shift) was not detected for the C-ter peptide (Fig. S14), suggesting that the globular domain of *Ec*uS11 is required for efficient turnover. Next, the N119 and neighboring residues in *Ec*uS11 were selected for mutational analysis (n = 9) (Table 1). Eight variants were purified and used as substrates for *in vitro* assays (Fig. S3), except *Ec*uS11 N119A, for which we were unable to obtain enough soluble protein. A +1 Da mass shift is expected when a sidechain amide (Asn or Gln) is converted to a carboxylate. LC–HRMS spectra showed that the +1 Da product predominated in reactions using substrates including *Ec*uS11 P117A (∼80%), *Ec*uS11 H118A (∼90%), *Ec*uS11 C121A (>95%), and *Ec*uS11 R122A (∼90%) (Fig. S15). The corresponding tryptic peptides **3**, **4**, **5**, and **6** were assigned as isoAsp-containing products based on HRMS/MS, AspN digestion, and the high-intensity y7 ion in MALDI–TOF/TOF spectra (Fig. S16-19). The efficient conversion of *Ec*uS11 C121A also argues against an alternative cysteine-engaged mechanism for isoAsp formation (Scheme S2). In contrast, only trace (<5%) +1 Da product (peptide **7**) was observed for the reaction with *Ec*uS11 G120A substrate. Peptide **7** was analogously assigned as an isoAsp-containing product following the same procedures (Fig. S20). For *Ec*uS11 N119Q, we observed a minor amount (∼15%) of +1 Da product (peptide **8**) with the modification localized to Q119 by HRMS/MS (Fig. S21). In MALDI-TOF/TOF spectrum, peptide **8** produced a high-intensity y7 ion (Fig. S22), supporting the presence of an α-E119 residue (Scheme S1). We tentatively assigned this as an isoglutamate (isoGlu)-containing product, potentially formed through a six-membered glutarimide intermediate (Scheme S3). For *Ec*uS11 N119D and *Ec*uS11 N119E, any analogous isomerization would be isobaric but should shift the t*_R_* due to structural rearrangements. However, no new peaks were observed after reaction with *Ec*YbeY (Fig. S15). The corresponding peptides **9** and **10** eluted at t*_R_* values distinct from those of the putative isomerized peptides (**1** and **8**, respectively). In addition, the y7 ion was markedly reduced in the MALDI-TOF/TOF spectra of peptides **9** and **10** (Fig. S23 and S22, respectively). AspN digestion of peptide **9** yielded peptide fragments distinct from those of peptide **1**, and cleavage between residues 118 and 119 was detected (Fig. S23). Collectively, these results indicate that while *Ec*uS11 N119Q is tolerated by *Ec*YbeY, the *Ec*uS11 N119D and N119E variants are not.

### Ribosomes from *E. coli* Δ*ybeY* show defects in association and *in vitro* translation activity

To evaluate the impact of the isoAsp modification on ribosome function, we tested the ability of ribosomal subunits isolated from *E. coli* Δ*ybeY* (ΔY-ribosomes) to associate into 70S ribosomes using analytical 20-50% sucrose gradients. First, we equilibrated purified 30S and 50S subunits from the parental strain (*E. coli* MRE600, used as the WT strain for ribosome assembly assays) or *E. coli* Δ*ybeY* (Fig. 5a,b), in a 2:1 molar ratio in the presence of 10 mM MgCl_2_ to visualize subunit association (Fig. 5c,d). As expected, the WT subunits were completely associated (Fig. 5c). However, subunits from the *ΔybeY* strain showed a significant association defect, with peaks corresponding to unassociated 50S subunits observed (Fig. 5d). To test whether this association defect resulted from unfolding that may have taken place while 30S subunits were isolated under low (1 mM) MgCl_2_ conditions, we loaded crude 70S ΔY-ribosomes directly onto a 20-50% sucrose gradient supplemented with 15 mM MgCl_2_ and observed significant quantities of unassociated 30S and 50S subunits (Extended Fig. 8a). These results indicated an inherent association defect in these ribosomes, perhaps due to deficiency in subunit assembly or incomplete rRNA processing in the absence of YbeY.^28,40^ We next tested whether the observed association defect correlated with the accumulation of unprocessed 16S rRNA. Using agarose gel electrophoresis, we visualized rRNA isolated from the peaks corresponding to 30S, 50S and 70S from *E. coli* Δ*ybeY* and WT (Extended Fig. 10i). Comparison of the bands showed the presence of 17S rRNA in the unassociated 30S sample from *E. coli* Δ*ybeY*, but only completely processed 16S rRNA in the associated 70S. These results indicated that the assembly defect may stem from the accumulation of unprocessed 17S rRNA in *E. coli* Δ*ybeY*.^28,40^ Next, we tested the activity of these ribosomes by supplementing *in vitro* translation reactions with associated WT or ΔY-ribosomes and measuring the luminescence output of an encoded nanoluciferase reporter. We found that despite being associated into 70S, ΔY-ribosomes only exhibited ∼30% of WT activity, indicating compromised translation activity beyond a simple association defect (Fig. 5e). Lastly, we measured growth curves of the WT and the Δ*ybeY* mutant in rich medium (Fig. 5f). The *ΔybeY* mutant exhibited a reduced growth rate, consistent with previous reports.^40^ This growth defect was largely rescued by expressing plasmid-encoded WT YbeY and the R59A YbeY variant that was previously shown to lack RNAse activity,^28^ but not by the D85R or H114A variants. Together, these results link isoAsp formation in *Ec*uS11 to growth rate, which is proportional to the abundance of active ribosomes *in vivo*.^50^

**Figure 5.**
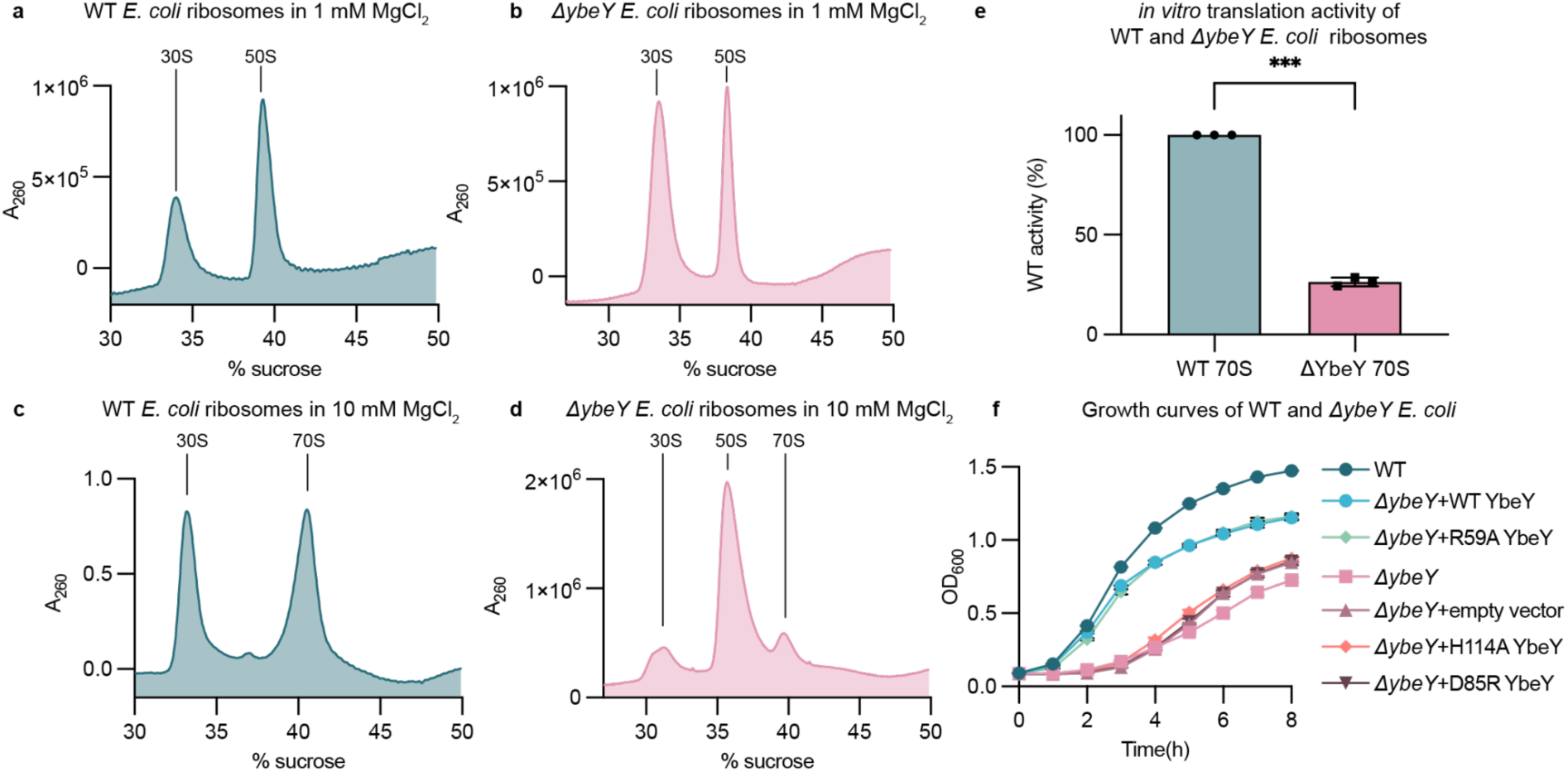
Ribosomes from *E. coli* Δ*ybeY* show a defect in association and activity. Sucrose gradient profiles of ribosomes isolated from WT and *E. coli* Δ*ybeY* under **a,b** low (1 mM) or **c,d** high (10 mM) Mg^2+^ conditions. **e,** *In vitro* translation activity of associated 70S ΔY-ribosomes compared to WT ribosomes. Activity is defined as the slope of the increase in luminescence signal observed upon translation of nanoluciferase. Activities were normalized to WT and data and error bars represent the mean and standard deviation of the three measurements, p-values were calculated to be 0.0003. **f,** Growth behavior analysis. Datapoints and error bars represent to the mean and standard deviation of three biological replicates.

### Unmodified uS11 results in minimal structural changes to 16S rRNA in associated 70S particles

MALDI–TOF MS analysis revealed that *Ec*uS11 in 70S ribosomes isolated from *E. coli* WT contains stoichiometric levels of IAS119, whereas *Ec*uS11 from the Δ*ybeY* mutant retains unmodified N119 (Fig. S24). To visualize the structural impact of having unmodified N119 in uS11, we obtained cryo-EM structures of associated 70S ΔY-ribosomes. Initial rounds of 2D and 3D classification were performed to select intact 70S ribosomal particles (Extended Fig. 8). The resulting cryo-EM maps contained complexes in the classical (non-rotated) state of the ribosome. To improve the resolution of uS11, we performed local refinement using a 30S subunit mask, resulting in a reconstruction with 1.91 Å global resolution (Extended Fig. 9). This resolution allowed for unambiguous modeling of the P117-G120 region of uS11 with clearly resolved density for unmodified N119 (Fig. 6, Extended Fig. 9). Remarkably, while previous structures have shown that the additional methylene group introduced by the isoAsp modification results in high shape complementarity between uS11 and the proximal 16S rRNA nucleotides (Fig. 6e-g), the present structure shows that unmodified N119 in uS11 also maintains this shape complementarity and results in tight packing with the groove formed by rRNA nucleotides 716-718 and stacking of H118 in uS11 with the conserved purine nucleotide A718 (Fig. 6h-j).

**Figure 6.**
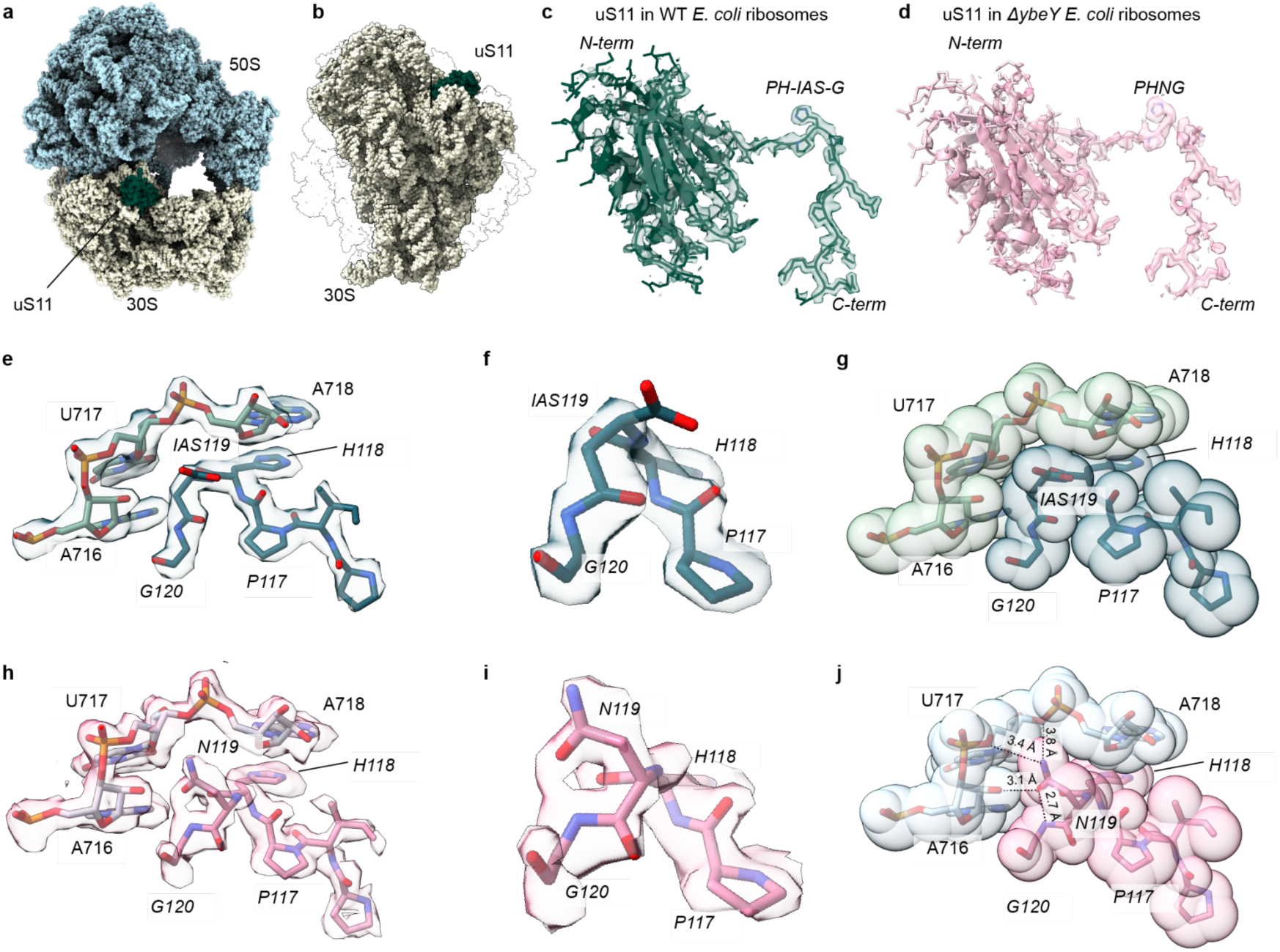
Unmodified N119 in *Ec*uS11 results in minimal local or global structural changes in the ribosome. **a**, side view of the *E. coli* ribosome (PDB ID 7K00) with the 50S subunit indicated in light blue and the 30S subunit indicated in light yellow. 30S ribosomal protein uS11 is shown in dark green. **b,** view from the intersubunit interface of the 30S subunit showing the location of uS11 (dark green) in the platform region. The 50S subunit is indicated in a transparent outline. **c,** Atomic structure and cryo-EM density of uS11 in WT *E. coli* ribosomes (green) and **d,** ΔY-ribosomes (pink) showing overall fold, protein domains, and PHNG motif. **e-g,** zoomed in view of cryo-EM density and space-filling model showing the accommodation of the PHNG loop within a rRNA groove created by nucleotides 716-718 (light green). **h-j,** atomic structure and cryo-EM density of the PHNG loop of uS11 (pink) and 16S rRNA (light blue) in ΔY-ribosomes.

Further inspection of the ΔY-ribosome structure and comparison to WT ribosomes, shows that the β-amide in unmodified N119 makes hydrogen bonds with the sugar-phosphate backbone of the 16S rRNA, and also hydrogen bonds to the G120 α-nitrogen (Fig. 6g,j). In contrast, in WT 30S subunits, the α-carboxylate of IAS119 is turned away and the peptide backbone primarily makes van der Waals contacts with 16S rRNA.

### Impact of loss of isoAsp on uS11 interactions with ribosomal protein bS21

Ribosomal protein bS21 physically interacts with the C-terminal region of uS11 within the WT 30S platform.^17^ Due to structural disorder in this region, we applied a low-pass filter of 3.5 Å to the 30S focus refined map from WT and ΔY-ribosomes to clarify the density of bS21. In the WT ribosome, the filtered map has clear density for the entire length of bS21 (Extended Fig. 10a). bS21 R35 is positioned proximal to uS11 IAS119 and can form H-bonding contacts with the IAS119 α-carboxylate and backbone amide carbonyl group of uS11 P117 (Extended Fig. 10b). In ΔY-ribosomes, this interaction is disrupted due to sidechain rotation of uS11 N119 away from bS21 (Extended Fig. 10e). In addition, bS21 has lower occupancy and poor density in ΔY-ribosomes, especially in the C-terminal region, where bS21 Y71 of the conserved RLY motif interacts with rRNA nucleotide A1167 in helix 40 of the 30S head domain (Extended Fig. 10c,d,f).^17^ Low-pass filtering of the maps reveals density for the Shine-Dalgarno helix at the 3′-end of the 16S rRNA from WT ribosomes, but not in ΔY-ribosomes, although this observation may be attributed to addition of exogenous mRNA and tRNA in the WT complex (Extended Fig. 10c,f). Lastly, reconstruction from particles classified on bS21 occupancy improves the cryo-EM density for this protein, but the interaction between uS11 N119 and bS21 R35 remains disrupted (Extended Fig. 10g,h). Collectively, the cryo-EM structure indicates that in associated 70S particles, the lack of the isoAsp modification results in minimal local structural changes within the overall fold of uS11 and shape complementarity within the 16S rRNA binding pocket but affects the accommodation of protein bS21 in the 30S subunit.

**Scheme 1.**
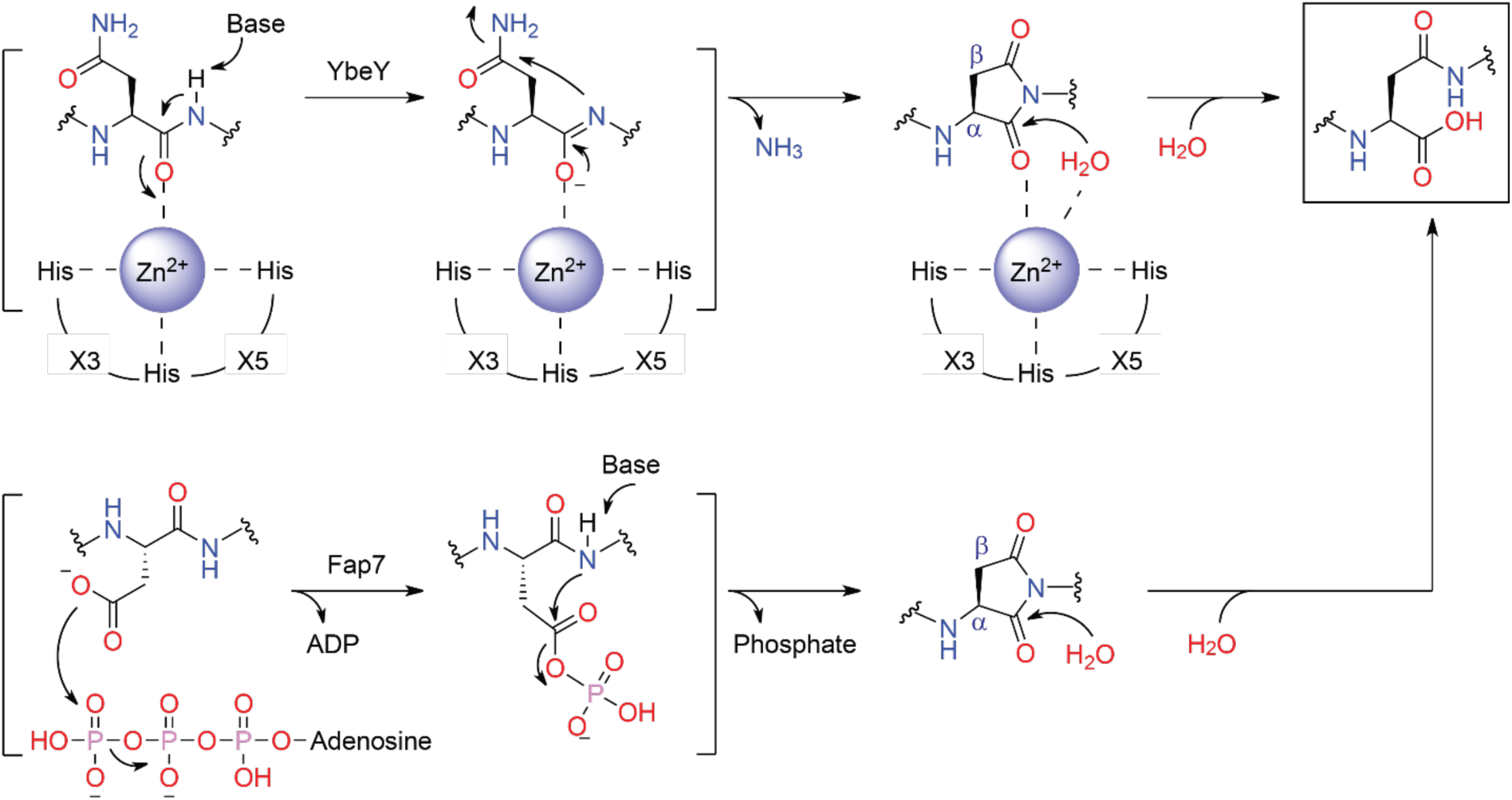
Proposed reactions for isoAsp formation. Possible reactions catalyzed by YbeY (upper) and Fap7 (bottom) are shown. The HX_3_HX_5_H motif for zinc binding is indicated for the catalysis by YbeY.

## Discussion

We have identified YbeY as a nearly universal enzyme in bacteria that catalyzes isoAsp formation at a conserved Asn residue in bacterial uS11. Although the Asn–Gly motif in the peptide backbone can spontaneously form an isoAsp residue over days,^5^ YbeY accelerates this conversion to match the minute timescale of ribosome assembly *in vivo*. YbeY family proteins have a conserved HX_3_HX_5_H motif for zinc binding,^32,33,38^ likely leaving the fourth zinc coordination site occupied by water in the apo protein and replaced by a uS11 backbone amide in the substrate-bound complex. As a Lewis acid, coordinating to zinc will polarize the backbone amide, making the N-H more acidic and the nitrogen more nucleophilic for subsequent reaction with the side chain amide (Scheme 1). Once the putative aspartimide intermediate is formed, hydrolysis appears to be regioselective for the α-position because we did not detect D119-containing product after reaction with YbeY. The regioselectivity likely arises from zinc binding of the α-carbonyl, which is polarized and positioned for hydrolysis by water. This hypothesis was supported by an AlphaFold model, in which the α-carbonyl of aspartimide was suggested for zinc-coordination (Fig. S25). We note that *Ec*YbeY was able to convert *Ec*uS11 N119Q variant to an isoGlu-containing product to a minor extent. Compared with the five-membered aspartimide intermediate, formation of a six-membered glutarimide is kinetically less favorable. This may explain why *Ec*uS11 N119Q variant is a poorer substrate for *Ec*YbeY. Nevertheless, the small amount of an isoGlu-containing product formed by *Ec*YbeY suggests an alternative enzymatic route to isoGlu formation beyond the previously reported side product of Gln degradation in human brain tissue.^51^ Although we suggest the D119-G120 motif as a possible YbeY substrate in certain bacteria (Fig. 4b), the *Ec*uS11 N119D variant was not a substrate for *Ec*YbeY. Computational studies on nonenzymatic aspartimide formation revealed activation energies of ∼15 kcal/mol for Asn versus ∼26 kcal/mol for Asp.^52,53^ *Ec*YbeY likely reduces these barriers, but the larger energetic penalty for Asp modification relative to Asn likely persists. Characterization of YbeY from bacteria that naturally encode a uS11 D119-G120 motif is needed to establish whether YbeY from these species may be able to enzymatically modify D119.

The pair of YbeY and the Asn-Gly motif in uS11 is widely distributed in bacteria and in bacteria-derived eukaryotic organelles, indicating that isoAsp formation in uS11 is prevalent and important for biological processes. In archaea and eukaryotes, Asn is replaced by Asp, implying that a different enzymatic pathway operates for isoAsp formation. We speculated that Fap7 catalyzes isoAsp formation in uS11 under circumstances where YbeY is not feasible. We find Fap7 and YbeY do not generally co-occur in bacteria (Fig. 4), and in eukaryotes they are not co-localized: YbeY is targeted to organelles (chloroplasts/mitochondria),^42,44^ whereas Fap7 functions in nuclear ribosome biogenesis.^47^ The catalytic function of Fap7 is supported by the isoAsp residue observed in the remodeled structure of the archaeal Fap7–uS11 complex (Extended Fig. 7).^24^ Previous studies indicated that only intact uS11 stimulated Fap7 ATPase activity, whereas a uS11 variant lacking the C-terminal Asp-containing tail did not.^49^ These observations suggest an ATP-dependent catalytic mechanism (Scheme 1), in which the β-carboxylate of Asp is phosphorylated and then undergoes nucleophilic attack by the amide nitrogen of the +1 residue. An aspartimide intermediate is tentatively formulated and hydrolyzed by water to release phosphate and the isoAsp product.

NMR titration revealed a stable complex between *Tm*YbeY and *Tm*uS11, consistent with previous studies of the human mitochondrial *Hs*YbeY–*Hs*uS11m complex.^44^ Complex formation likely promotes extensive contacts and additional conformational changes in *Tm*YbeY, as indicated by perturbations of numerous backbone amide resonances upon addition of *Tm*uS11. A similarly stable complex was reported for Fap7–uS11.^24,49^ Archaeal Fap7 binds uS11 with picomolar affinity, and this interaction protects uS11 from premature and nonspecific RNA interactions and in particular shields the otherwise highly protease-susceptible C terminus.^49^ We suspect that YbeY also prevents premature release of uS11 before it is assembled into the 30S ribosomal subunit, which may explain why stoichiometric amounts of isoAsp are observed in mature ribosomes and why endogenous PIMT cannot repair this modification.

During ribosome biogenesis in bacteria, the rRNA is produced as a single transcript that is rapidly cleaved into three precursor fragments which are further processed into mature 23S, 5S and 16S rRNAs.^54^ The 17S rRNA precursor to mature 16S rRNA contains 115 extra nucleotides at the 5′ end and 33 extra nucleotides at the 3′ end.^54^ While the 5′ processing enzymes have been identified as RNase E and RNase G, 3′ end processing has been less well understood, and several enzymes have been proposed for this role.^55^ Multiple studies,^40,56^ including the present work, have shown that YbeY-deficient *E. coli* have growth defects, impaired *in vitro* translation activity, and defective ribosome assembly. These findings, coupled with the observation that ΔY-ribosomes contain unprocessed 17S rRNA led to the annotation of YbeY as the enzyme responsible for 3′ end maturation of the 17S rRNA.^28,40^ In this study, we have demonstrated and extensively characterized the role of YbeY in modifying the uS11 residue N119 to IAS119. The 21 r-proteins associated with the 30S subunit begin binding the 17S rRNA in hierarchical manner even before the rRNA is fully processed. Early binding proteins initially bind to the 17S rRNA and drive conformational changes that create binding sites for mid- and late-binding r-proteins.^57–59^ *In vivo*, assembly factors such endonucleases, GTPases, and rRNA and r-protein modifying enzymes facilitate maturation and folding, and the deletion of many of these factors result in growth defects, impaired translation, loss of r-proteins, and accumulation of unprocessed rRNA. For instance, the deletion of the GTPase assembly factor YjeQ results in loss of 30S r-proteins, and the accumulation of unprocessed 17S rRNA that prevents 70S formation by distortion of helix 44 (h44) and occlusion of the inter-subunit bridges B3 and B2a.^60^

ΔY-ribosomes show impaired 70S formation, and analysis of rRNAs indicate the presence of unprocessed 17S rRNA in unassociated 30S subunits but not in fully associated 70S ribosomes. In addition, the reported cryo-EM structure does not have density extending beyond the mature 3′ end of the 16S rRNA, or the distortion of h44 observed upon deletion of assembly factor YjeQ.^60^ Rather, the structure reveals that the absence of the isoAsp modification in uS11 results in minimal perturbations to the local structure of the protein and surrounding rRNA. However, while IAS119 packs against nucleotides 716-718 with the α-carboxylate pointed away from the pocket, the unmodified N119 positions the β-amide towards the rRNA and forms hydrogen bonds with the sugar-phosphate backbone.

This sidechain reorganization in turn ablates the contact between the α-carboxylate of uS11 IAS119 and R35 of bS21, which is located close to the Shine-Dalgarno (SD) helix, and reported to play a role in translation initiation.^61,62^ Since bS21 is known to readily dissociate in solution,^58^ it often appears disordered in cryo-EM maps, prompting us to apply a low-pass filter of 3.5 Å to our reconstructions. The filtered maps reveal clear cryo-EM density for almost the entire length of bS21 in WT ribosomes but poor density in ΔY-ribosomes.^17^ Focused classification of the cryo-EM map indicates that only ∼20% of the particles harbor bS21. This partial occupancy in ΔY-ribosomes may explain the growth defect observed in this strain since bS21 is an essential gene in *E. coli*.^63,64^ Furthermore, *in vitro* functional assays have shown that *E. coli* ribosomes lacking bS21 have decreased activity compared to WT,^65,66^ consistent with our *in vitro* translation data. Taken together, in the *E. coli* ribosome the isoAsp modification in uS11 appears to anchor bS21 to the 30S subunit, which in turn stabilizes the SD helix, and facilitates translation initiation by the ribosome. Bacteria that lack YbeY and the associated isoAsp modification have likely evolved compensatory mutations to anchor bS21 within the 30S subunit, may not require the Shine-Dalgarno sequence for translation initiation, or may lack bS21 entirely.^67,68^

In conclusion, our work uncovers a prevalent isoAsp modification in uS11 across all three domains of life, supports YbeY as the enzyme responsible for Asn deamidation and isomerization in bacteria and eukaryotic organelles, and suggests that Fap7 mediates Asp isomerization in archaea and eukaryotes. These two distinct yet convergent pathways to isoAsp formation suggest a unique role for this modification in ribosome biogenesis.

## Methods

### Materials

Oligonucleotides were purchased from Integrated DNA Technologies with standard desalting treatment. Sanger DNA sequencing was performed by GENEWIZ from Azenta Life Sciences. Restriction endonucleases, DNA polymerases, T4 DNA ligase, HiFi DNA assembly master mix, and AspN endoprotease were obtained from New England Biolabs. Trypsin protease was purchased from Promega. Amicon Ultra centrifugal filters and C-18 Ziptip were purchased from EMD Millipore. Other chemicals were purchased from Sigma-Aldrich or Thermo Fisher Scientific unless stated otherwise.

### General methods

Protein electrophoresis was conducted using Bio-Rad Mini-PROTEAN systems and stained with Coomassie blue. Matrix-assisted laser desorption/ionization time-of-flight (MALDI–TOF) mass spectrometry analysis was performed using a Bruker Neoflex MALDI TOF mass spectrometer at the Mass Spectrometry Research Core (MSRC) at Vanderbilt University. High-performance liquid chromatography (HPLC) was performed on a ThermoFisher Scientific Vanquish UHPLC system with C18 columns. HPLC-coupled high resolution (HR) MS and MS/MS were collected by a ThermoFisher Scientific Q Exactive Orbitrap Mass Spectrometer in the MSRC at Vanderbilt University. Protein NMR data were collected on a 900 MHz Bruker AV-III NMR spectrometer with a CPTCI probe in the Biomolecular NMR Facility at Vanderbilt. Isothermal titration calorimetry experiments were carried out on a TA Instruments Affinity ITC in the Vanderbilt Center for Structural Biology Labs and Instrumentation Facility.

### AlphaFold3 proteome-wide interaction screening

The *E. coli* uS11 sequence was used as a query sequence in proteome-wide pairwise complex-structure predictions against each protein in the *E. coli* K-12 reference proteome (UniProt proteome: UP000000625). Predictions were performed using AlphaFold 3 version 3.0.0 with Google DeepMind-provided model parameters. Precomputed unpaired MSAs extracted from the AlphaFold Protein Structure Database^69^ were supplied directly in the input JSON files. No paired MSAs or structural templates were used. Five diffusion samples were generated from a single model seed for each bait–prey pair. The mean ipTM score was averaged across the five resulting models. The PAE plots for the pairwise complex predictions were visualized using the PAE viewer.^70^

### Construction of the *E. coli* Δ*ybeY* strain

In-frame deletion of *ybeY* was constructed using the established λ-Red recombination system.^71^ The primers used for the *ybeY* deletion are listed in Table S3. A kanamycin-resistant Δ*ybeY* mutant was derived from the *E. coli* K-12 (BW25113) parental strain. The genotypes were verified by PCR and by Sanger DNA sequencing.

### Ribosome isolation

LB medium (1 L) supplemented with 34 µg/mL kanamycin sulfate was inoculated with 50 mL of *E. coli ΔybeY* preculture and incubated at 37 °C for 3.5 h (OD_600_ = 0.8). The culture was then centrifuged at 6,000 x *g* for 30 min at 4 °C, and the pellet was resuspended in ribosome buffer A (20 mM Tris-HCl pH 7.5, 100 mM NH_4_Cl, 10 mM MgCl_2_). The cells were lysed by sonication and subjected to centrifugation in an F14-14×50cy rotor (ThermoFisher) at 11,000 x *g* for 30 min at 4 °C. The supernatant was then split evenly and layered over two identical sucrose cushions of 24 mL buffer B (20 mM Tris-HCl pH 7.5, 500 mM NH_4_Cl, 10 mM MgCl_2_) with 0.5 M sucrose and 17 mL buffer C (20 mM Tris-HCl pH 7.5, 60 mM NH_4_Cl, 6 mM MgCl_2_) with 0.7 M sucrose in ti-45 tubes (Beckman-Coulter). Crude ribosomes were pelleted in a Ti-45 rotor at 27,000 rpm (57,031 x *g*) for 17 h at 4 °C. The pellets were resuspended in dissociation buffer (20 mM Tris-HCl pH 7.5, 60 mM NH_4_Cl, 1 mM MgCl_2_), flash frozen, and stored at -80 °C. Crude ribosomes (0.5 mg) were layered over each of six 20-50% sucrose gradients in reassociation buffer (20 mM Tris-HCl pH 7.5, 60 mM NH_4_Cl, 15 mM MgCl_2_) and spun at 27,000 rpm (57,031 x *g*) for 17 h at 4 °C in a SW-32 rotor (Beckman Coulter). The gradients were fractionated on a BioComp piston gradient fractionator, and the peaks corresponding to 30S, 50S, and 70S ribosomes were fractionated and collected separately. Each peak was concentrated and then buffer exchanged into reassociation buffer in a 100 kDa molecular weight cut-off spin filter (Millipore). The fractions were then flash frozen and stored at -80 °C. Concentrations of each subunit were estimated using the approximation that 1 A_260_ = 72 nM 30S, 36 nM 50S, and 24 nM 70S.

### Ribosomal RNA isolation and analysis

rRNA was extracted from samples corresponding to 30S, 50S and 70S of WT and ΔY-ribosomes using an equal volume of acidic phenol-chloroform (Invitrogen) followed by back-extraction with chloroform. Samples were then precipitated by the addition of one-tenth volume of 3M KCl and three volumes of 100% ethanol and incubated for 1 h at -80 °C. rRNA pellets were obtained by centrifugation on a bench-top centrifuge at 13,000 rpm for 30 minutes at 4 °C and resuspended in nuclease-free water. rRNAs were run on a 4% agarose gel stained with SYBR-safe.

### Nanoluciferase reporter assay

Nanoluciferase (nLuc) was generated using *in vitro* transcription/translation (IVTT) from an expression plasmid DNA template containing a T7 promoter, a ribosome binding site, and the coding sequence for nLuc. IVTT reactions were performed at a 12.5 µL scale using a Δribosome PURExpress kit (NEB). PURExpress reagents (5 μL Reagent A and 1.5 μL factor mix), a 1:50 dilution of Nano-Glo substrate (Promega), followed by 250 nM of associated 70S ribosomes from either *E. coli* WT or Δ*ybeY* were added to the reaction mixture on ice. The reaction was initiated by the addition of 125 ng of DNA template, and 2 µL of the above reaction mixture was immediately aliquoted into four wells of a 384-well plate on ice. Luminescence was measured over time at 37 °C in a Spark Plate Reader (Tecan). Initial slopes were calculated from the linear region of the increase in luminescence signal.

### Ribosomal protein crude preparation

Briefly, 400 µg of 30S subunits and 70S ribosomes from WT and *E. coli ΔybeY* strains were treated with 66% acetic acid and 0.1 M magnesium acetate for 2 h on ice, followed by pelleting ribosomal RNA using a tabletop centrifuge. Crude ribosomal proteins were recovered from the supernatant by lyophilization and stored at -80 °C until further analysis.

### HPLC enrichment of native uS11 and MS analysis

The ribosomal protein crude fraction was separated on a C18 column (Thermo Scientific Acclaim, 300 Å, 3 μm, 4.6 x 150 mm) using an HPLC system. Solvents A (water) and B (acetonitrile) both contained 0.1% formic acid (FA). The following gradient was used at 1 mL/min: held at 10% B for 5 min, 10–60% B in 35 min, 60–90% B in 5 min, held at 90% B for 3 min, 90–10% B in 2 min, and held at 10% B for 10 min. The fractions were continuously collected per minute, dried out by lyophilization, and resuspended in a 1% TFA solution. Samples were mixed with α-cyano-4-hydroxycinnamic acid (CHCA), and the spots were analyzed by MALDI–TOF MS in a linear positive mode (5-20 kDa) using 20 keV of total acceleration energy. External calibration was performed using a combination of cytochrome c and myoglobin with the same CHCA matrix.

For tryptic digestion analysis, fractions containing uS11 were resuspended with 50 mM NH_4_HCO_3_ and mixed with 0.5 μg trypsin. Reactions were allowed to proceed at 37 °C for 2 h before quenching with 1% FA. Samples were desalted by C18 ziptip, mixed with CHCA, and analyzed by MALDI–TOF MS in a reflector positive mode (700-3500 Da) using 20 keV of total acceleration energy and a 25 kV reflectron voltage. External calibration was performed using the peptide calibration standard (ProteoMass, Sigma-Aldrich). Fragmentation (MS/MS) of selected ions was recorded using the MALDI-TOF/TOF method (the “LIFT” technology), and 6,000 laser shots were accumulated to create the spectra. MALDI-TOF/TOF spectra were processed using the Bruker flexAnalysis software.

### Plasmid preparation and construction

Plasmid construction was conducted using a T4 ligation strategy or Gibson assembly, except for site-directed mutagenesis, which was performed using the QuikChange method. Genes encoding *E. coli* YbeY and uS11 were amplified by PCR from *E. coli* K-12 and cloned into a pET28 vector with an N-terminal MBP-His_6_ tag and a pRSFDuet vector with an N-terminal His_6_ tag, respectively. *Ec*uS11 WT and variants were subcloned into pBAD24 vectors with an N-terminal His_6_ tag for expression in *E. coli* K-12 Δ*ybeY* strains. Genes encoding *Ec*YbeY WT and variants were subcloned into pBAD24 vectors without any purification tag for plasmid-based expression in *E. coli* K-12 Δ*ybeY* strains. A DNA sequence encoding the amino acid sequence of I_110_TDVTPIPHN_119_GCRPPKKRRV_129_ was inserted into a pET28 vector with an N-terminal MBP-His_6_ tag for C-terminal peptide production. DNA sequences encoding the amino acid sequence of RI_107_TNITDVTPIPH(N/D)_119_GCRPPKKRRV_129_ were inserted into the pET28 vector with an N-terminal MBP-His_6_ tag for N119 or D119-containing standard peptide production. Genes encoding *T. maritima* YbeY and uS11 were codon-optimized to *E. coli* and synthesized by Genewiz (Azenta US, Inc.). *Tm*YbeY and *Tm*uS11 were individually inserted into a pRSFDuet vector with an N-terminal His_6_ tag. *E. coli* DH5α strains were used for all cloning purposes. The primers used in this study are listed in Table S3. All constructs were verified by Sanger DNA sequencing.

### Protein expression and purification

The plasmids encoding *Ec*YbeY WT and variants were transformed into *E. coli* BL21(DE3) cells for protein expression. Cells were grown overnight on Luria–Bertani (LB) agar plates with appropriate antibiotics at 37 °C. A single colony was picked to inoculate 10 mL of liquid LB with appropriate antibiotics and grown overnight at 37 °C, 250 rpm. The seed culture was used to inoculate 1 L of LB with appropriate antibiotics at 37 °C, 250 rpm. Isopropyl β-D-1-thiogalactopyranoside (IPTG) was then added to a final concentration of 0.5 mM when cells grew to an OD_600_ of 0.6–0.8 at 37 °C. Cells were further cultivated overnight at 18 °C, 250 rpm, before pelleting by centrifugation at 4,000 x *g* for 10 min at 4 °C. For overexpression of *Ec*uS11 C-terminus peptides, similar conditions were used except that 1 mM IPTG was used for induction at 37 °C for 4 h. *Tm*YbeY and *Tm*uS11 were expressed in *E. coli* BL21(DE3) cells using 1 mM IPTG and induction at 30 °C overnight. *Ec*uS11-related constructs were transformed into *E. coli* K-12 Δ*ybeY* strains for overexpression. L-arabinose was added to a final concentration of 0.2% (w/w) when cells grew to an OD_600_ of 0.6–0.8 at 37 °C. Cells were further cultivated overnight at 30 °C, 250 rpm, before pelleting by centrifugation at 4,000 x *g* for 10 min at 4 °C.

For purification of MBP-His_6_-tagged proteins, cell pellets were thawed on ice and resuspended in 30 mL of lysis buffer A (50 mM Tris, 0.5 M NaCl, 2.5% glycerol (*v/v*), pH 8.0) containing 0.5 mM phenylmethylsulfonyl fluoride (PMSF). Cells were then gently rocked for 30 min at 4 °C before 4 x 45 s of sonication, with 10 min of rocking between sonication rounds. Cellular debris was removed by centrifugation (20,000 x g, 45 min x 2). The supernatant was mixed with pre-equilibrated Ni-NTA resin and gently rocked for 30 min at 4 °C. The column was first washed with 10 column volumes (CV) of lysis buffer, then 30 CV of wash buffer A (50 mM Tris, 0.5 M NaCl, and 30 mM imidazole, pH 8). The protein of interest was eluted using 10 CV of elution buffer (50 mM Tris, 0.5 M NaCl, and 250 mM imidazole pH 8). The eluate was concentrated using an Amicon ultracentrifugal filter (EMD Millipore) with a 30 kDa molecular weight cutoff. Protein of interest was buffer exchanged using a HiTrap Desalting column (Cytiva) in storage buffer (50 mM HEPES, 150 mM NaCl, 2.5% glycerol (*v/v*) pH 7.5). His_6_-*Tm*YbeY was expressed and purified in the same way as described above.

His_6_-*Tm*uS11 was expressed overnight at 30 °C and then purified using a denaturing method. For denaturing purification, cell pellets were resuspended in 5 volumes (w/v) of lysis buffer B (6 M guanidine hydrochloride (Gn-HCl), 20 mM Na_2_HPO_4_, 150 mM NaCl, pH 8.0, 0.5 mM PMSF) and disrupted by sonication. Cellular debris was pelleted by centrifugation, and the supernatant was incubated with pre-equilibrated Ni-NTA resin for 30 min at 4 °C. The column was first washed with 5 CV of wash buffer B (4 M Gn-HCl, 30 mM imidazole, 20 mM Na_2_HPO_4_, 150 mM NaCl, pH 8.0), then 20 CV of wash buffer A. The protein of interest was eluted using 10 CV of elution buffer. The eluate was concentrated and buffer-exchanged in the storage buffer. *Ec*uS11-related proteins were purified using the denaturing method described above. Protein concentrations were estimated using both the Bradford assay and A280. Liquid nitrogen was used to flash-freeze protein aliquots before storage at -80 °C. The purity of proteins was inspected by SDS–PAGE analysis and Coomassie blue staining (Fig. S3).

### *In vitro* reconstitution of isoAsp formation in uS11 by YbeY

*In vitro* assays were performed using 1 μM MBP-*Ec*YbeY WT or variants with 10 μM *Ec*uS11 purified from *E. coli* K-12 Δ*ybeY* in a total of 50 μL of 50 mM Tris pH 7.5, 100 mM NaCl, 1 μM Tobacco Etch Virus (TEV) protease, and 2 mM DTT. EDTA was added to a final concentration of 5 mM when required. Reactions were allowed to proceed at 30 °C for 1 h and digested by trypsin (1:20 w/w) for another 2 h at 37 °C. The reaction mixture was desalted by C18 ziptip and analyzed by MALDI–TOF MS or LC–LCMS. For the *T. maritima* system, the reaction conditions were the same as above, except that *Tm*uS11 was purified from BL21(DE3). *Ec*uS11 variants were assayed under the same reaction conditions described above, except that the *Ec*uS11 and *Ec*YbeY concentrations were adjusted to 10 μM and 5 μM, respectively, and the reactions were extended to 2 h at 30 °C.

### LC–HRMS analysis of tryptic products

Tryptic products of *in vitro* assays were analyzed on a C18 column (Thermo Scientific Hypersil GOLD, 175 Å, 1.9 μm, 50 x 2.1 mm) in the HPLC–HRMS system. Solvents A (water) and B (acetonitrile) both contained 0.1% FA. The following gradient was used at 0.3 mL/min: held at 5% B for 2 min, 5–45% B in 7 min, 45–95% B in 0.5 min, held at 95% B for 2 min, 95–5% B in 0.5 min, and held at 5% B for 2 min. MS1 spectra were collected in the positive mode using the following parameters: resolution, 120,000; maximum IT, 200 ms; scan range, 150–2000 m/z. MS/MS spectra were collected in the positive mode of the PRM method using the following parameters: resolution, 30,000; maximum IT, 100 ms; isolation window, 2.0 m/z; NCE, 20 or 25. External calibration was performed using the positive ion calibration standard (Thermo Scientific Pierce LTQ ESI positive ion calibration solution). HRMS/MS spectra were annotated using the interactive peptide spectrum annotator.^72^

### HPLC isolation, AspN digestion, and MALDI–TOF MS analysis of peptides of interest

Tryptic products of *in vitro* assays were separated on a C18 column (Thermo Scientific Acclaim, 300 Å, 3 μm, 4.6 x 150 mm) in the HPLC system. Solvents A (water) and B (acetonitrile) both contained 0.1% FA. The following gradient was used at 1 mL/min: held at 5% B for 5 min, 5–35% B in 40 min, 35–50% B in 4 min, 50–95% B in 1 min, 95% B in 3 min, 95–5% B in 1 min and held at 5% B for 6 min. The fraction was continuously collected every 30 or 60 seconds, dried out by lyophilization, and resuspended in a 1% TFA solution. Samples were analyzed by MALDI–TOF MS. MS/MS was performed using the MALDI-TOF/TOF method.

For AspN digestion analysis, endoprotease (0.5 μg) was mixed with the peptide of interest (∼ 1 μg) isolated by HPLC in a total of 20 μL of 50 mM NH_4_HCO_3_. Reactions were allowed to proceed at 37 °C for 2 h before quenching with 1% FA. The reaction mixture was desalted by C18 ziptip and analyzed by MALDI–TOF MS and MALDI-TOF/TOF.

### ITC assay

For zinc-binding experiments, EDTA (10 mM) was added to MBP-*Ec*YbeY and incubated at room temperature for 30 min. Excess EDTA was removed by a HiTrap Desalting column (Cytiva). ZnCl_2_ was first made at a concentration of 1 M and serially diluted with the binding buffer before each measurement. The zinc solution (0.75 mM) was degassed and filtered through a 0.22 μm membrane before loading into the syringe. MBP-*Ec*YbeY (50 μM) was loaded into the cell. The titration process was performed at 25 °C by injecting the zinc solution into the cell with intervals (up to 200 s) to ensure that the titration peak returned to the baseline. Titration of ZnCl_2_ into the binding buffer was used as the blank control. MBP-*Ec*YbeY variants were treated and analyzed under the same conditions, except that a half-volume was used for the first injection in the ITC experiments. The titration data were analyzed in the NanoAnalyze software with the independent model.

### Preparation of ^15^N-labeled apo-*Tm*YbeY and unlabeled *Tm*uS11

BL21(DE3) cells for His_6_-*Tm*YbeY protein expression were grown to an OD_600_ of 0.6-0.8 and then pelleted by centrifugation at 4,000 x *g* for 10 min. The pellet was washed with M9 minimal salts. For ^15^N-labeling, the BioExpress Bacterial Cell Media (U-^15^N, 98%; Cambridge Isotope Laboratories, Inc.) was used to resuspend the pellet and grow the cells for 1 h at 30 °C. IPTG was added to a final concentration of 1 mM for induction at 30 °C overnight. The ^15^N-labeled His_6_-*Tm*YbeY was purified the same way as described for other His_6_-tagged proteins. Superdex 75 Increase 10/300 GL column (Cytiva) was used for secondary purification and buffer exchange to the NMR buffer (50 mM MES, pH 6.5, 100 mM NaCl, 2 mM DTT). The obvious monomer peak was collected and evaluated by trypsin digestion and MALDI-TOF MS analysis before NMR analysis. Natural abundance His_6_-*Tm*uS11 was prepared similarly and buffer exchanged to the NMR buffer.

### Chemical shift perturbation by NMR

The chemical shift assignment of *Tm*YbeY was obtained from the Biological Magnetic Resonance Bank (BMRB) accession code 6256.^38^ An ^15^N-TOCSY-HSQC spectrum was collected using apo ^15^N-*Tm*YbeY to support assignment transfer under the NMR conditions used. NMR samples were prepared by titrating *Tm*uS11 into ^15^N-*Tm*YbeY (100 μM) at molar ratios of 0, 0.1, 0.25, 0.5, 0.75, 1, 2, 3, and 5 in a total volume of 220 μL NMR buffer containing 5% D_2_O in 3-mm NMR tubes. ^15^N-^1^H HSQC spectra were acquired at 55 °C. Spectra were processed by the Bruker TopSpin software and analyzed in the CcpNmr AnalysisAssign (v3) software.^73^ Amide groups of the uS11-bound form (holo-*Tm*YbeY) were assigned using the minimum distance approach, where each peak in holo spectra was assigned the identity of the nearest peak in apo-*Tm*YbeY. Chemical shift perturbations (CSPs) were calculated from the HSQC spectra using the CSP analysis macro in CcpNmr AnalysisAssign (v3) software.^73^ The calculation was conducted based on equation (1), in which *Δδ_NH_* stands for the change in amide chemical shifts, *Δδ_N_* is the change in amide nitrogen’s chemical shift, and *Δδ_H_* is the change in amide proton’s chemical shift. CSP values were plotted on the secondary structure of *Tm*YbeY using the SSDraw program.^74^

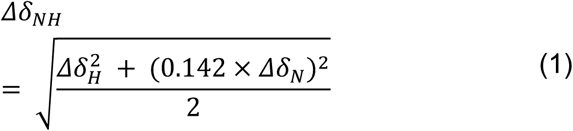

### Growth rate assay

Growth of *E. coli* BW25113 (parental strain), Δ*ybeY*, and Δ*ybeY* strains complemented with a pBAD24 (Amp^R^) empty vector or encoding *Ec*YbeY WT or variants was assessed by monitoring OD_600_. The strains used in this study are listed in Table S4. Strains were streaked on LB agar plates containing the appropriate antibiotics. For strains carrying pBAD24-derived plasmids, carbenicillin (100 µg/mL) was used. Single colonies were used to inoculate 5 mL of LB medium containing the appropriate antibiotics and grown overnight at 37 °C with shaking at 250 rpm. Overnight cultures were diluted 1:100 (250 µL into 25 mL) into fresh LB medium and grown at 37 °C with shaking at 220 rpm. For strains carrying pBAD24-derived plasmids, carbenicillin (100 µg/mL) and L-arabinose (0.1% (w/v)) were included. Aliquots (200 µL) were transferred to clear flat-bottom 96-well plates (Thermo Scientific) for OD_600_ measurement using a Tecan Spark plate reader. Growth was recorded every hour. Each strain was analyzed in three biological replicates.

### Bioinformatic analysis

To annotate bacterial homologs of uS11, YbeY, and Fap7, we searched predicted proteomes of all 189,801 representative bacterial genomes from the Genome Taxonomy Database (GTDB, r232.0) using the homology search tool HMMER 3.4.^75,76^ A list of all homologs can be found in Supplementary File 1 and sequences are provided in Supplementary Files 2-4. To identify uS11 homologs, we searched with the uS11 TIGRFAM profile hidden Markov model (HMM) (TIGR03632) using the HMMER hmmsearch program with an e-value cutoff of 1e-3.^77^ 85.3% of bacterial genomes had exactly one protein hit, consistent with uS11 being a bacterial single-copy gene. For each genome, we kept the protein hit with the most significant e-value that covered at least 40% of the TIGRFAM model, resulting in 168,840 hits. To extract the motif surrounding the modified residue, uS11 protein sequences were aligned to the TIGR03632 profile HMM using the HMMER hmmalign program, and consensus positions corresponding to *E. coli* positions 117-120 were extracted. To identify YbeY homologs, we searched with the YbeY TIGRFAM profile HMM (TIGR00043) using the HMMER hmmsearch program with an e-value cutoff of 1e-3. 91.2% of bacterial genomes had exactly one protein hit, consistent with YbeY being a bacterial single copy gene. For each genome, we kept the protein hit with the most significant e-value that covered at least 40% of the TIGRFAM model, resulting in 174,716 hits. To search for Fap7 homologs, we used a custom profile HMM constructed by searching the yeast Fap7 sequence (UniProtKB Q12055) against SwissProt (release 2026_01) using the HMMER phmmer program with an e-value cutoff of 1e-5, aligning the sequences using MAFFT v7.525 with default parameters,^78^ and building a profile HMM using the HMMER hmmbuild program. Searches were performed with the HMMER hmmsearch program with an e-value cutoff of 1e-3, additionally applying a single domain e-value cutoff of 1e-7, resulting in 11,007 hits. To identify which of these hits are Fap7 as opposed to other homologous proteins, we aligned these hits together with the SwissProt Fap7 sequences used to build the profile HMM using MAFFT, trimmed the alignment to remove columns with more than 98% gaps, and built a protein tree using FastTree v2.1.11 with default settings.^79^ Most of the SwissProt Fap7 sequences formed a distinct clade within the tree, clustering together with 35 bacterial sequences, which we considered as candidate bacterial Fap7 homologs. To validate these hits, we used AlphaFold3 to co-fold each candidate Fap7 sequence together with the uS11 gene from the same genome, ATP, and a magnesium ion and individually confirmed the predicted positioning of the uS11 C-terminal tail inside the active site of Fap7 (Extended Fig. 7). Phylogenetic trees were visualized using the Interactive Tree of Life v6 web server.^80^

### Modeling of isoAsp in published cryo-EM reconstructions and crystal structures

For the cryo-EM reconstruction of the *Arabidopsis thaliana* 80S ribosome (PDB: 9H3G),^81^ the deposited coordinates and EM map were downloaded and used for real space refinement of isoAsp into uS11 at residue 137, using the the phenix.real_space_refine command in PHENIX^82^ at a resolution of 1.82 Å and using the relevant .cif and .def files required for linking IAS residues in a protein chain. Information in the required files is given in Supplementary File 5.

To model Fap7 crystal structures with isoAsp in uS11, coordinates and structure factors of PDB entries 4CVN and 4CW7 were used for isoaspartate modeling.^24^ The crystal structures include Fap7 complexed with uS11 from *Pyrococcus abyssi* GE5, along with ATP-Mg^2+^ (4CW7) or ADP-Mg^2+^ (4CVN). In each crystal form, four copies of the Fap7-uS11 complex are present in the asymmetric unit. For 4CVN, difference density maps of uS11 chains E and H showed clear evidence for isoaspartate (IAS) instead of aspartate. IAS was refined in all four uS11 chains in each crystal form using PHENIX (phenix.refine),^82^ eliminating the difference density, but with no significant effect on the Rfree value. For 4CW7, difference density of uS11 chains B, D, F, and H showed evidence for isoaspartate (IAS) instead of aspartate, with chain F providing the clearest difference density. The auxiliary files required for isoAsp residues (IAS) in phenix.refine are included in Supplementary File 5.

### Cryo-EM sample preparation

Reassociated 70S ribosomes (100 nM) in Grid Freezing Buffer (20 mM HEPES-KOH pH 7.5, 50 mM NH₄Cl, 50 mM KCl, 15 MgCl₂, 2 mM DTT) were incubated for 15 min at 37 °C and kept on ice prior to plunge freezing grids. The samples were applied on 300 mesh R1.2/1.3 UltrAuFoil grids (Quantifoil) with an additional layer of float-transferred amorphous carbon support film. The grids were washed in chloroform prior to carbon floating. Before applying the sample, grids were glow discharged in a PELCO easiGlow at 0.37 mBar and 20 mAmp for 12 seconds. The sample (4 μL) was deposited onto each grid and equilibrated for 1 min. Grids were blotted and plunge-frozen in liquid ethane with an FEI Mark IV Vitrobot using the following settings: 4 °C, 100% humidity, blot force 3, and blot time 2 s. Grids were clipped for autoloading and stored in liquid nitrogen.

### Cryo-EM data collection

Cryo-EM data collection parameters are summarized in Table S2. Dose fractionated movies were collected on a Titan Krios G3i microscope at an accelerating voltage of 300 kV and with a BIO Quantum energy filter. Movies were recorded on a GATAN K3 direct electron detector operated in CDS mode. A total dose of 40 e^−^/Å^2^ was split over 40 frames per movie. The physical pixel size was set to 0.67 Å. Data collection was automated with SerialEM,^83^ which was also used for astigmatism correction by CTF and coma-free alignment by CTF. The defocus ramp was set to range between -0.5 and -1.5 μm.

### Image processing

Data processing was performed using cryoSPARC v5 (beta).^84^ Patch Motion Correction was used to motion correct the raw movies and Patch CTF was used to estimate CTF parameters, and micrographs with poorly fitting CTF estimates were omitted. Particles were picked using the Blob-Picker and then extracted at one-fourth binning of the full box size and subjected to 2D refinement to select 70S particles. 3D classification was performed using the Heterogeneous Refinement job using a 70S reference volume generated from the coordinates of PDB entry 1VY4 using EMAN2.^85^ Classes corresponding to clean 70S volumes were selected and subjected to 30S-focused refinement. The particles were polished using Reference Based Motion Correction^86^ and then subjected to a final round of 30S-focused refinement to obtain a final map at 1.91 Å global resolution.

### Pixel size calibration

The pixel size was calibrated in UCSF ChimeraX^87^ using the “Fit to Map” function and the high-resolution maps against the coordinates from the X-ray crystal structure PDB 4YBB.^88^ The best cross-correlation value was obtained at pixel size 0.6395 Å.

### Modeling

The 30S subunit of PDB entry 8EMM was used as the starting model.^89^ Real space refinement of the coordinates was performed in PHENIX^82^ and further adjustments to the model were done manually in Coot,^90^ mainly in the vicinity of uS11 and the surrounding 16S rRNA. Further additions to the model included Mg²⁺ and K⁺ ions and water molecules. The map-vs.-model FSC was calculated in PHENIX.^82^

## Data availability

All data are available in the main text, Supplementary Information, and Supplementary Files. The predicted complex structures discussed in this study have been deposited in the ModelArchive database under dataset ma-yg0ih. Atomic coordinates have been deposited with the Protein Data Bank under accession code 37WT. Cryo-EM maps have been deposited with the Electron Microscopy Data Bank under the accession codes EMD-78572 (30S focus refined map from the *ΔybeY E. coli* ribosome) and EMD-78573 (30S focus refined map of particles with occupied bS21 from the *ΔybeY E. coli* ribosome).

## Acknowledgements

We thank Dr. Satish Nair and Dr. Raphaël Méheust for helpful discussions. This work was supported in part by grants from the National Institutes of Health (GM158411 to D.A.M.) and the National Science Foundation Center for Genetically Encoded Materials (CHE-2002182, CHE-2503885, C.M. and J.H.D.C.). C-GEM supported the ribosome cryo-EM, modeling, and biochemistry experiments. Protein NMR work was supported in part by grants from the NSF-MRI (0922862) for the acquisition of a 900 MHz Ultra-High Field NMR. We thank the University of California at Berkeley Cal-Cryo QB3-Berkeley Core Facility, RRID:SCR_028363 and Dr. Kedar Sharma for help with cryo-EM data collection and management. Y.S. is a Don Brown Awardee of the Life Sciences Research Foundation.

## Author Contributions

Y.X., D.A.M, and J.H.D.C. conceptualized the project. Y.X. and T.W. conducted AlphaFold predictions under the supervision of H.M.. Y.X. and S.O.S. performed biochemical experiments. Y.S. and R.N. performed bioinformatic analysis. Y.X. and M.V performed protein NMR experiments. C.M. conducted ribosome studies and acquired cryo-EM data. J.H.D.C. and C.M. revised wrong uS11 structures. Y.X., C.M., Y.S., and R.N. wrote the initial draft with inputs from all authors. All authors reviewed and edited the paper. D.A.M. and J.H.D.C. were responsible for funding and supervised the project.

---

**Extended Figure 1.**
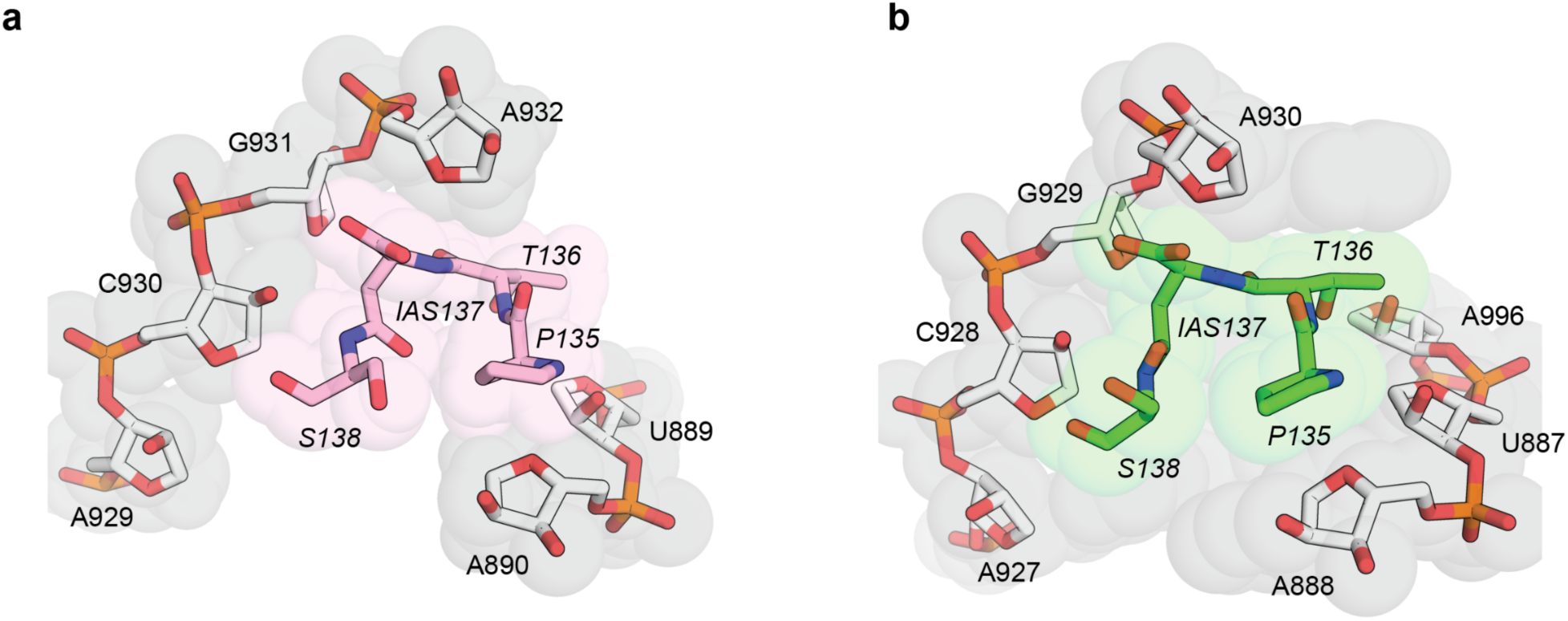
Space-filling model showing the shape complementarity between uS11 and rRNA nucleotides. **a**, tomato, PDB: 7QIX. **b**, thale cress, PDB: 9H3G. The uS11 D137 in thale cress was remodeled as IAS137. 18S rRNA is shown in gray with residues labeled, uS11 is shown in pink for tomato and green for thale cress, and uS11 residues are italicized.

**Extended Figure 2.**
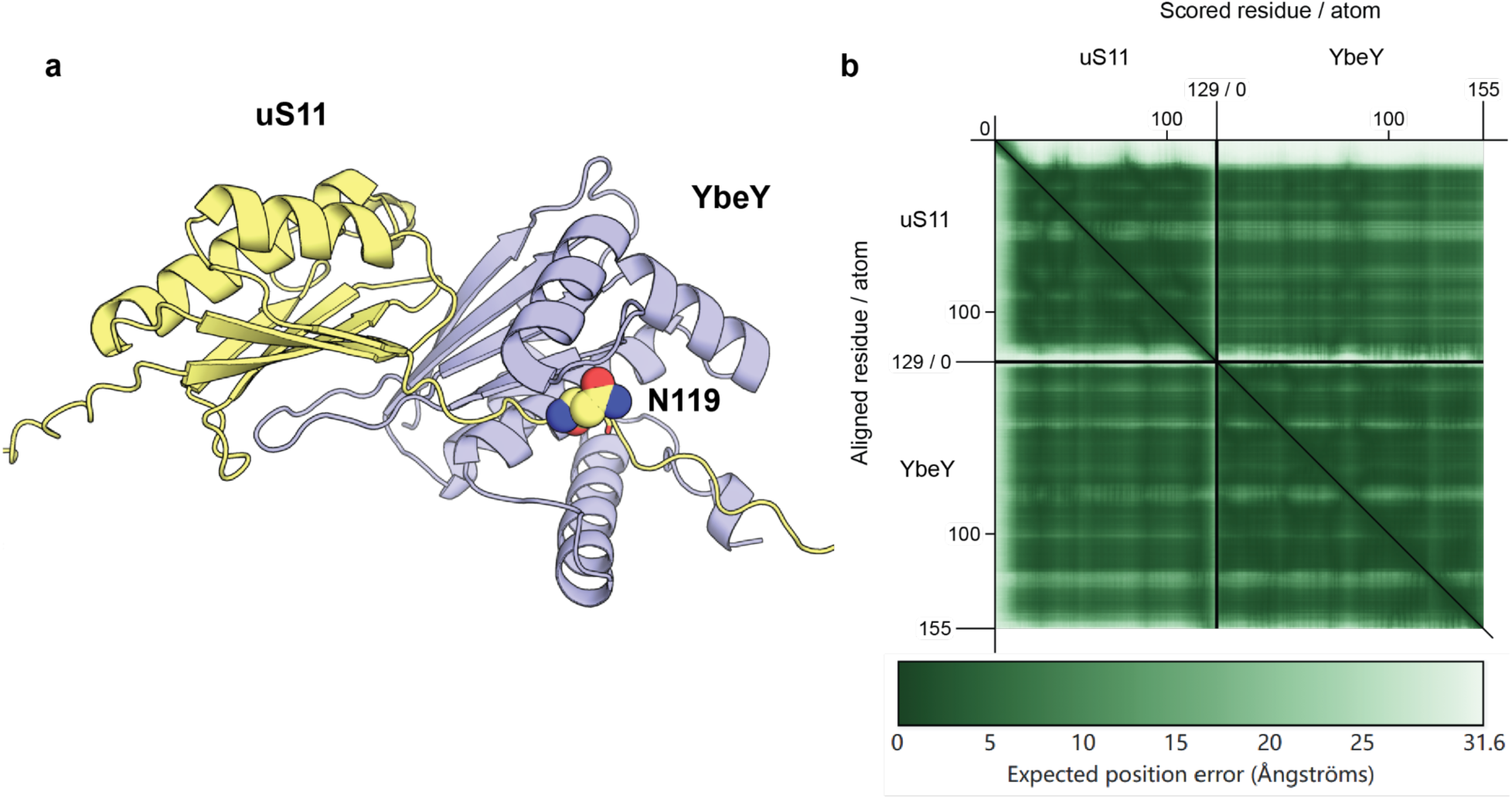
AlphaFold3 model of uS11 and YbeY. **a**, The predicted complex of uS11 and YbeY. uS11 and YbeY are colored in yellow and purple, respectively. N119 in uS11 is highlighted. **b**, The predicted aligned error (PAE) matrix for the predicted model of uS11 and YbeY. The axes ticks indicate residue position in each protein in the model.

**Extended Figure 3.**
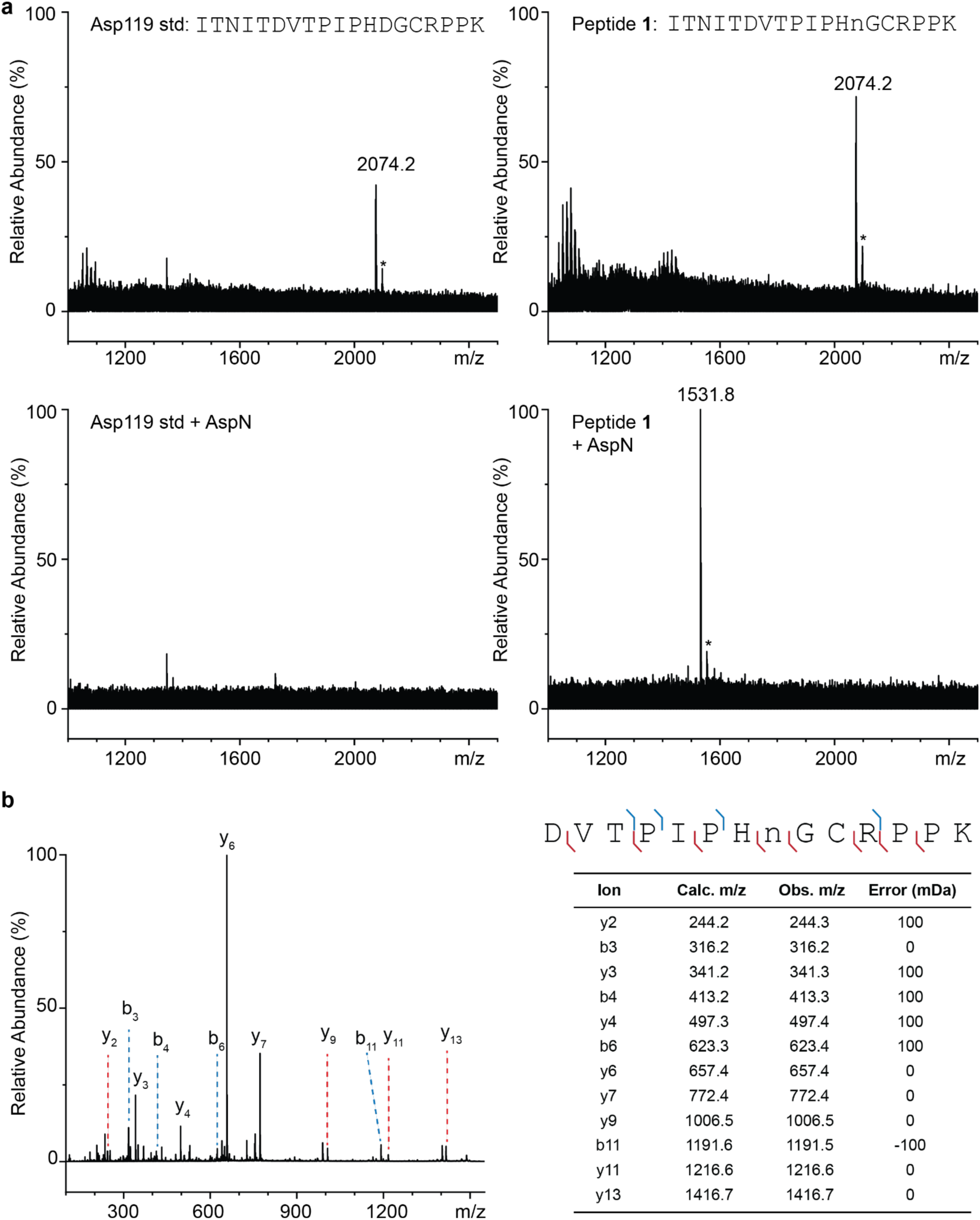
AspN digestion of peptide 1. **a**, D119-containing standard and peptide **1** were digested by protease AspN. Observed m/z values are labeled. Peptide sequences are indicated. For the peptide standard and peptide **1**, calc. [M+H]^+^ = 2074.0749. AspN removed the sequence N-terminal to D112 in peptide **1**, and the remaining peptide sequence was `DVTPIPHnGCRPPK`, with calc. [M+H]^+^ = 1531.7685. Na^+^ adducts are labeled with asterisks. **b**, MALDI-TOF/TOF spectrum of m/z 1531.8. Assigned ions are indicated in the spectrum, sequence, and table. n represents the modified residue. Mass errors were calculated and reported in mDa. *Error* (*mDa*) = (*Observed* − *Calculated*) × 10^3^.

**Extended Figure 4.**
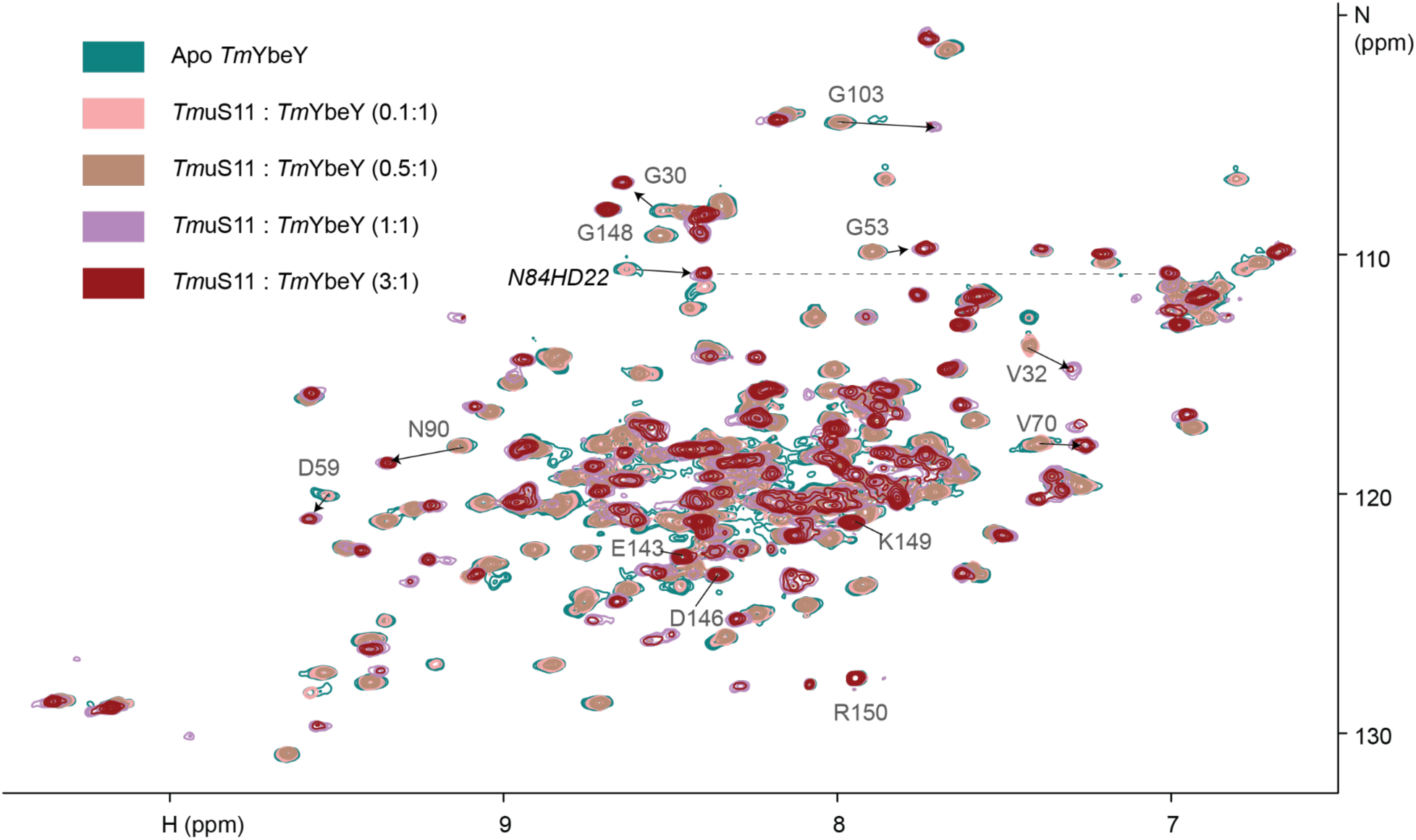
^1^H-^15^N HSQC titration spectra of *Tm*YbeY with *Tm*uS11. Apo peaks of interest are assigned, while the corresponding homo peaks are indicated by arrows. N84 β-amide protons are indicated with dot lines.

**Extended Figure 5.**
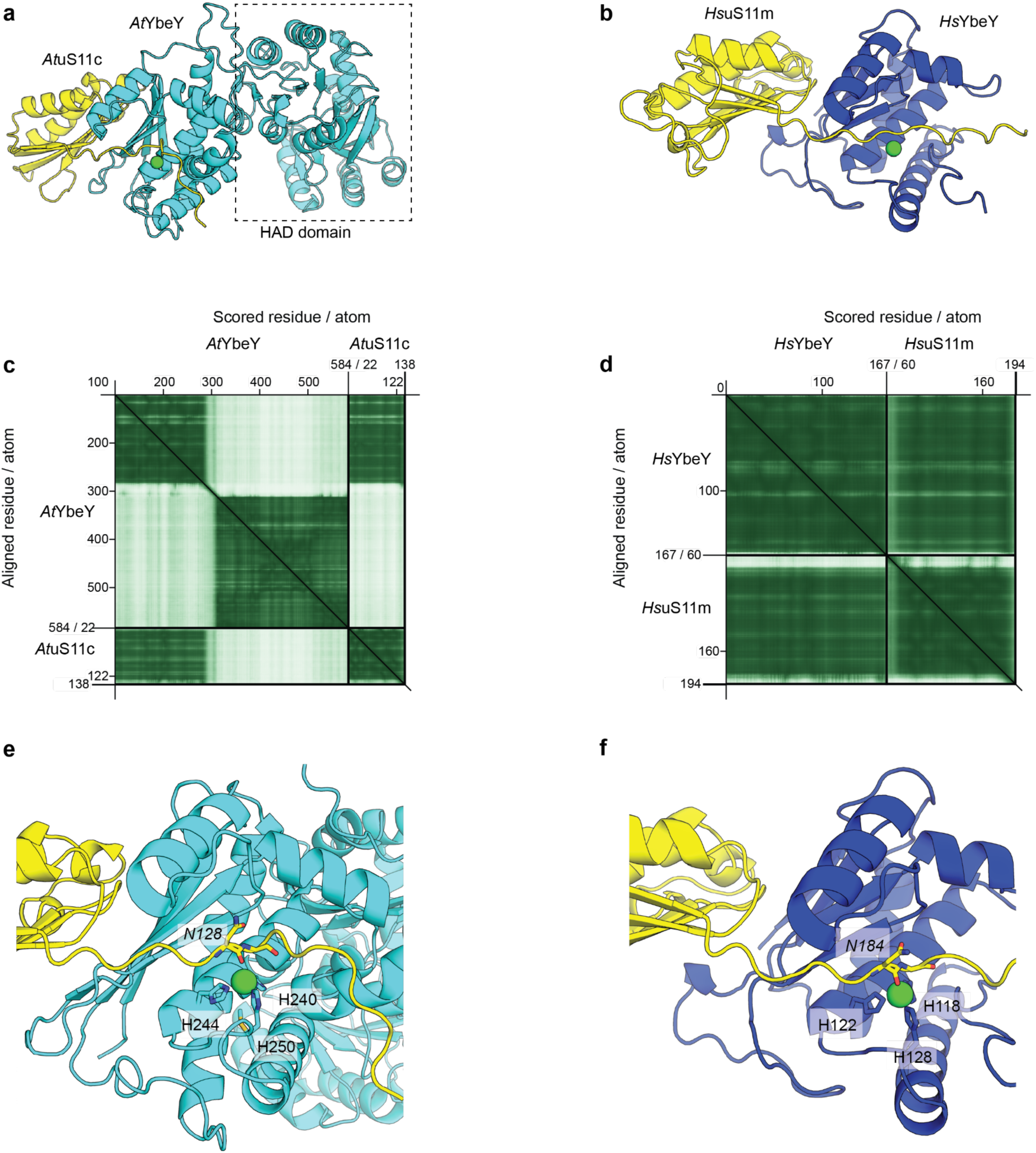
AlphaFold3 models of *At*YbeY–*At*uS11c–zinc and *Hs*YbeY–*Hs*uS11m– zinc. **a,** Overall view of the *At*YbeY–*At*uS11c–zinc model. *At*uS11c, *At*YbeY, and zinc are shown in yellow, cyan, and green, respectively. The C-terminal haloacid dehalogenase (HAD)-like hydrolase domain of *At*YbeY is indicated. The unstructured N-terminal 100 amino acids of *At*YbeY and the N-terminal 22 amino acids of *At*uS11c were removed for the prediction. ipTM = 0.88. **b,** Overall view of the *Hs*YbeY–*Hs*uS11m–zinc model. *Hs*uS11m, *Hs*YbeY, and zinc are shown in yellow, blue, and green, respectively. The unstructured N-terminal 60 amino acids of *Hs*uS11m were removed for the prediction. ipTM = 0.89. **c,d,** PAE matrices for the models shown in **a** and **b**, respectively. Axis tick marks indicate residue positions in each protein. **e,f,** Zoomed-in views of the putative zinc-binding site and the substrate-binding pocket in **a** and **b**, respectively. uS11 residues are italicized.

**Extended Figure 6.**
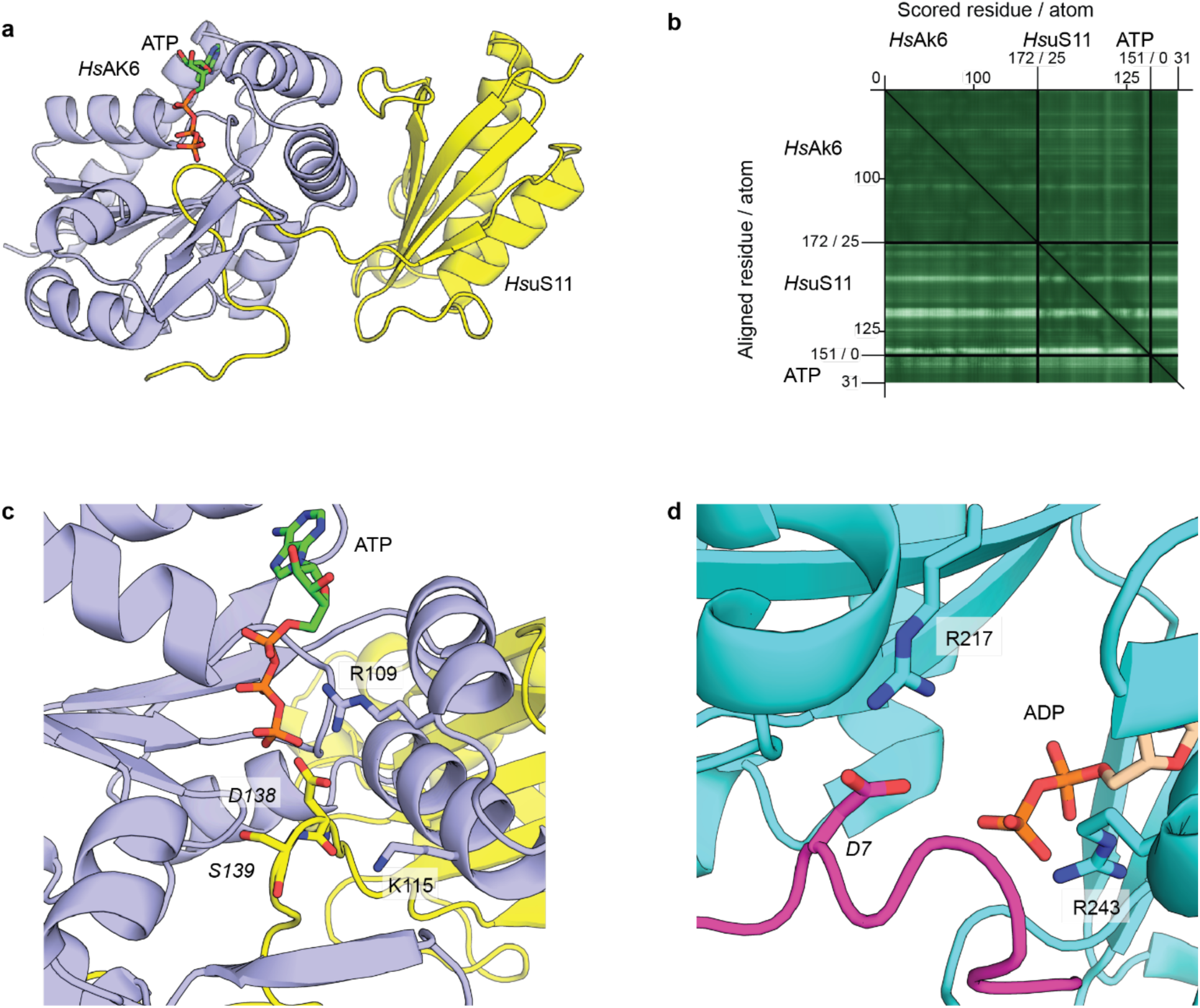
AlphaFold3 models of *Hs*AK6–*Hs*uS11–ATP. **a**, Overall view of the model. *Hs*uS11, *Hs*AK6, and ATP are shown in yellow, purple, and green, respectively. The unstructured N-terminal 24 amino acids of *Hs*uS11 were removed for the prediction. ipTM = 0.90. **b**, PAE matrices for the models shown in **a**. Axis tick marks indicate residue positions in each protein. **c**, Zoomed-in view of the putative ATP-binding site and the substrate-binding pocket in **a**. *Hs*uS11 residues are italicized. **d**, Zoomed-in view of ATP-grasp ligase/substrate complex. PDB: 7MGV. Substrate peptide, ATP-grasp ligase, and ADP are shown in pink, cyan, and wheat, respectively. D7 in the substrate is shown in sticks and labeled in italics.

**Extended Figure 7.**
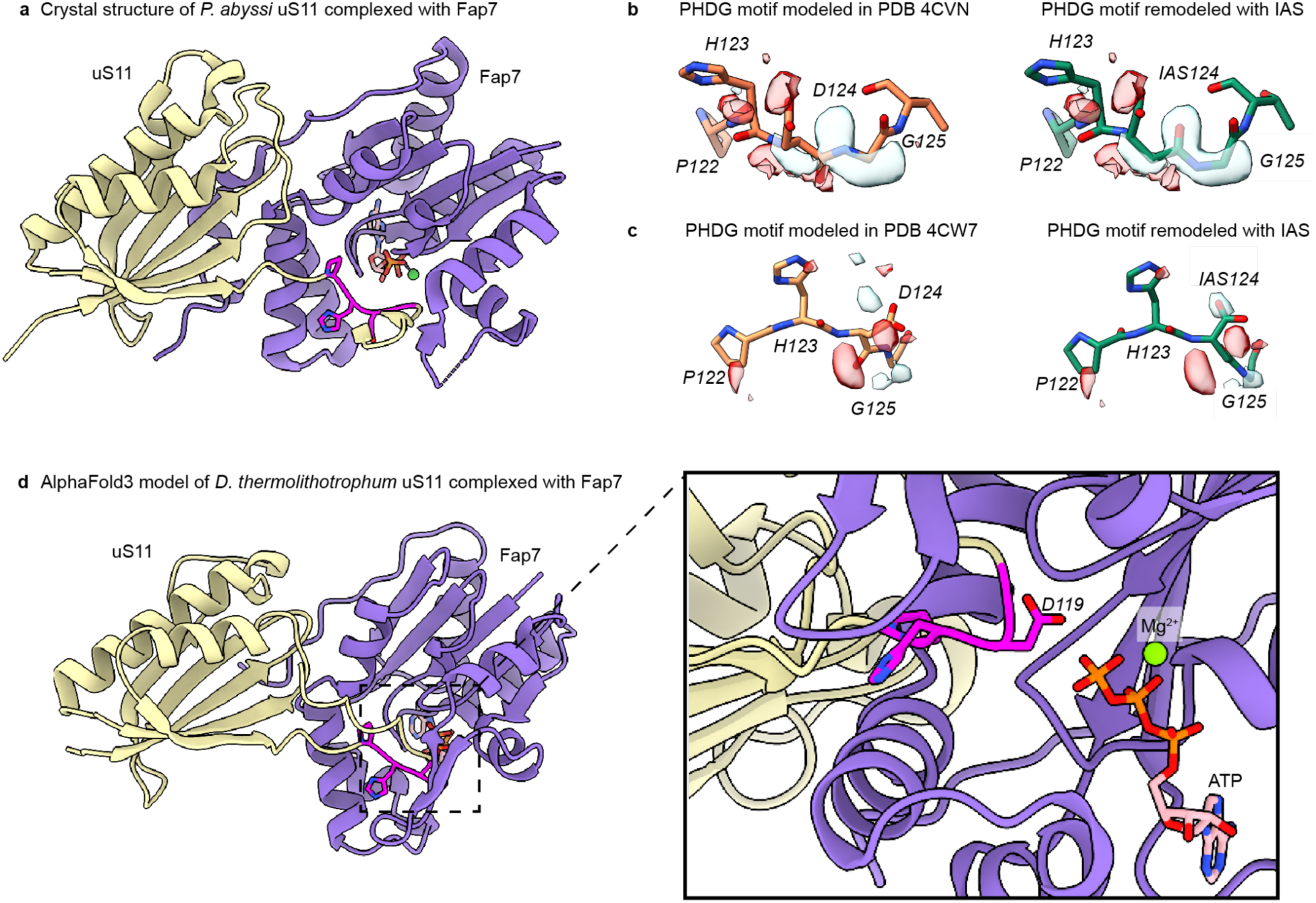
Structural models of archaeal and bacterial Fap7 with uS11. **a**, crystal structure of the archaeal *Pyrococcus abyssi* uS11 (yellow) complexed with Fap7 (purple) in the presence of ADP and Mg^2+^ (PDB ID 4CVN)^24^ showing the C-terminal loop carrying the PHDG motif (magenta) within the Fap7 active site. **b,** Fo-Fc map from 4CVN showing unmodeled density (light blue, left) that is accounted for by remodeling D124 as IAS124 (right). **c,** Fo-Fc map from PDB ID 4CW7^24^ showing unmodeled density (light blue, left) that is accounted for by remodeling D124 as IAS124 (right). **d,** AlphaFold3 structure of the *Desulfurobacterium thermolithotrophum* (RS_GCF_000191045.1) Fap7 homolog (purple) co-folded with the *D. thermolithotrophum* uS11 (yellow), ATP, and Mg^2+^. Inset: zoomed in view of PHDG motif (magenta, *E. coli* numbering) within the active site of Fap7.

**Extended Figure 8.**
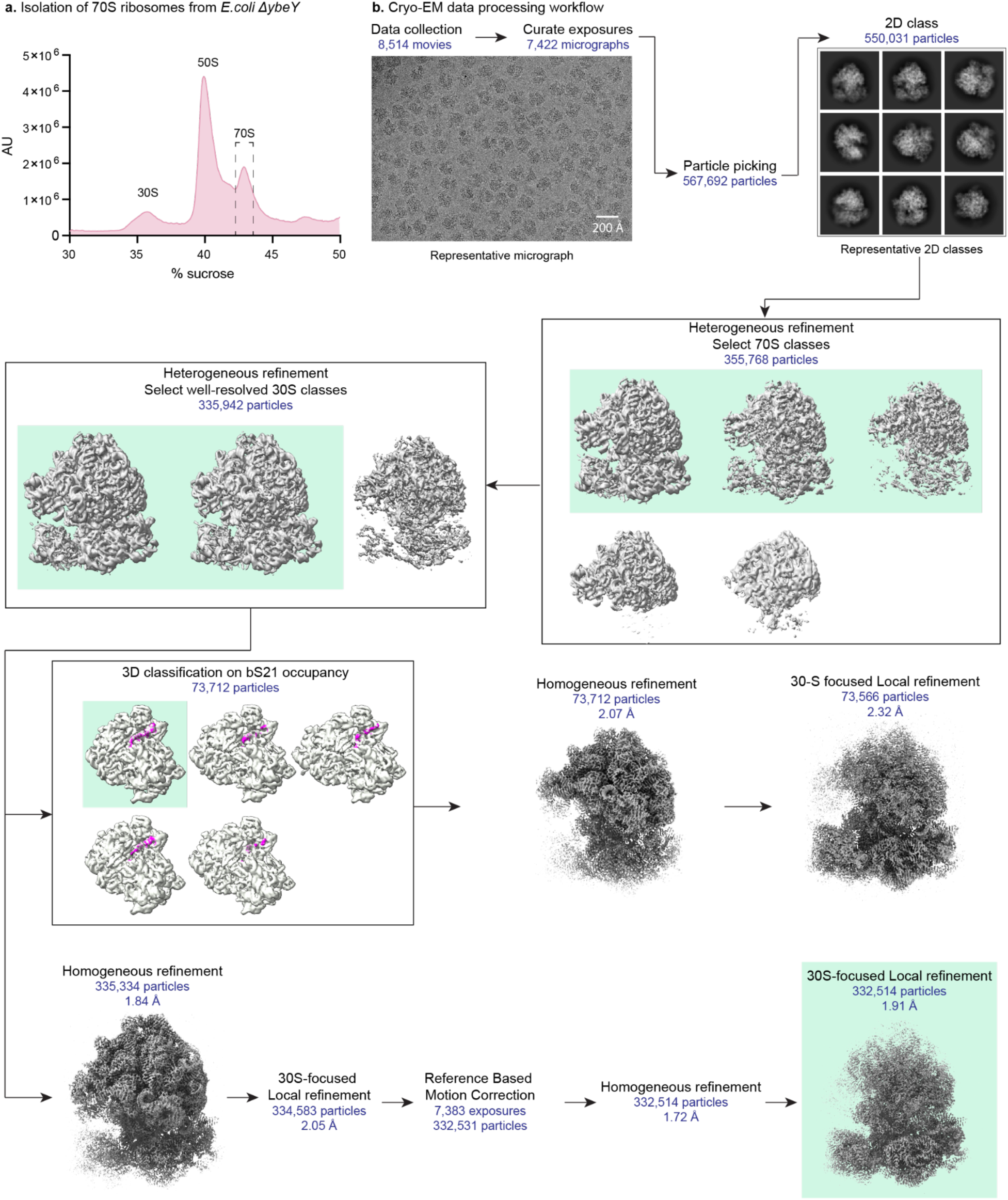
Cryo-EM data processing workflow. **a**, Isolation of associated 70S ribosomes from *E. coli ΔybeY* using a sucrose gradient. **b**, Cryo-EM data processing workflow for the *E. coli ΔybeY* ribosome dataset. A representative micrograph and representative 2D classes from the dataset are shown. Maps chosen for further classification and refinement, and the final refined map are boxed in green. Density for bS21 is shown in magenta.

**Extended Figure 9.**
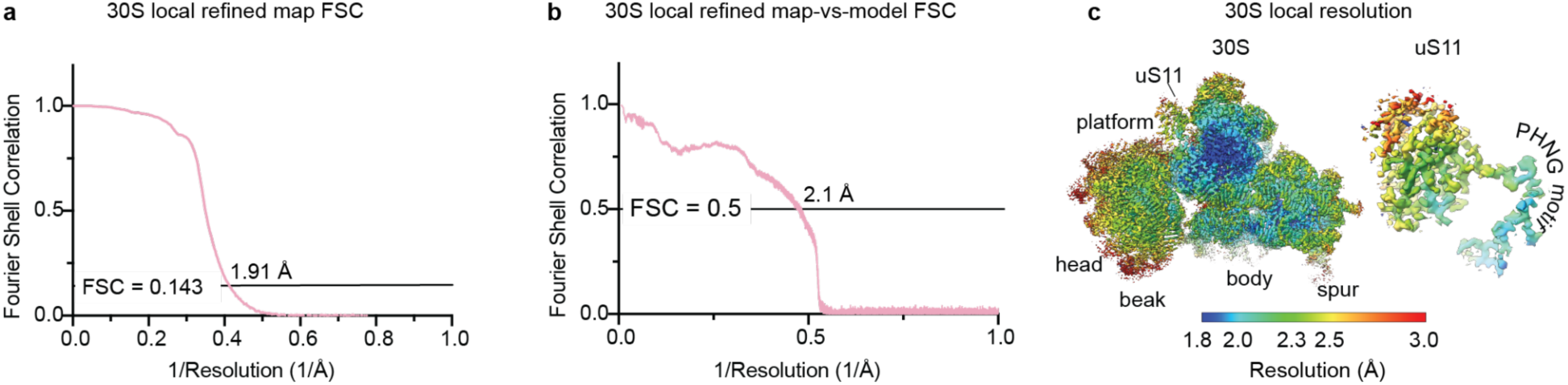
Resolution estimates of ΔY-ribosomes **a,** FSC curve of the 30S local refined map showing global resolution of the map is 1.9 Å at the gold-standard cutoff value of 0.143. **b,** Map-vs-model resolution is at 2.1 Å at FSC cutoff value of 0.5. **c,** 30S subunit color coded by local resolution values. Inset: ribosomal protein uS11 colored by local resolution values.

**Extended Figure 10.**
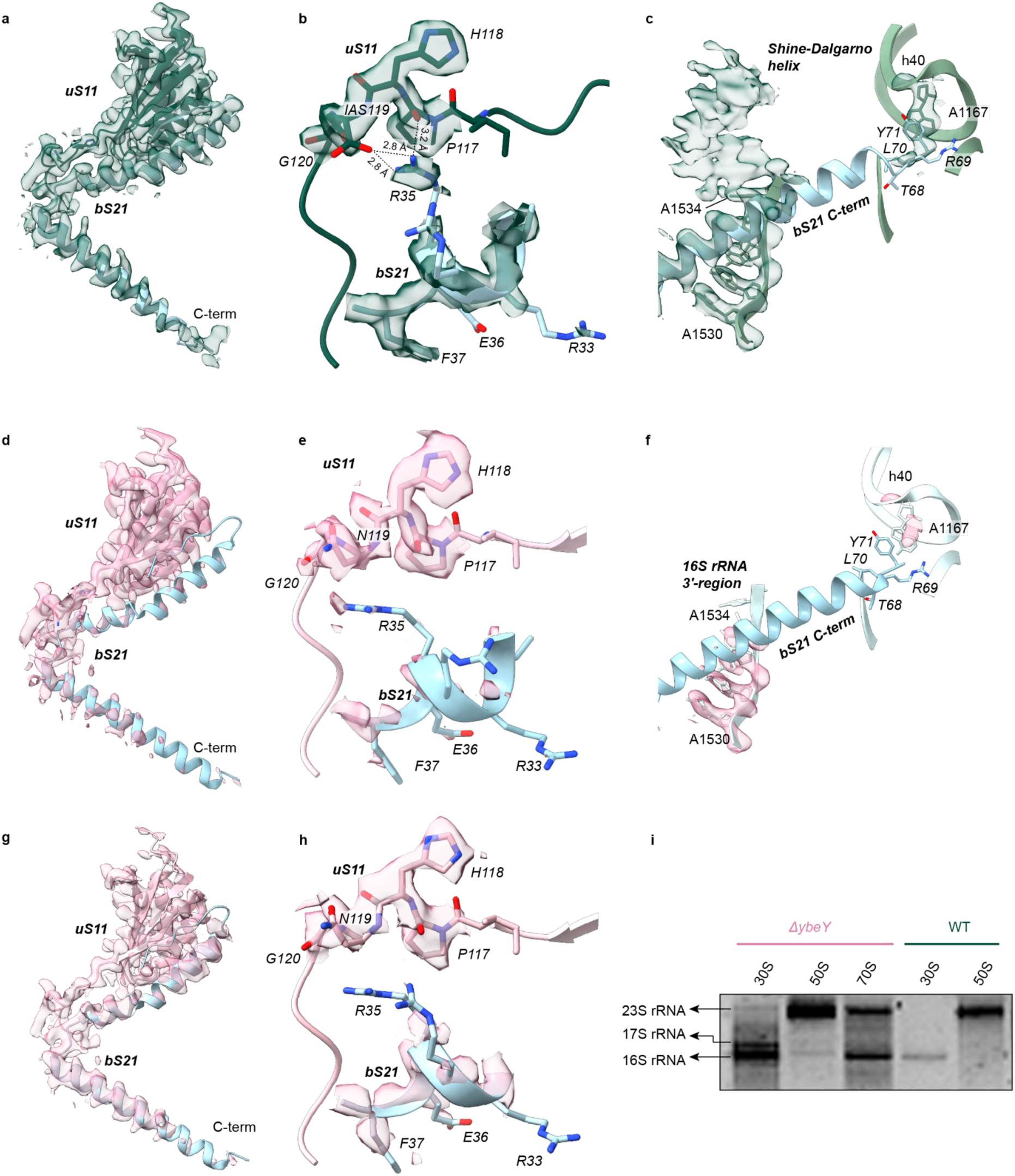
Interaction of uS11 with bS21 and 16S rRNA. **a,** Atomic coordinates and cryo-EM density of uS11 and bS21 in WT *E. coli* ribosomes showing the C-terminal domain of uS11 (green) interacting with bS21 (light blue). **b,** Interface between uS11 and bS21 shows R35 within H-bonding distance of IAS119. **c**, Zoomed view showing the conserved C-terminal RLY motif of bS21 interacting with nucleotide A1167 in helix 40 of the 30S head-domain. Also shown is the 3′-region of the 16S rRNA with density for the Shine-Dalgarno helix. **d,** Atomic coordinates and cryo-EM density of uS11 and bS21 from ΔY-ribosomes showing the C-terminal domain of uS11 (pink) interacting with bS21 (light blue). **e,** Interface between uS11 and bS21 shows that in the presence of unmodified N119, residue R35 is unresolved and loses the H-bonding interactions with uS11. **f**, Zoomed view showing the C-terminal RLY motif of bS21 and A1167 in helix 40 of the 30S head-domain are poorly resolved. The 3′-end of the 16S rRNA with nucleotides 1530-1533 are resolved, but the anti-Shine-Dalgarno sequence from 1534-1541 shows no cryo-EM density. Note: Low-pass filtering to 3.5 Å resolution was applied to all displayed maps to clarify cryo-EM density of bS21. **g**, Classification and reconstruction from particles with occupied bS21 improves its cryo-EM density, however **h**, bS21 R35 residue remains unresolved due to loss of H-bonding interactions with uS11. **i,** Agarose gel showing rRNA extracted from 30S, 50S and 70S particles obtained from *ΔybeY* (pink bar) *E. coli* showing the presence of unprocessed 17S rRNA in 30S subunits from ΔY-ribosomes. No unprocessed rRNA was detected in associated 70S of ΔY-ribosomes; 16S and 23S rRNA from WT ribosomes are shown for comparison.

## Notes

### Competing Interest Statement

The authors have declared no competing interest.

