## Supplementary Information for "Enzymatic formation of a conserved isoaspartate in ribosomal protein uS11"

**Table of contents**


**Supplementary Figures** 3

Figure S1. AlphaFold3 model of uS11 and bS21. 3

Figure S2. AlphaFold3 model of uS11 and YchA. 4

Figure S3. SDS-PAGE analysis of proteins used in this work. 5

Figure S4. MALDI-TOF MS analysis of trypsin-digested uS11 overexpressed in BL21(DE3). 6

Figure S5. MS analysis of uS11 overexpressed in *E. coli* Δ*ybeY*. 7

Figure S6. MS analysis of peptide **1**. 8

Figure S7. MS analysis of peptide **2**. 9

Figure S8. Analysis of the catalytic activity of *Tm*YbeY. 10

Figure S9. $^{1}H$-$^{15}N$ HSQC spectrum of apo *Tm*YbeY at 55 °C. 11

Figure S10. LC–HRMS spectra of uS11 modification by YbeY wild-type and variants. 13

Figure S11. ITC analysis of YbeY zinc-binding affinity. 14

Figure S12. Zoomed-in AlphaFold3 model of *Ec*uS11 and *Ec*YbeY. 15

Figure S13. Shortened C-terminus of Bipolaricaulota uS11. 16

Figure S14. MALDI-TOF MS analysis of the uS11-Cter peptide. 17

Figure S15. LC–HRMS spectra of uS11 variants modified by wild-type YbeY. 18

Figure S16. MS analysis of peptide **3**. 19

Figure S17. MS analysis of peptide **4**. 20

Figure S18. MS analysis of peptide **5**. 21

Figure S19. MS analysis of peptide **6**. 22

Figure S20. MS analysis of peptide **7**. 23

Figure S21. MS analysis of peptide **8**. 24

Figure S22. MALDI-TOF/TOF spectra of peptides **8** and **10**. 25

Figure S23. MS analysis of peptide **9**. 26

Figure S24. MALDI-TOF MS analysis of uS11 isolated from *E. coli* WT (upper) and the Δ*ybeY* mutant (bottom). 27

Figure S25. AlphaFold3 models of *E. coli* YbeY–uS11–aspartimide–zinc. 28

**Supplementary Schemes** 29

Scheme S1. Model for enhanced y ion on the N-terminal side of isoaspartate (n = 1) or isoglutamate (n = 2). 29

Scheme S2. A proposed cysteine-engaged mechanism for isoAsp formation. 30

Scheme S3. A proposed mechanism for isoglutamate formation. 31

**Supplementary Tables** 32

Table S1 Candidates from AlphaFold3 scanning. 32

Table S2 Cryo-EM data collection, refinement, and validation statistics. 35

Table S3 Oligonucleotide primers used in this study. 37

Table S4 *E. coli* strains used in this study. 40

**Supplementary References** 41

**Supplementary Figures.**

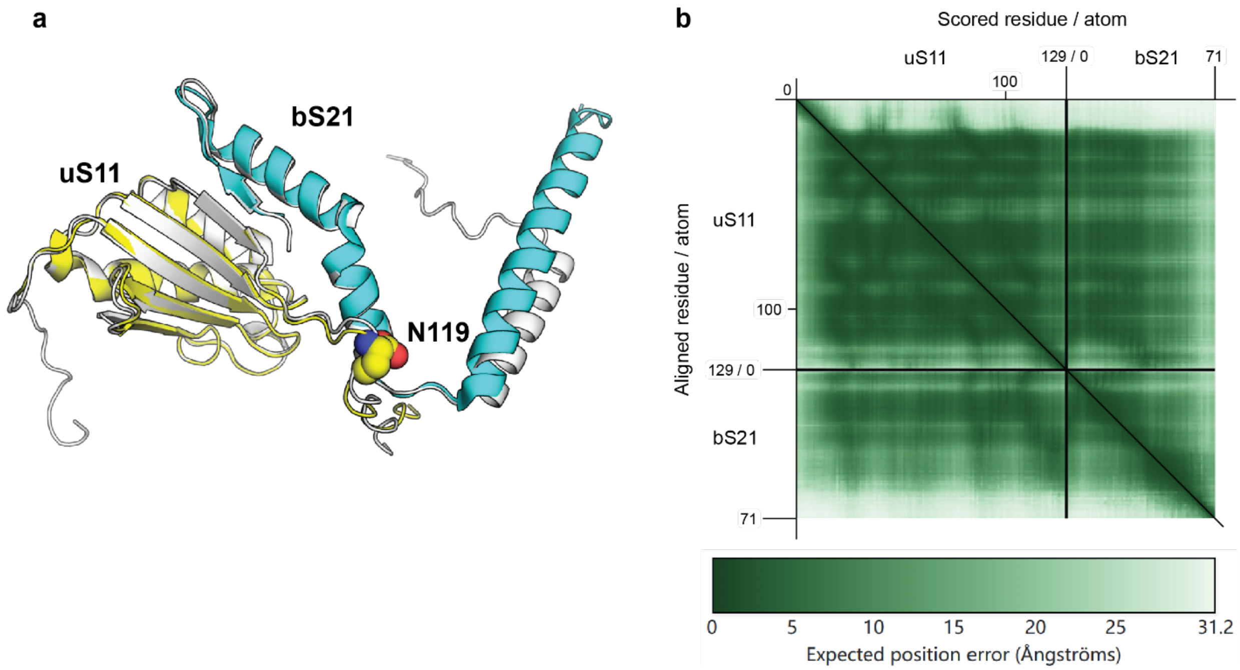


**Figure S1. AlphaFold3 model of uS11 and bS21. a,** Superposition of the predicted model with the cryo-EM structure (PDB: 7K00). The predicted model is in gray, while uS11 and bS21 in the structure are in yellow and cyan, respectively. N119 in uS11 is highlighted. **b,** The predicted aligned error (PAE) matrix for the predicted model of uS11 and bS21. The axes ticks indicate residue position in each protein in the model.

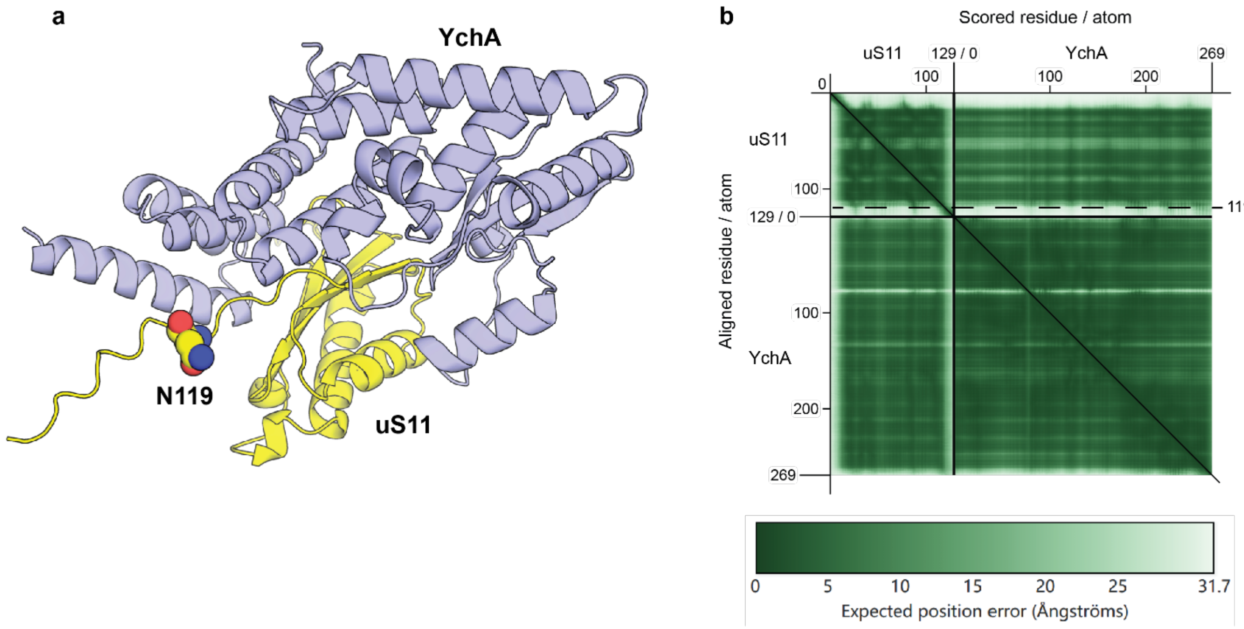


**Figure S2. AlphaFold3 model of uS11 and YchA. a,** The predicted complex of uS11 and YchA. uS11 and YchA are colored in yellow and purple, respectively. N119 in uS11 is highlighted. **b,** The predicted aligned error (PAE) matrix for the predicted model of uS11 and YchA. The axes ticks indicate residue position in each protein in the model.

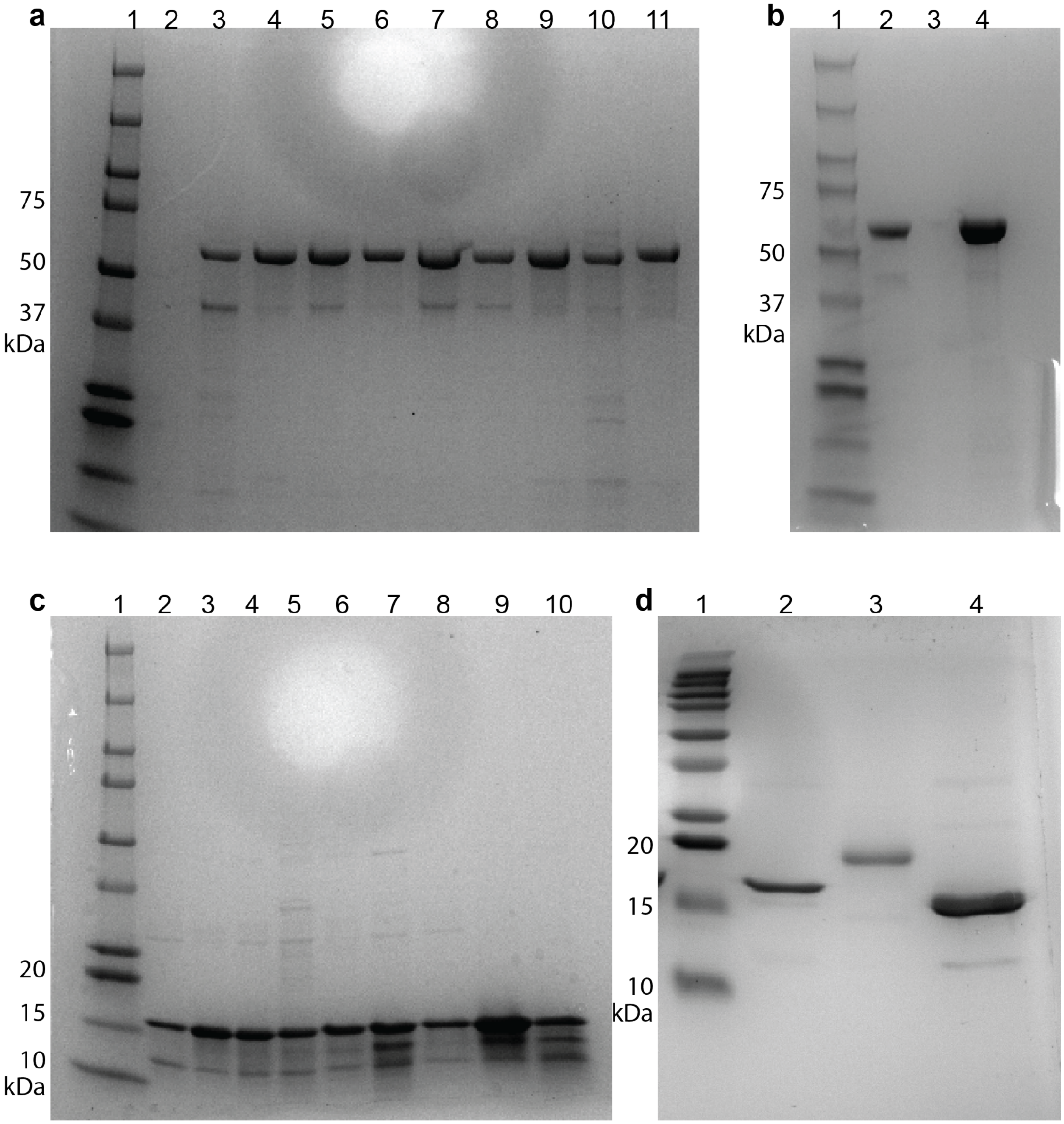


**Figure S3. SDS-PAGE analysis of proteins used in this work. a,** *Ec*YbeY wide-type (WT) and variants were purified with an N-terminal MBP-tag. Lane: 1, ladder (Kaleidoscope, Bio-Rad); 2, loading buffer; 3, WT; 4, R44D; 5, N55A; 6, R59A; 7, N66A; 8, D85R; 9, H114A; 10, H118A; 11, H124A. **b,** *Ec*YbeY variants (continue). Lane: 1, ladder; 2, T42F; 3, loading buffer; 4, T65A. **c,** *Ec*uS11 WT and variants were fused with an N-terminal $His_6$-tag. Lane: 1, ladder; 2, WT; 3, P117A; 4, H118A; 5, N119D; 6, N119E; 7, N119Q; 8, G120A; 9, C121A; 10, R122A. **d,** *Tm*YbeY and *Tm*uS11 were fused with an N-terminal $His_6$-tag. Lane: 1, ladder; 2, *Ec*uS11 WT; 3, *Tm*YbeY; 4, *Tm*uS11. Gels were stained with Coomassie blue.

**a**

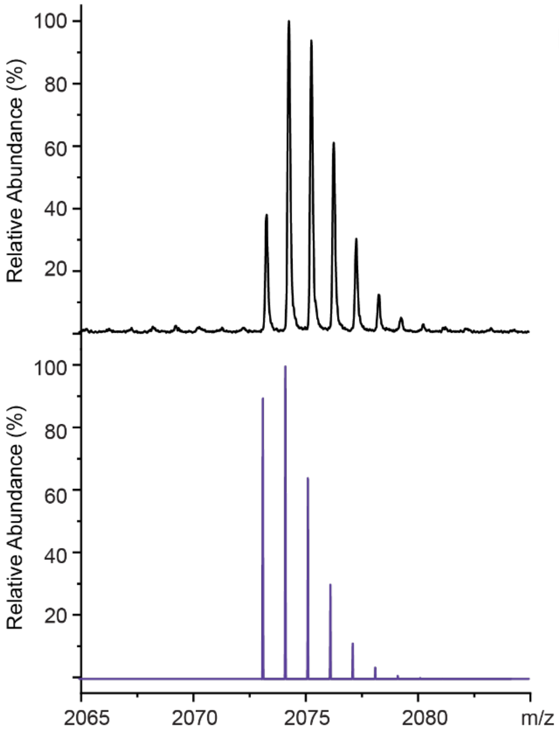


**b**

| Obs. | Abund.% | Calc. | Abund.% |
|---|---|---|---|
| 2073.3 | 38 | 2073.1 | 90 |
| 2074.3 | 100 | 2074.1 | 100 |
| 2075.3 | 94 | 2075.1 | 64 |
| 2076.3 | 61 | 2076.1 | 30 |
| 2077.3 | 30 | 2077.1 | 11 |
| 2078.3 | 12 | 2078.1 | 4 |

**Figure S4. MALDI-TOF MS analysis of trypsin-digested uS11 produced in BL21(DE3). a**, Observed (top) and calculated (bottom) isotopic mass distribution of the peptide of interest. Sequence of interest: $\texttt{ITNITDVTPIPHN}_{119}\texttt{GCRPPK}$. Formula: $C_{90}H_{149}N_{27}O_{27}S$. $[M+H]^+$ = 2073.0909. The observed *m/z* is slightly higher than the predicted value (+0.2), likely due to MALDI-TOF calibration variability. **b**, Isotopic abundances for the observed and calculated mass distributions of the peptide of interest. Abund.%, relative abundance (%).

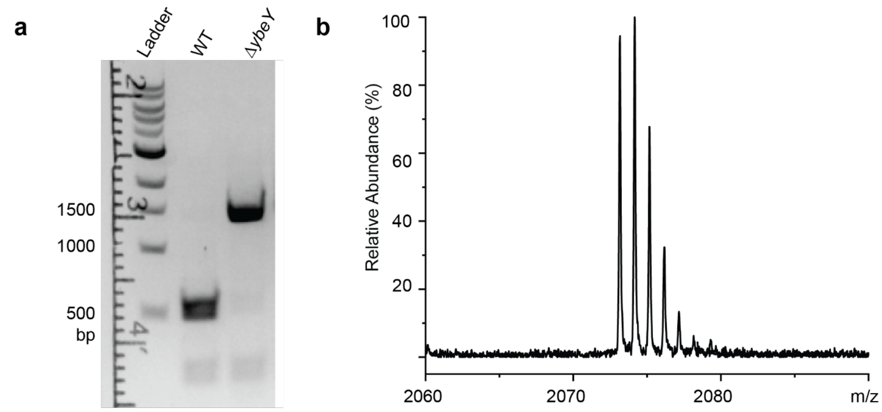


c

| Obs. | Abund.% | Calc. | Abund.% |
|---|---|---|---|
| 2073.2 | 94 | 2073.1 | 90 |
| 2074.2 | 100 | 2074.1 | 100 |
| 2075.2 | 68 | 2075.1 | 64 |
| 2076.2 | 32 | 2076.1 | 30 |
| 2077.2 | 13 | 2077.1 | 11 |
| 2078.2 | 6 | 2078.1 | 4 |

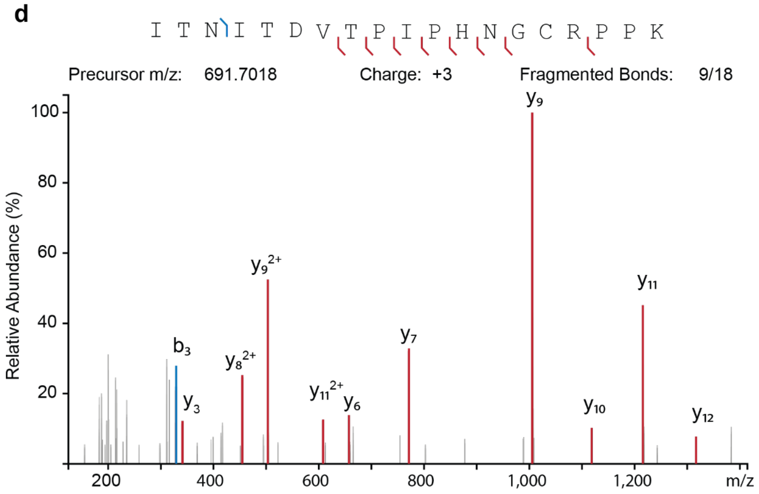


| Ion | Obs. m/z | Calc. m/z | Error (ppm) |
|---|---|---|---|
| b3 | 329.1819 | 329.1819 | 0 |
| y3 | 341.2183 | 341.2181 | 0.6 |
| $y8^{2+}$ | 454.7285 | 454.7296 | -2.4 |
| $y9^{2+}$ | 503.2570 | 503.2560 | 2.0 |
| $y11^{2+}$ | 608.3203 | 608.3244 | -6.7 |
| y6 | 657.3482 | 657.3501 | -2.9 |
| y7 | 771.3988 | 771.3930 | 7.5 |
| y9 | 1005.5065 | 1005.5047 | 1.8 |
| y10 | 1118.5885 | 1118.5888 | -0.3 |
| y11 | 1215.6432 | 1215.6415 | 1.4 |
| y12 | 1316.6922 | 1316.6892 | 2.3 |

**Figure S5. MS analysis of uS11 overexpressed in *E. coli* Δ*ybeY*. a**, Genotype verification by PCR. *ybeY* (468 bp), *ybeY*::*kan*$^R$ (1415 bp). **b-c**, MALDI-TOF MS analysis of trypsin-digested uS11 overexpressed in *E. coli* Δ*ybeY*. Isotopic abundances for the calculated mass distributions of the peptide of interest are indicated in the table. The observed *m/z* is slightly higher than the predicted value (+0.1), likely due to MALDI-TOF calibration variability. Abund., relative abundance. **d**, HRMS/MS spectrum of m/z 691.7018. Assigned ions are indicated in the sequence, spectrum, and table. $Error\ (ppm) = \frac{Observed - Calculated}{Calculated} \times 10^6$, and this formula is used hereafter to calculate errors in the HRMS and HRMS/MS spectra.

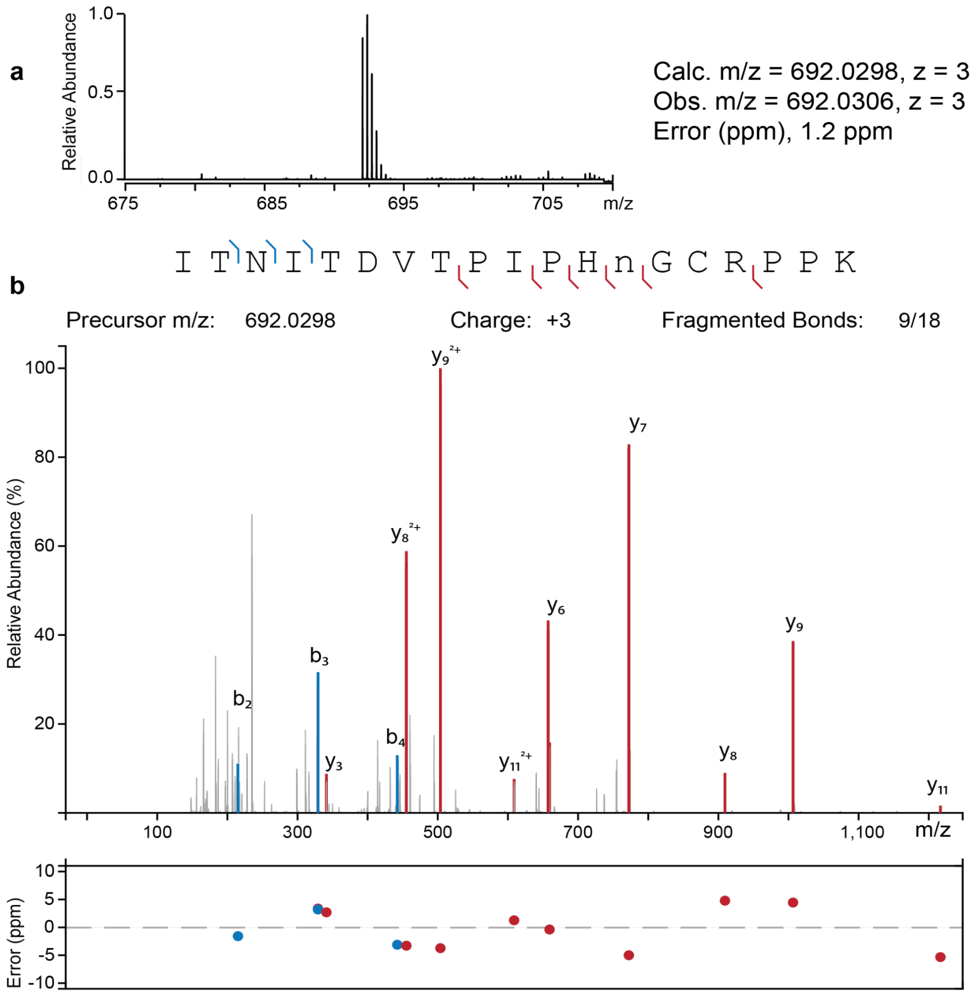


**Figure S6. MS analysis of peptide 1. a**, HRMS of peptide **1**. Calculated and observed m/z values and the mass error (ppm) are labeled. **b**, HRMS/MS spectrum of m/z 692.0298. Assigned ions are indicated in the sequence and spectrum. The mass error (ppm) was plotted at the bottom. n represents the modified Asn residue.

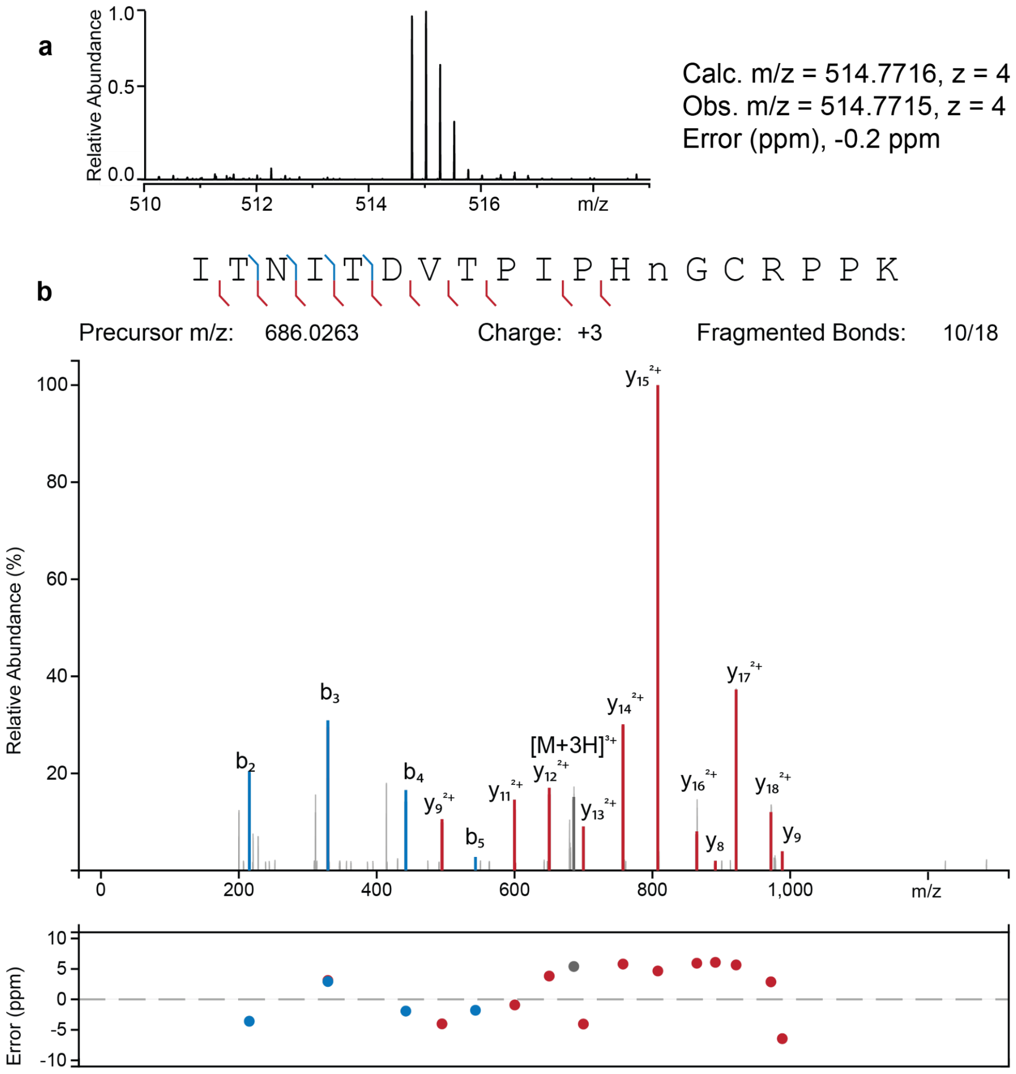


**Figure S7. MS analysis of peptide 2. a**, HRMS of peptide **2**. Calculated and observed m/z values and the mass error (ppm) are labeled. **b**, HRMS/MS spectrum of m/z 686.0263. Assigned ions are indicated in the sequence and spectrum. The mass error (ppm) was plotted at the bottom. n represents the modified Asn residue.

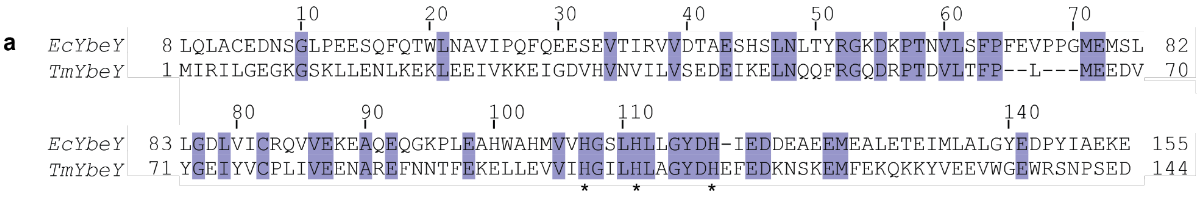


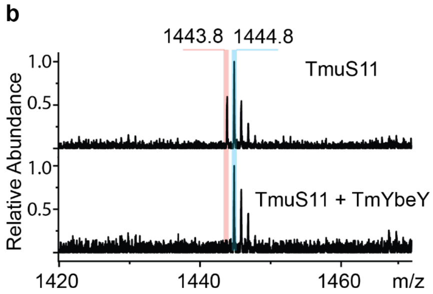


**c**

| Ion | Calc. m/z | Obs. m/z | Error (mDa) |
|---|---|---|---|
| y2 | 244.2 | 244.1 | -100 |
| b3 | 316.2 | 316.2 | 0 |
| y3 | 400.3 | 400.3 | 0 |
| y4 | 503.3 | 503.3 | 0 |
| y5 | 560.3 | 560.3 | 0 |
| y6 | 675.3 | 675.3 | 0 |
| y8 | 919.4 | 919.4 | 0 |
| y10 | 1129.6 | 1129.5 | -100 |
| b11 | 1201.6 | 1201.5 | -100 |
| y12 | 1329.7 | 1329.6 | -100 |

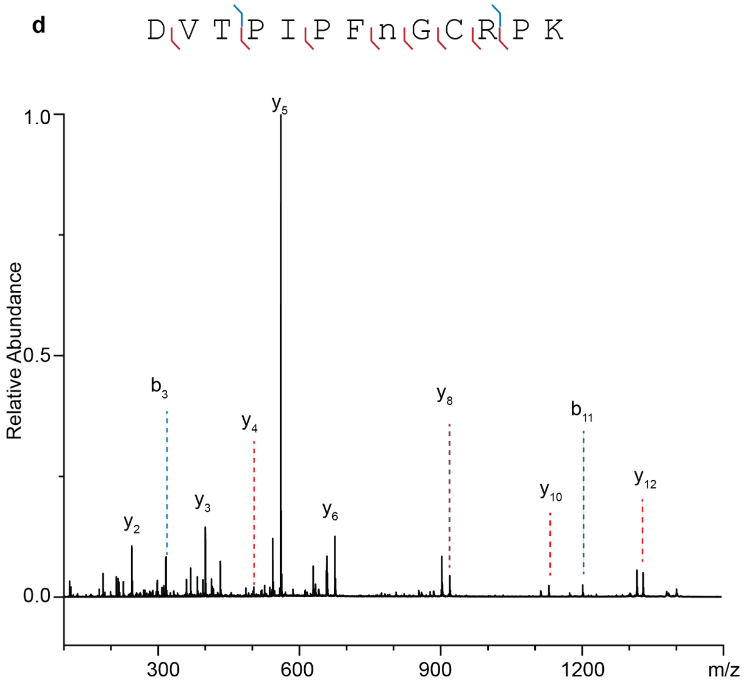


**Figure S8. Analysis of the catalytic activity of *Tm*YbeY. a**, Pairwise alignment of *Ec*YbeY and *Tm*YbeY. Identical residues are colored in purple. Histidine residues for zinc-binding are indicated by stars. **b**, In vitro reaction of *Tm*uS11 substrate (up) and with *Tm*YbeY (bottom). Sequence of interest: DVTPIPFN$_{134}$GCRPK. Formula: $C_{64}H_{102}N_{18}O_{18}S$. $[M+H]^+$ = 1443.7413. Partial modification of *Tm*uS11 was observed after purification, likely due to endogenous YbeY in the BL21(DE3) host. **c**, Table for the assigned ions in the low-resolution MALDI-TOF/TOF spectrum. Mass errors were calculated and reported in mDa. $Error\ (mDa) = (Observed - Calculated) \times 10^3$, and this formula is used hereafter to calculate errors in the MALDI-TOF/TOF spectra. **d**, MALDI-TOF/TOF spectrum of m/z 1444.8. Assigned ions are indicated in the sequence and spectrum. n represents the modified Asn residue.

**a**

1 11 21 31 41 51 61

GSSHHHHHHSQDP MIRILGEGKG SKLLENLKEK LEEIVKKEIG DVHVNVILVS EDEIKELNQQ FRGQDRPTDV LTFPLMEEDV

71 81 91 101 111 121 131 141

YGEIYVCPLI VEENAREFNN TFEKELLEVV IHGILHLAGY DHEFEDKNSK EMFEKQKKYV EEVWGEWRSN PSEDSDPGKR

**b**

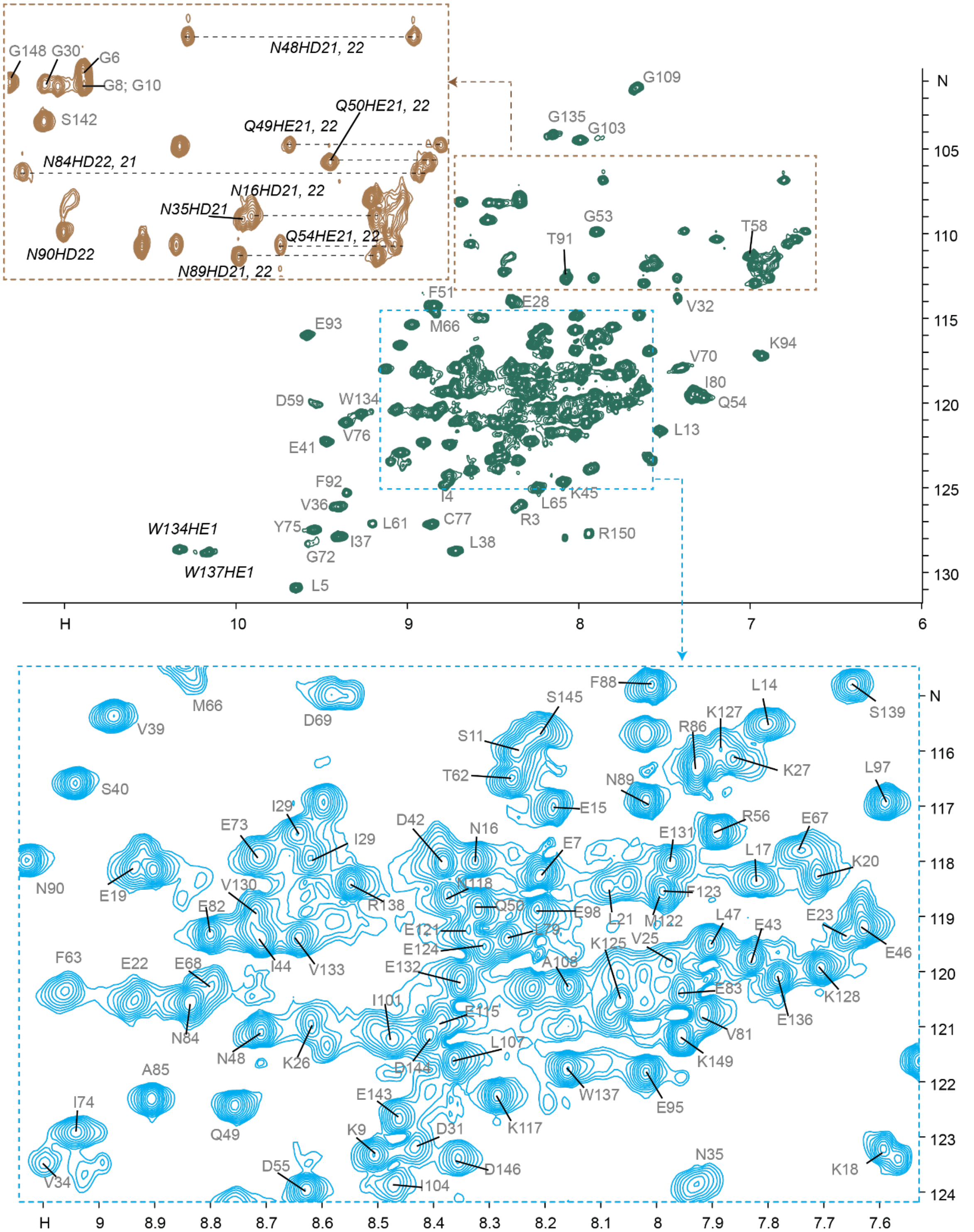

**Figure S9. $^1$H-$^{15}$N HSQC spectrum of apo *Tm*YbeY at 55 °C. a**, Sequence map with backbone-assigned residues highlighted in blue. The underlined region is not part of the native *Tm*YbeY sequence and arises from molecular cloning. **b**, Backbone amides are labeled in gray. Side chain NH signals are labeled in italic black. Two zoomed views are provided.

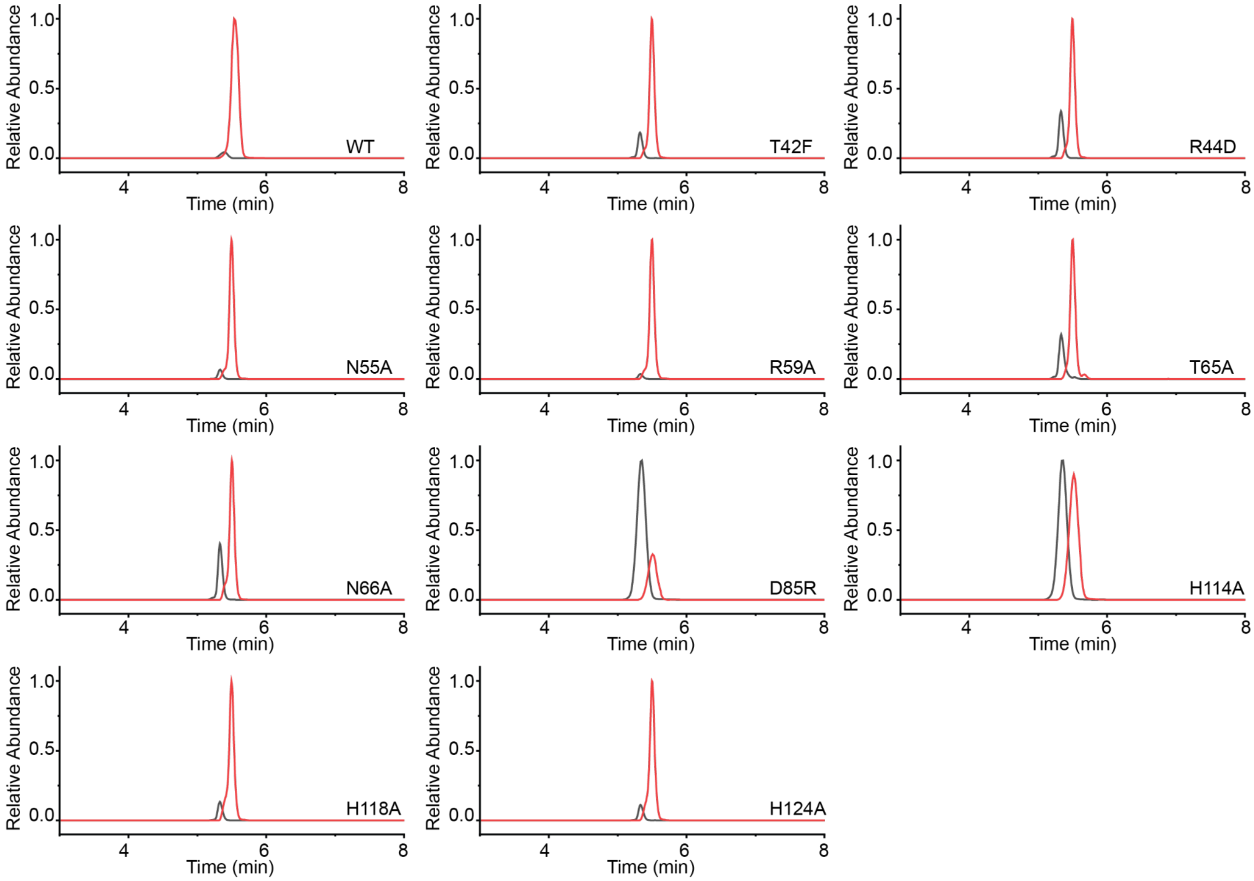


**Figure S10. LC-HRMS spectra of uS11 modification by YbeY wild-type and variants.** Extracted ion chromatograms (EIC) show the unmodified peptide in black and the modified peptide in red. Sequence: `ITNITDVTPIPH`$X_{119}$`GCRPPK`, X = Asn or isoAsp. The exact *m/z* value of 519.0282 is used to extract the unmodified (black) EICs, and 519.2742 is used for the modified (red) EICs.

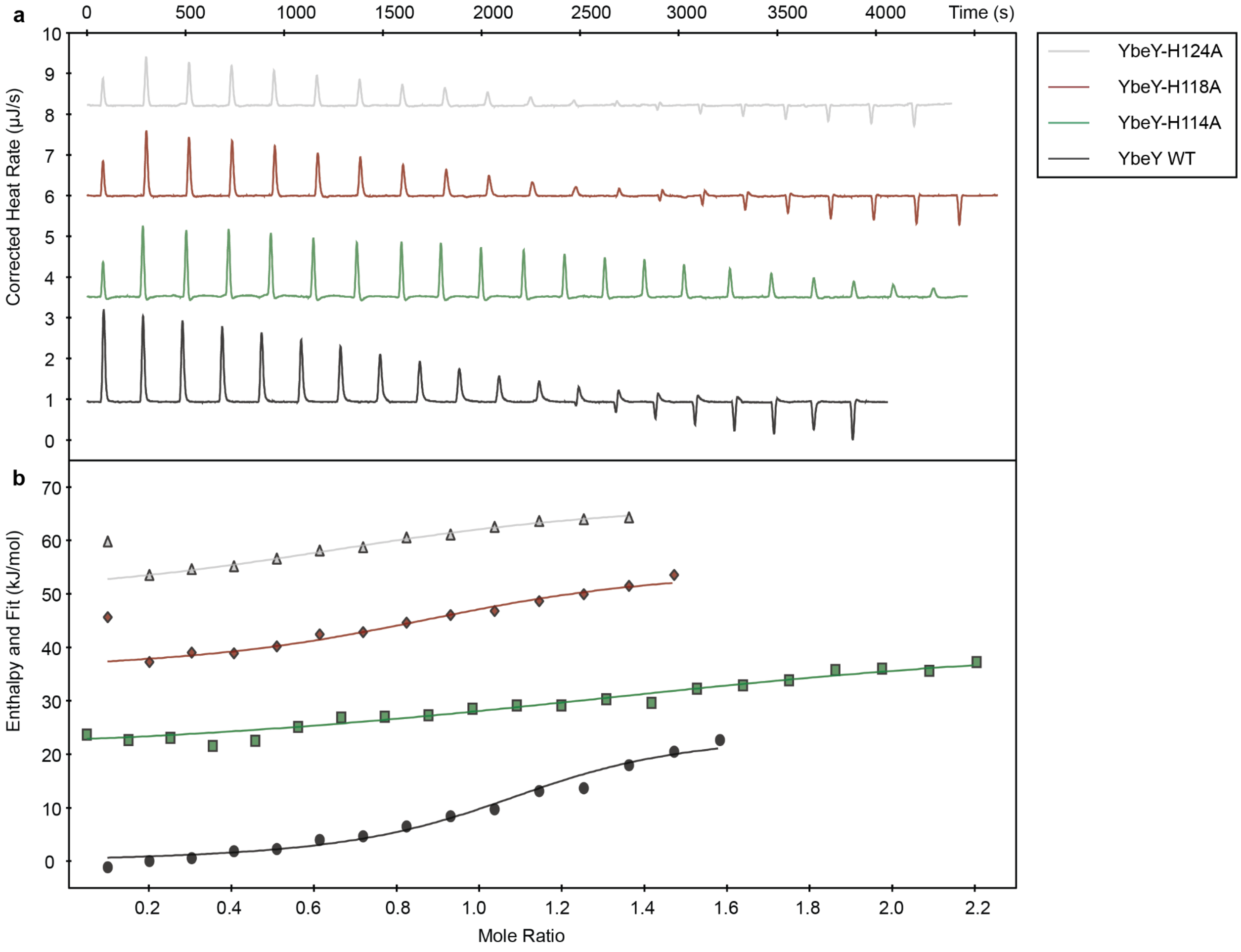


**Figure S11. ITC analysis of YbeY zinc-binding affinity. a,** The heat released after each addition of zinc into the protein solution. The titration of zinc to buffer was used as the blank control and subtracted from each titration with protein. A half-volume was used for the first injection in the ITC experiments with MBP-*Ec*YbeY variants. At higher Zn:protein ratios, the apparent enthalpy shifted from exothermic to endothermic. **b,** The data were fit using a single binding constant to calculate the thermodynamic parameters (continuous line). Late-stage endothermic heats were excluded from fitting. The dissociation constant ($K_D$ ± s.e.) and binding stoichiometry (n): YbeY WT, $K_D$ = 2.3 ± 0.8 µM, n = 1.1; YbeY-H118A, $K_D$ = 5.4 ± 2.8 µM, n = 0.9; YbeY-H124A, $K_D$ = 5.8 ± 2.1 µM, n = 0.9; YbeY-H114A, $K_D$ = 16.8 ± 4.7 µM, n = 1.5.

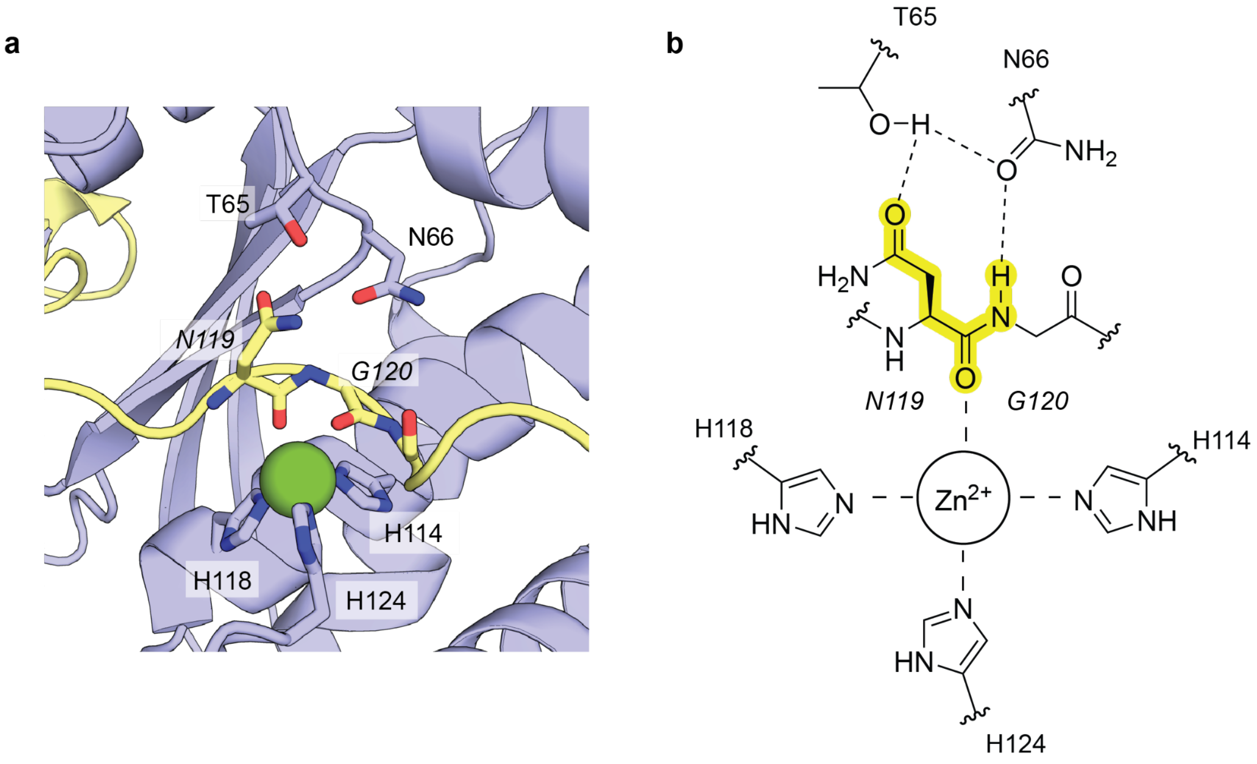


**Figure S12. Zoomed-in AlphaFold3 model of *Ec*uS11 and *Ec*YbeY. a**, Potential interactions of residues T65 and N66 in *Ec*YbeY with the *Ec*uS11 N119 and G120 residues. **b**, Schematic of panel **a**. Potential contacts are indicated by dotted lines.

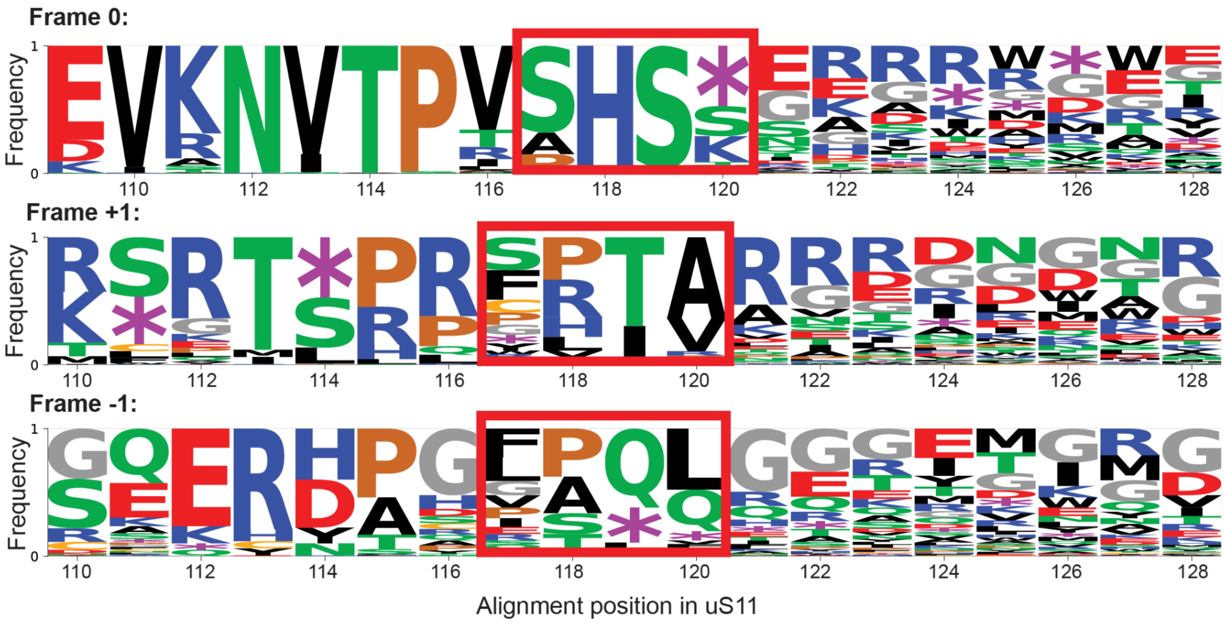


**Figure S13. Shortened C-terminus of Bipolaricaulota uS11**. Nucleotide windows were extracted from Bipolaricaulota genomes ($n$ = 179) at uS11 position 117, localized via the bacterial uS11 HMM, and translated in three reading frames: the natural frame (top), a +1 frameshift (middle), and a −1 frameshift (bottom). Stacked-letter heights give the per-position amino acid frequency, with the x-axis labeled by the *E. coli* uS11 residue positions. The red box marks the four-residue PHNG-equivalent window (HMM alignment positions 107-110; *E. coli* residues 117–120).

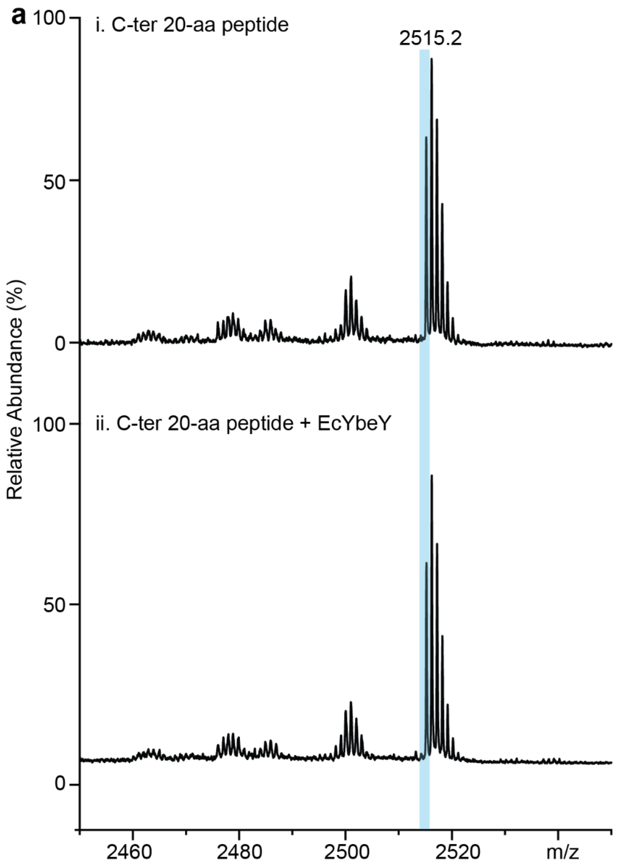


**b**

| Calc. | Abund.% | Obs. (i) | Abund.% (i) |
|---|---|---|---|
| 2515.4 | 75 | 2515.2 | 73 |
| 2516.4 | 100 | 2516.2 | 100 |
| 2517.4 | 75 | 2517.2 | 79 |
| 2518.4 | 40 | 2518.2 | 50 |
| 2519.4 | 17 | 2519.2 | 23 |
| 2520.4 | 6 | 2520.2 | 10 |

**c**

| Calc. | Abund.% | Obs. (ii) | Abund.% (ii) |
|---|---|---|---|
| 2515.4 | 75 | 2515.2 | 70 |
| 2516.4 | 100 | 2516.2 | 100 |
| 2517.4 | 75 | 2517.2 | 77 |
| 2518.4 | 40 | 2518.2 | 45 |
| 2519.4 | 17 | 2519.2 | 21 |
| 2520.4 | 6 | 2520.2 | 9 |

**Figure S14. MALDI-TOF MS analysis of the uS11-Cter peptide**. **a**, MS spectra of C-ter 20-aa peptide substrate (i) and with YbeY (ii). Sequence: SGSITNITDVTPIPHN$_{119}$GCRPPK. The SGS tripeptide at the N-terminus was introduced during cloning. Formula: $C_{107}H_{183}N_{37}O_{31}S$. $[M+H]^+$ = 2515.3674. The calculated and observed mass distributions are listed in the table **b** (i) and **c** (ii). Abund., relative abundance.

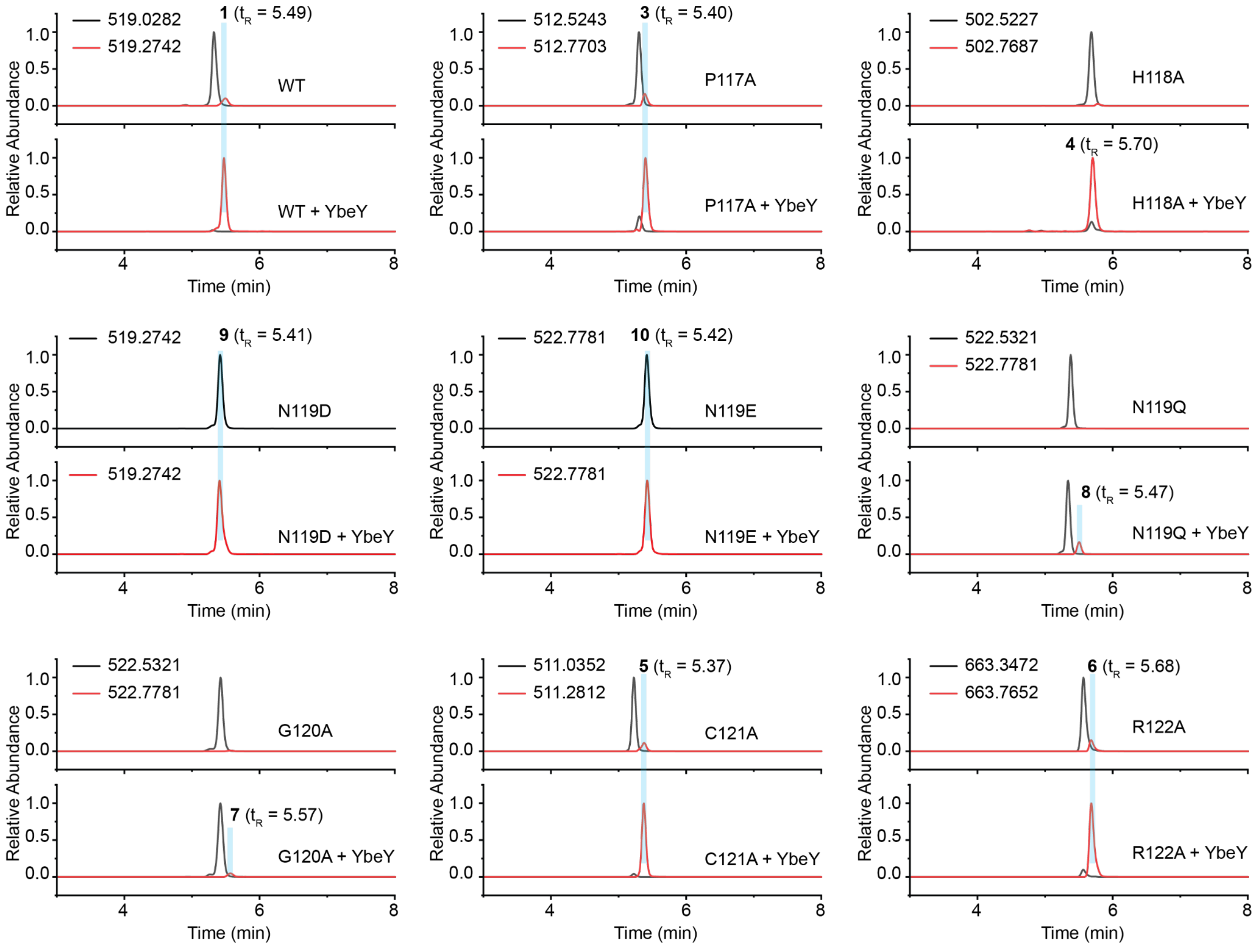


**Figure S15. LC–HRMS spectra of uS11 variants modified by wild-type YbeY.** Extracted ion chromatograms (EICs) show the unmodified peptide in black and the modified peptide in red. Exact m/z values for each EIC are indicated, and the retention time for each peptide is labeled. Trace amounts of modified peptides were detected in uS11 WT and several variants in the reactions without YbeY, likely due to spontaneous formation during protein expression.

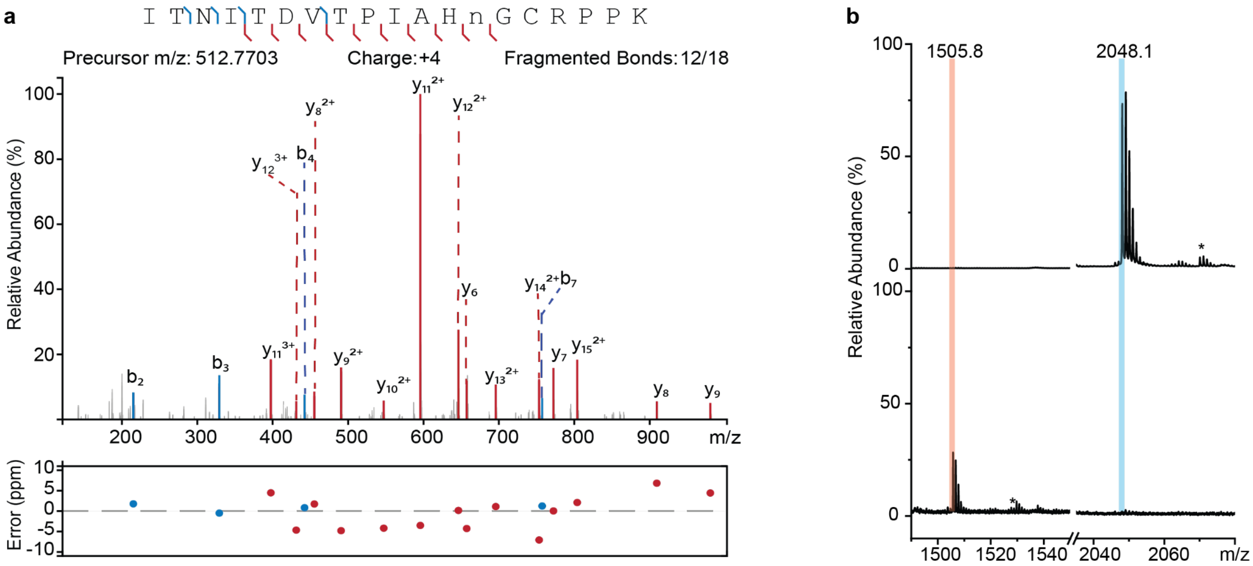


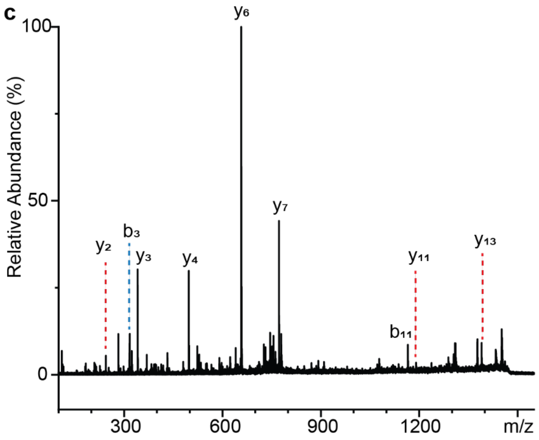


**d**

D V T P I A H n G C R P P K

| Ion | Calc. m/z | Obs. m/z | Error (mDa) |
|---|---|---|---|
| y2 | 244.2 | 244.1 | -100 |
| b3 | 316.2 | 316.1 | -100 |
| y3 | 341.2 | 341.1 | -100 |
| y4 | 497.3 | 497.4 | 100 |
| y6 | 657.4 | 657.4 | 0 |
| y7 | 772.4 | 772.4 | 0 |
| b11 | 1165.4 | 1165.5 | 100 |
| y11 | 1190.6 | 1190.6 | 0 |
| y13 | 1390.7 | 1390.6 | -100 |

**Figure S16. MS analysis of peptide 3. a**, HRMS/MS spectrum of m/z 512.7703 (z = 4). Assigned ions are indicated in sequence and spectrum. The mass error (ppm) was plotted at the bottom. **b**, MALDI-TOF spectra of peptide **3** (upper) and the product of AspN digestion (bottom). Sequence of the product: DVTPIAHn$_{119}$GCRPPK, Formula: $C_{64}H_{104}N_{20}O_{20}S$, $[M+H]^+$ = 1505.7529. $Na^+$ adducts are labeled with asterisks. **c**, MALDI-TOF/TOF spectrum of m/z 1505.8. Assigned ions are indicated in the spectrum and are summarized in **d** (sequence and table). n represents the modified Asn residue.

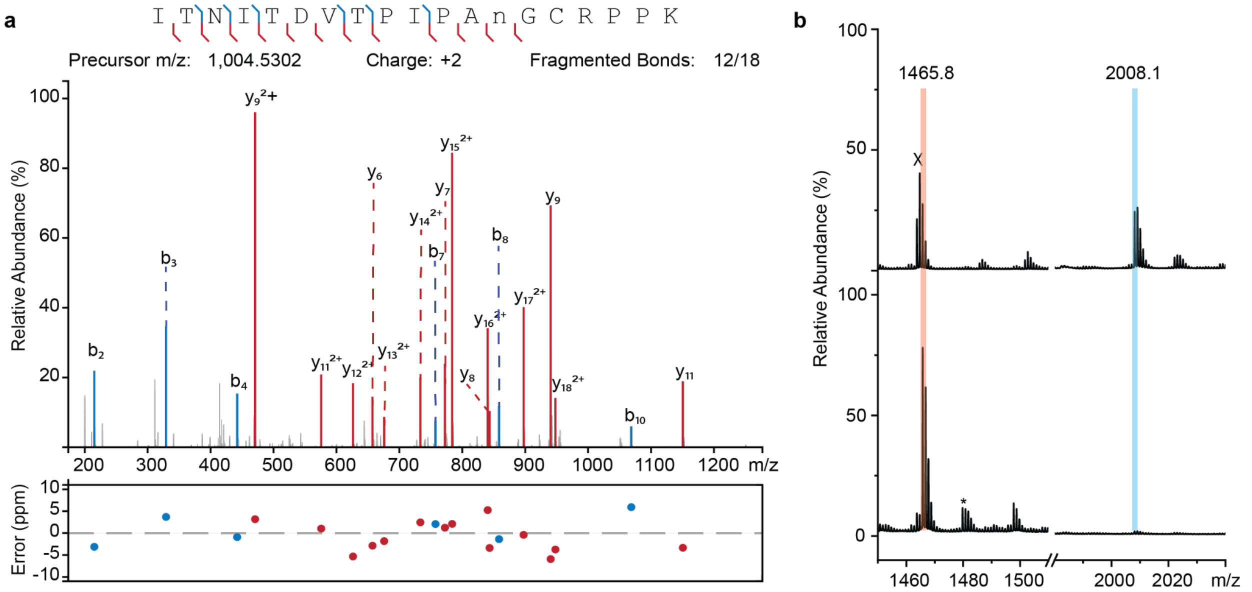


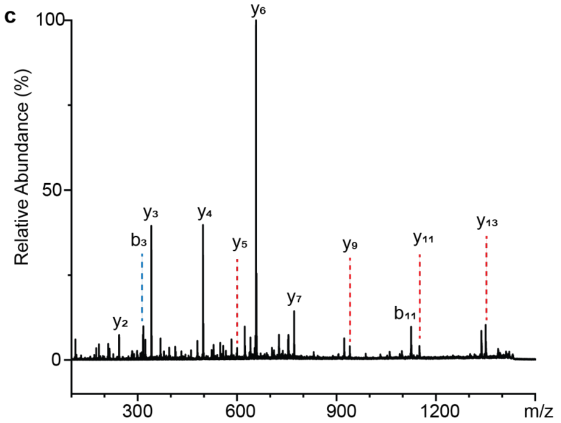


d

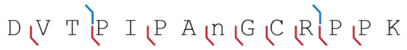


| Ion | Calc. m/z | Obs. m/z | Error (mDa) |
|---|---|---|---|
| y2 | 244.2 | 244.1 | -100 |
| b3 | 316.2 | 316.1 | -100 |
| y3 | 341.2 | 341.1 | -100 |
| y4 | 497.3 | 497.3 | 0 |
| y5 | 600.3 | 600.2 | -100 |
| y6 | 657.4 | 657.2 | -200 |
| y7 | 772.4 | 772.3 | -100 |
| y9 | 940.5 | 940.3 | -200 |
| b11 | 1125.5 | 1125.3 | -200 |
| y11 | 1150.6 | 1150.4 | -200 |
| y13 | 1350.7 | 1350.4 | -300 |

**Figure S17. MS analysis of peptide 4. a**, HRMS/MS spectrum of m/z 1004.5302 (z = 2). Assigned ions are indicated in sequence and spectrum. The mass error (ppm) was plotted at the bottom. **b**, MALDI-TOF spectra of peptide **4** (upper) and the product of AspN digestion (bottom). Sequence of the product: DVTPIPAn$_{119}$GCRPPK, Formula: $C_{63}H_{104}N_{18}O_{20}S$, $[M+H]^+$ = 1465.7467. $Na^+$ adducts are labeled with asterisks. The unrelated peak that overlaps with the 1465 m/z signal is labeled X. **c**, MALDI-TOF/TOF spectrum of m/z 1465.8. Assigned ions are indicated in the spectrum and are summarized in **d** (sequence and table). n represents the modified Asn residue.

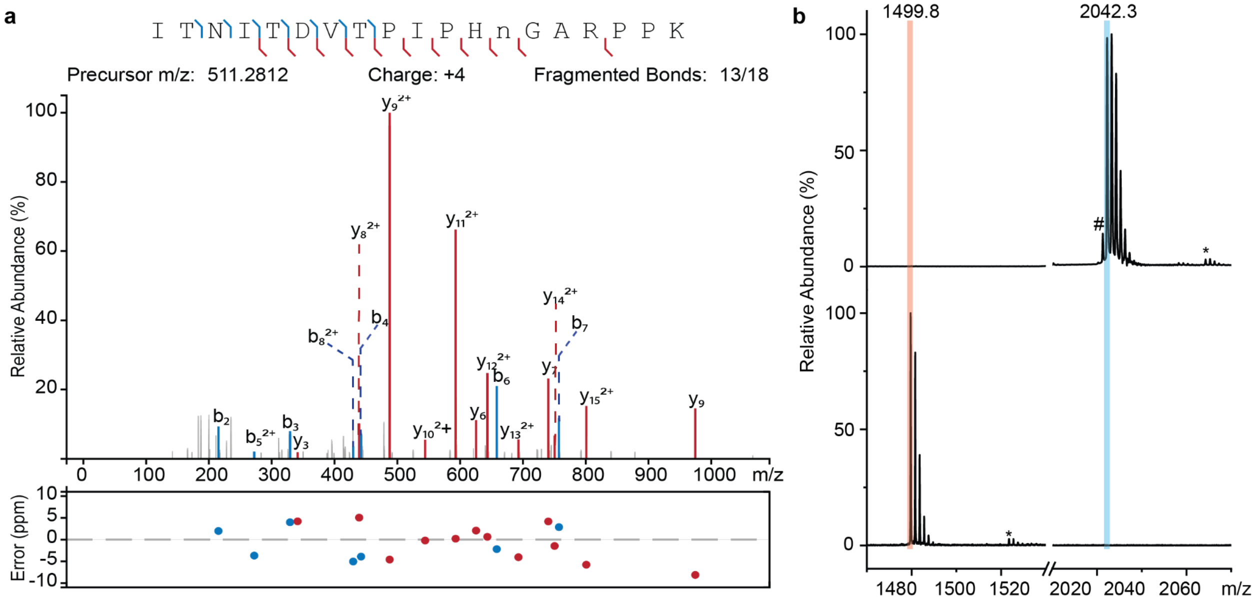


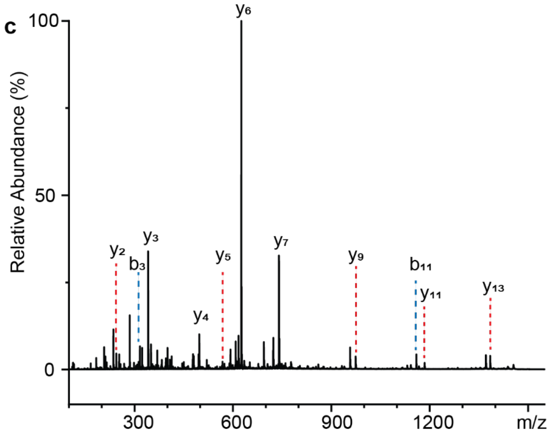


d

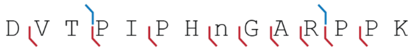


| Ion | Calc. m/z | Obs. m/z | Error (mDa) |
|---|---|---|---|
| y2 | 244.2 | 244.1 | -100 |
| b3 | 316.2 | 316.1 | -100 |
| y3 | 341.2 | 341.1 | -100 |
| y4 | 497.3 | 497.2 | -100 |
| y5 | 568.4 | 568.2 | -200 |
| y6 | 625.4 | 625.2 | -200 |
| y7 | 740.4 | 740.2 | -200 |
| y9 | 974.5 | 974.2 | -300 |
| b11 | 1159.6 | 1159.2 | -400 |
| y11 | 1184.7 | 1184.3 | -400 |
| y13 | 1384.8 | 1384.4 | -400 |

**Figure S18. MS analysis of peptide 5. a**, HRMS/MS spectrum of m/z 511.2812 (z = 4). Assigned ions are indicated in sequence and spectrum. The mass error (ppm) was plotted at the bottom. **b**, MALDI-TOF spectra of peptide **5** (upper) and the product of AspN digestion (bottom). Sequence of the product: DVTPIPHn$_{119}$GARPPK, Formula: $C_{66}H_{106}N_{20}O_{20}$, $[M+H]^+$ = 1499.7965. $Na^+$ adducts are labeled with asterisks. A small amount of unmodified peptide that overlaps with the 2042 m/z signal is labeled #. **c**, MALDI-TOF/TOF spectrum of m/z 1499.8. Assigned ions are indicated in the spectrum and are summarized in **d** (sequence and table). n represents the modified Asn residue.

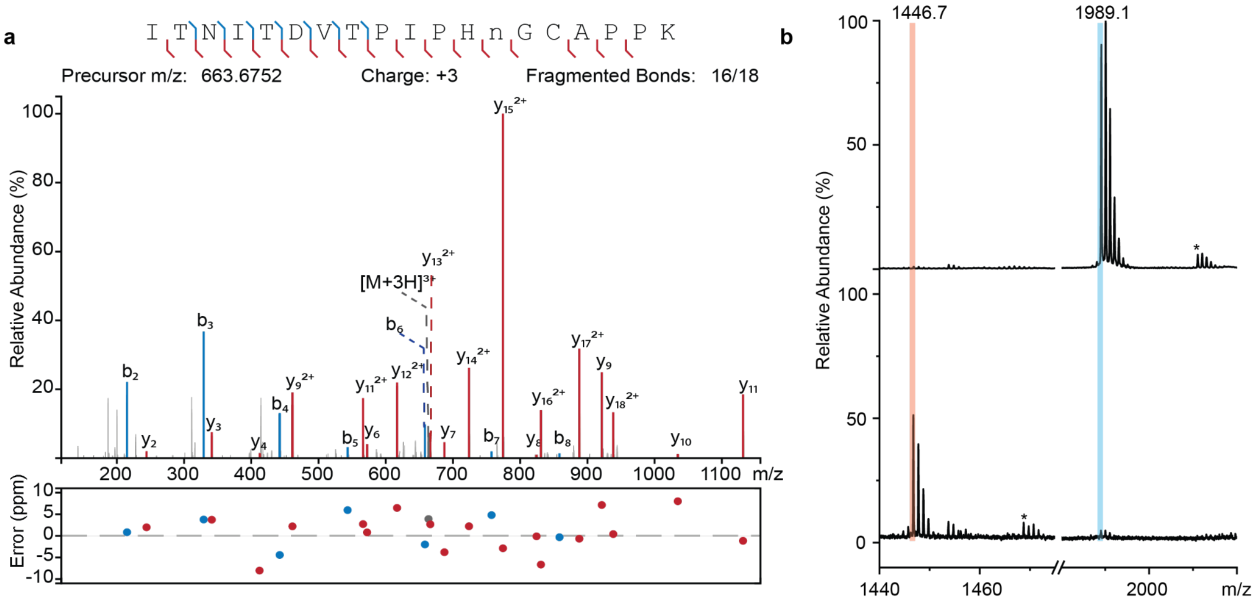


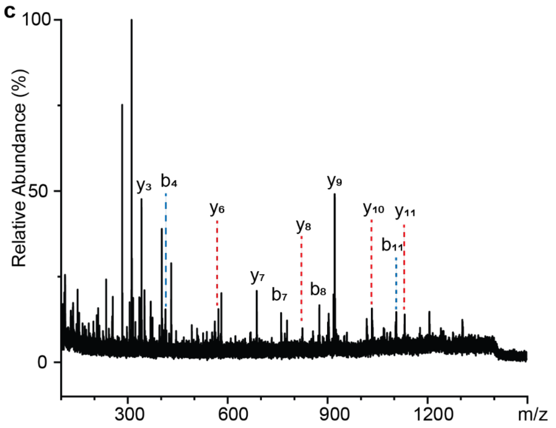


d

D V T P I P H n G C A P P K

| Ion | Calc. m/z | Obs. m/z | Error (mDa) |
|---|---|---|---|
| y3 | 341.2 | 341.1 | -100 |
| b4 | 413.2 | 413.1 | -100 |
| y6 | 572.3 | 572.1 | -200 |
| y7 | 687.3 | 687.1 | -200 |
| b7 | 760.4 | 760.2 | -200 |
| y8 | 824.4 | 824.1 | -300 |
| b8 | 875.4 | 875.2 | -200 |
| y9 | 921.4 | 921.1 | -300 |
| y10 | 1034.5 | 1034.2 | -300 |
| b11 | 1106.5 | 1106.2 | -300 |
| y11 | 1131.6 | 1131.2 | -400 |

**Figure S19. MS analysis of peptide 6. a**, HRMS/MS spectrum of m/z 663.6752 (z = 3). Assigned ions are indicated in sequence and spectrum. The mass error (ppm) was plotted at the bottom. **b**, MALDI-TOF spectra of peptide **6** (upper) and the product of AspN digestion (bottom). Sequence of the product: DVTPIPHn$_{119}$GCAPPK, Formula: $C_{63}H_{99}N_{17}O_{20}S$, $[M+H]^+$ = 1446.7046. $Na^+$ adducts are labeled with asterisks. **c**, MALDI-TOF/TOF spectrum of m/z 1446.7. Assigned ions are indicated in the spectrum and are summarized in **d** (sequence and table). n represents the modified Asn residue.

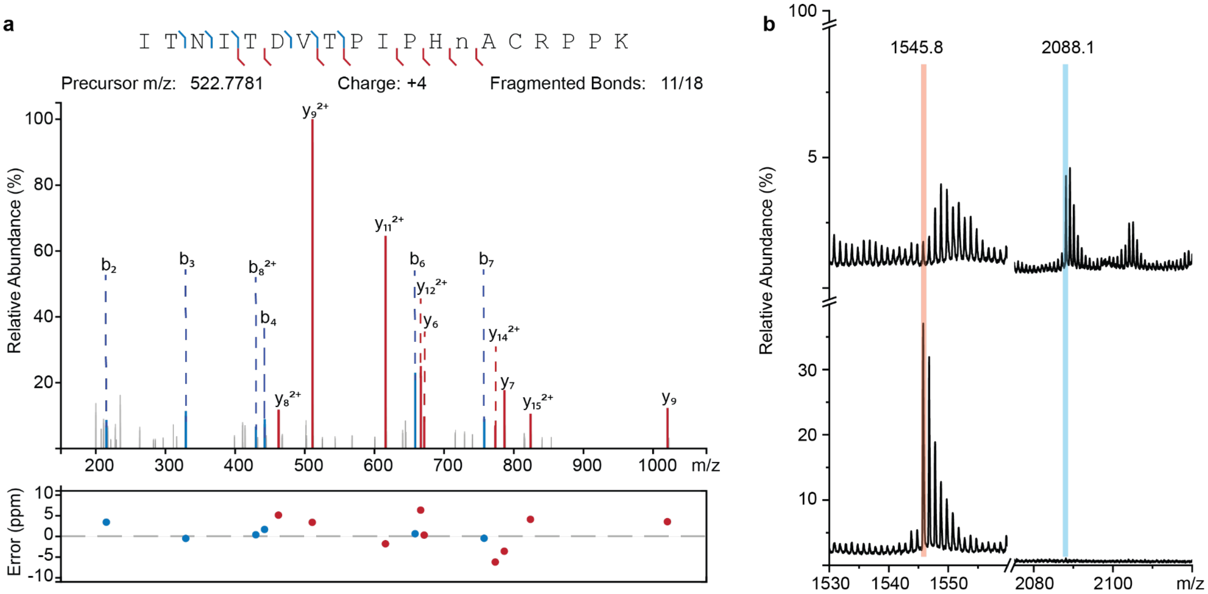


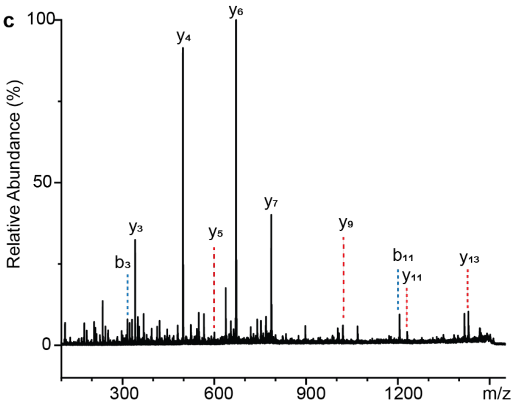


d

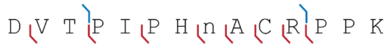


| Ion | Calc. m/z | Obs. m/z | Error (mDa) |
|---|---|---|---|
| b3 | 316.2 | 316.1 | 60 |
| y3 | 341.2 | 341.2 | 70 |
| y4 | 497.3 | 497.2 | 100 |
| y5 | 600.3 | 600.2 | 150 |
| y6 | 671.4 | 671.2 | 180 |
| y7 | 786.4 | 786.3 | 120 |
| y9 | 1020.5 | 1020.3 | 180 |
| b11 | 1205.6 | 1205.3 | 240 |
| y11 | 1230.6 | 1230.3 | 340 |
| y13 | 1430.8 | 1430.4 | 390 |

**Figure S20. MS analysis of peptide 7. a**, HRMS/MS spectrum of m/z 522.7781 (z = 4). Assigned ions are indicated in sequence and spectrum. The mass error (ppm) was plotted at the bottom. **b**, MALDI-TOF spectra of peptide **7** (upper) and the product of AspN digestion (bottom). Sequence of the product: DVTPIPHn$_{119}$ACRPPK, Formula: $C_{67}H_{108}N_{20}O_{20}S$, $[M+H]^+$ = 1545.7842. **c**, MALDI-TOF/TOF spectrum of m/z 1545.8. Assigned ions are indicated in the sequence, table, and spectrum. n represents the modified Asn residue.

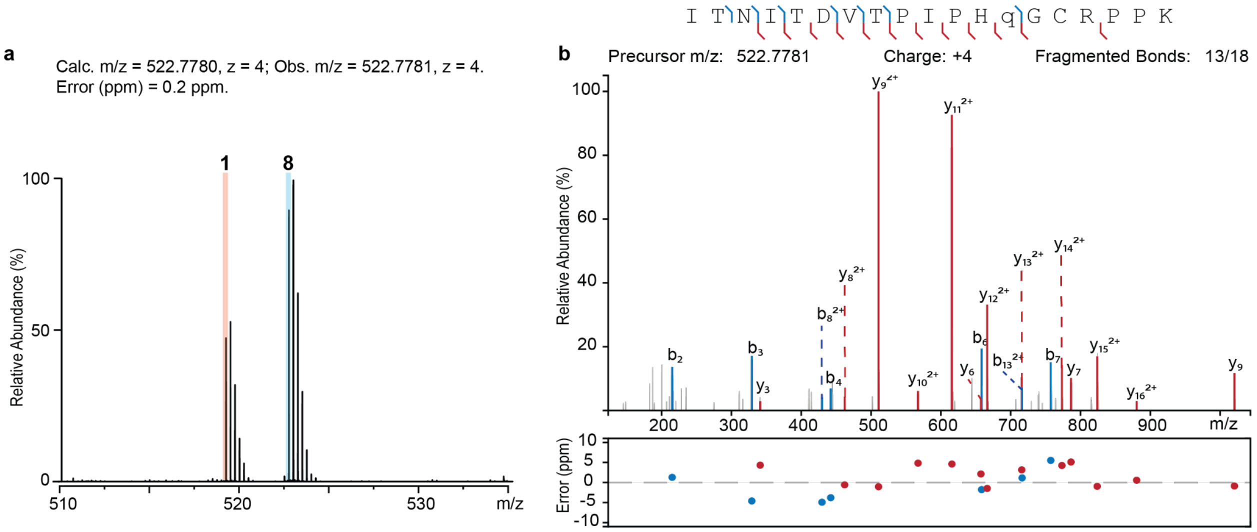


**Figure S21. MS analysis of peptide 8. a**, HRMS of peptide **8**. Peptide **1** co-elutes with peptide **8**. We speculate that peptide **1** originates from a small amount of uS11 WT that co-purified with the YbeY enzyme used for the *in vitro* studies, because this peak was detected in different uS11 variants. **b**, HRMS/MS spectrum of m/z 522.7781 (z = 4). Assigned ions are indicated in sequence and spectrum. The mass error (ppm) was plotted at the bottom. q represents the modified Gln residue.

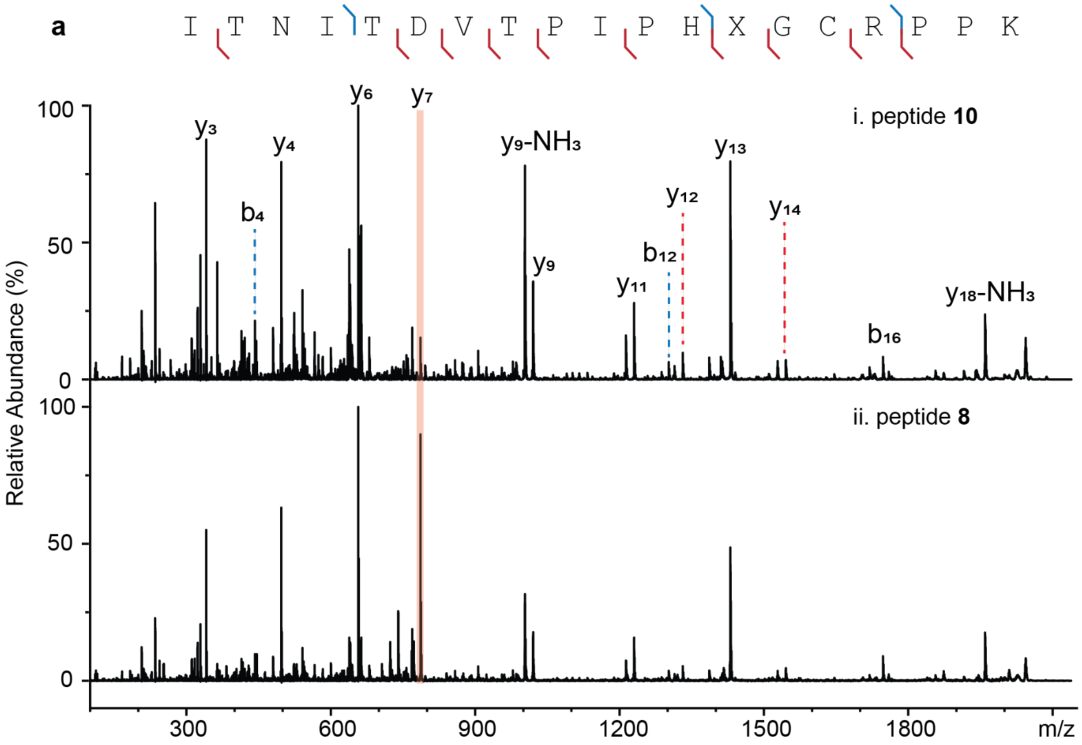


**b**

| Ion | Calc. m/z | Obs. (i, m/z) | Error (i, mDa) | Obs. (ii, m/z) | Error (ii, mDa) |
|---|---|---|---|---|---|
| y3 | 341.2 | 341.1 | -100 | 341.2 | 0 |
| b4 | 442.3 | 442.1 | -200 | 442.2 | -100 |
| y4 | 497.3 | 497.2 | -100 | 497.2 | -100 |
| y6 | 657.4 | 657.1 | -300 | 657.2 | -200 |
| y7 | 786.4 | 786.1 | -300 | 786.2 | -200 |
| y9-$NH_3$ | 1003.5 | 1003.2 | -300 | 1003.2 | -300 |
| y9 | 1020.5 | 1020.2 | -300 | 1020.2 | -300 |
| y11 | 1230.6 | 1230.2 | -400 | 1230.3 | -300 |
| b12 | 1302.7 | 1302.2 | -500 | 1302.3 | -400 |
| y12 | 1331.7 | 1331.2 | -500 | 1331.3 | -400 |
| y13 | 1430.8 | 1430.3 | -500 | 1430.3 | -400 |
| y14 | 1545.8 | 1545.2 | -600 | 1545.3 | -500 |
| b16 | 1747.9 | 1747.3 | -600 | 1747.3 | -600 |
| y18-$NH_3$ | 1958.0 | 1958.3 | -700 | 1958.4 | -600 |

**Figure S22. MALDI-TOF/TOF spectra of peptides 8 and 10. a**, Assigned ions from MALDI-TOF/TOF are denoted on the sequence and spectra. **b**, Table of assigned ions. The mass error (ppm) was plotted at the bottom. i, peptide **10**, X = Glu; ii, peptide **8**, X = isoGlu.

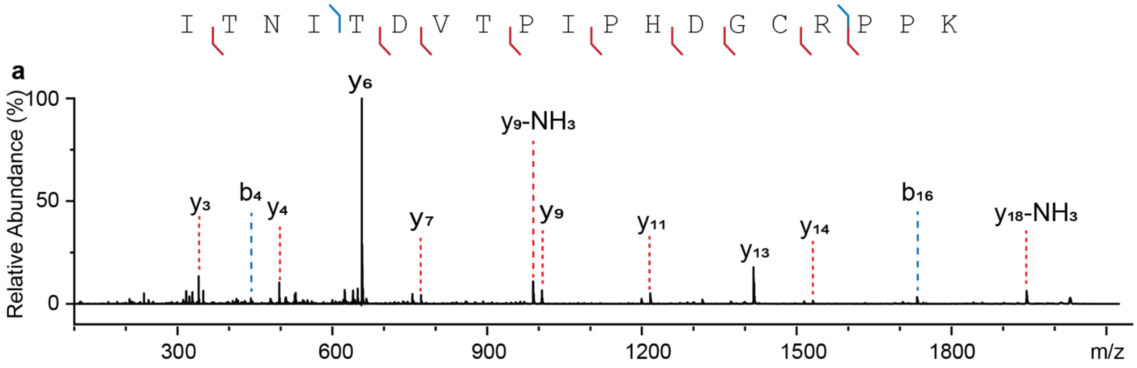


| Ion | Calc. m/z | Obs. m/z | Error (mDa) | Ion | Calc. m/z | Obs. m/z | Error (mDa) |
|---|---|---|---|---|---|---|---|
| y3 | 341.2 | 341.2 | 0 | y9 | 1006.5 | 1006.2 | -300 |
| b4 | 442.3 | 442.2 | -100 | y11 | 1216.6 | 1216.3 | -300 |
| y4 | 497.3 | 497.2 | -100 | y13 | 1416.7 | 1416.3 | -400 |
| y6 | 657.4 | 657.2 | -200 | y14 | 1531.8 | 1531.3 | -500 |
| y7 | 772.4 | 772.2 | -200 | b16 | 1733.9 | 1733.3 | -600 |
| y9-$NH_3$ | 989.5 | 989.2 | -300 | y18-$NH_3$ | 1944.0 | 1943.4 | -600 |

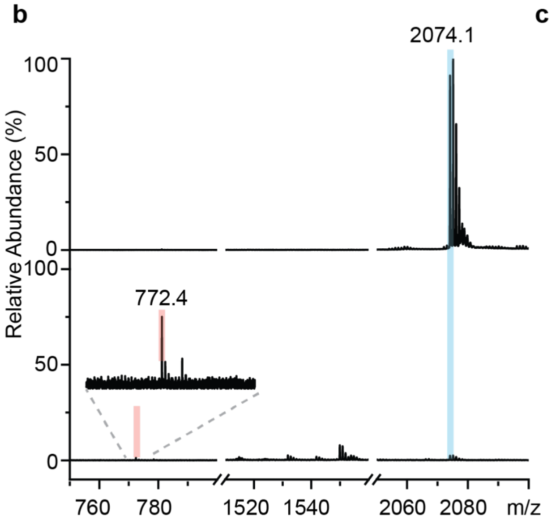


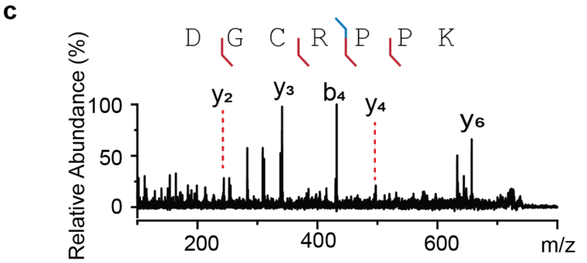


| Ion | Calc. m/z | Obs. m/z | Error (mDa) |
|---|---|---|---|
| y2 | 244.2 | 244.1 | -100 |
| y3 | 341.2 | 341.1 | -100 |
| b4 | 432.2 | 432.1 | -100 |
| y4 | 497.3 | 497.2 | -100 |
| y6 | 657.4 | 657.2 | -200 |

**Figure S23. MS analysis of peptide 9. a**, MALDI-TOF/TOF spectrum of m/z 2074.1. Assigned ions are indicated in the sequence, table, and spectrum. **b**, MALDI-TOF spectra of peptide **9** (upper) and the product of AspN digestion (bottom). Sequence of the product: $D_{119}$GCRPPK, Formula: $C_{31}H_{53}N_{11}O_{10}S$, $[M+H]^+$ = 772.3770. **c**, MALDI-TOF/TOF spectrum of m/z 772.4. Assigned ions are indicated in the sequence, table, and spectrum.

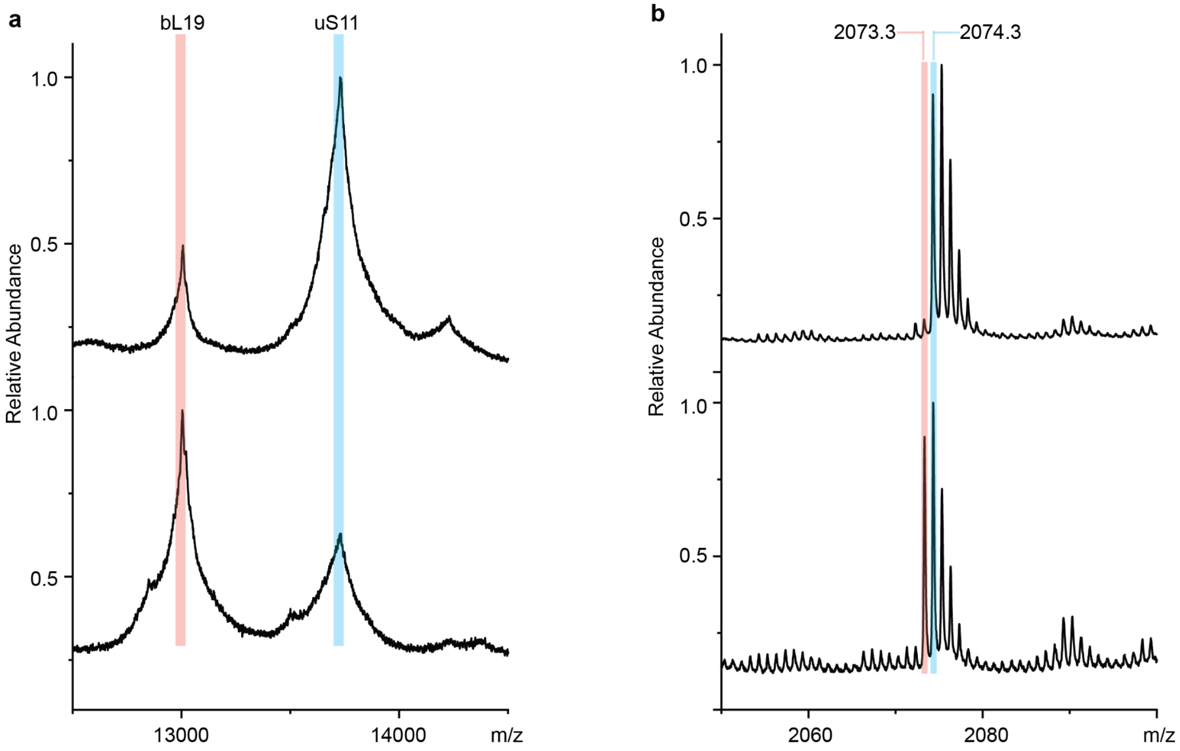


**Figure S24**. **MALDI-TOF MS analysis of uS11 isolated from *E. coli* WT (upper) and the Δ*ybeY* mutant (bottom). a**, Intact ribosomal proteins. bL19 has a molecular weight of 13,002 Da, while uS11 has an expected molecular weight of 13,728 Da, including the removal of Met1 and *N*-methylation of Ala2 modification.[1] **b**, Tryptic peptides of enriched uS11 samples. Observed m/z values are indicated. Sequence of interest: ITNITDVTPIPH$n_{119}$GCRPPK. n = N (unmodified), $[M+H]^+$ = 2073.0909; n = isoAsp (modified), $[M+H]^+$ = 2074.0749.

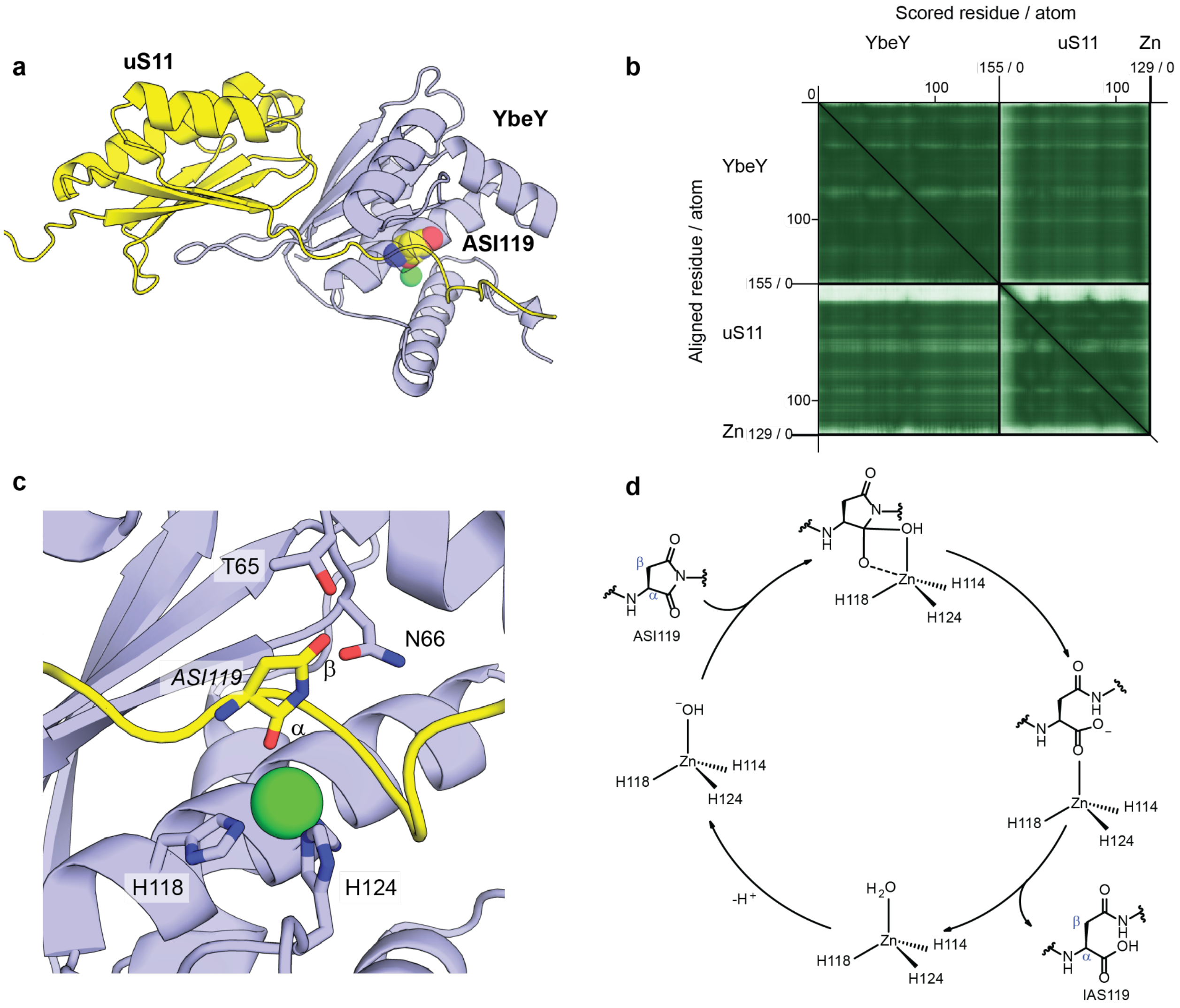


**Figure S25. AlphaFold3 models of *E. coli* YbeY–uS11–aspartimide–zinc. a**, Overall view of the model. *Ec*uS11, *Ec*YbeY, and zinc are shown in yellow, purple, and green, respectively. ipTM = 0.87. The aspartimide residue (ASI119) is shown as a sphere. **b**, PAE error plot for the models shown in **a**. Axis tick marks indicate residue positions in each protein. **c**, Zoomed view of the putative aspartimide intermediate in **a**. **d**, Scheme of hydrolytic preference based on a typical catalytic mechanism of zinc-dependent hydrolases.[2] IAS, isoaspartate.

**Supplementary Schemes.**

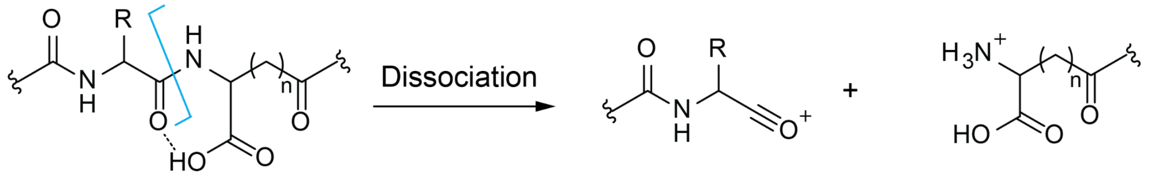


n = 1, isoAsp; n = 2, isoGlu.

b ion

y ion

**Scheme S1. Model for enhanced dissociation on the N-terminal side of isoaspartate (n = 1) or isoglutamate (n = 2).**

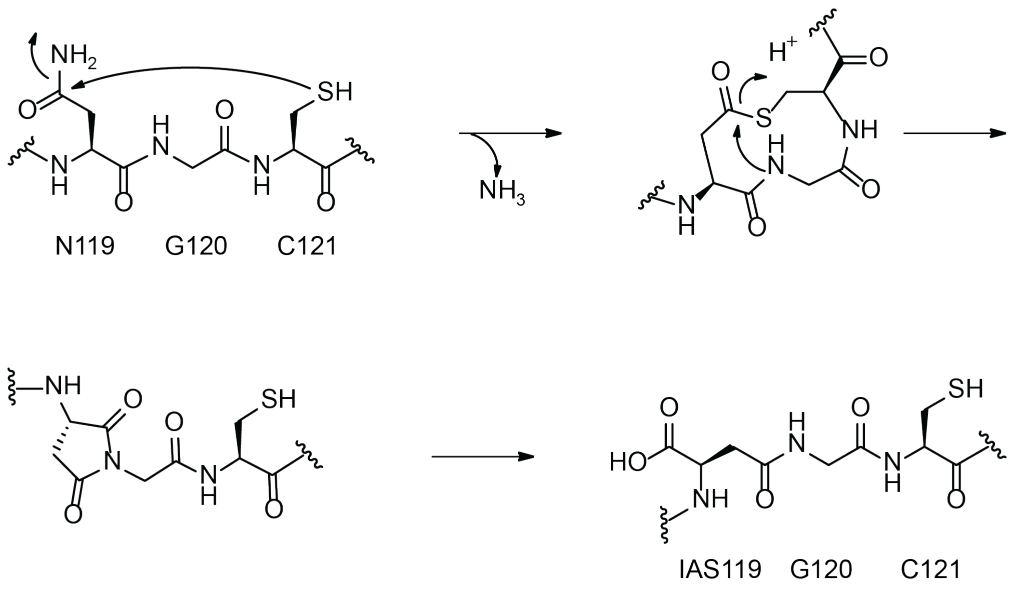


**Scheme S2. A proposed cysteine-engaged mechanism for isoAsp formation.** IAS, isoaspartate.

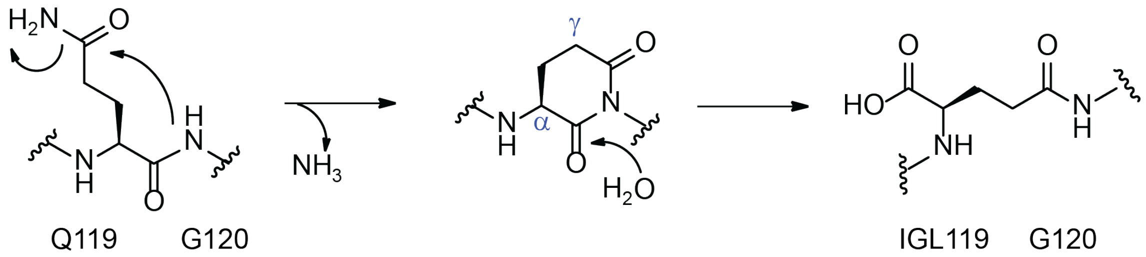


**Scheme S3. A proposed mechanism for isoglutamate formation.** IGL, isoglutamate.

**Supplementary Tables.**

**Table S1 Candidates from AlphaFold3 scanning.**

| UniProt ID | Protein | Function | Mean ipTM |
|---|---|---|---|
| P0AGM5 | YchA | Transglutaminase-like/TPR repeat-containing protein | 0.852 |
| P0A898 | YbeY | Endoribonuclease | 0.847 |
| P68679 | bS21 | 30S ribosomal subunit protein S21 | 0.824 |
| P42615 | MzrA | Modulator protein | 0.789 |
| P0A7G2 | RbfA | 30S ribosome-binding factor | 0.788 |
| P0DPP8 | YsdD | Unannotated small protein | 0.774 |
| P0A8D6 | YmdB | *O*-acetyl-ADP-ribose deacetylase | 0.771 |
| P0DSE9 | YchT | Unannotated small protein | 0.768 |
| P76172 | YnfD | DUF1161 domain-containing protein | 0.750 |
| P27128 | WaaO | Lipopolysaccharide glucosyltransferase | 0.749 |
| P0DSG3 | YqgH | Unannotated small protein | 0.740 |
| P76035 | YciW | Putative peroxidase | 0.736 |
| P03835 | InsG | IS4 putative transposase | 0.731 |
| P0DSG1 | YfiS | Unannotated small protein | 0.722 |
| P42914 | YraI | Putative fimbrial chaperone | 0.715 |
| P77437 | HyfF | Hydrogenase 4 component F | 0.715 |
| P45753 | HofM | DNA utilization protein | 0.714 |
| P77263 | EcpE | Probable fimbrial chaperone | 0.714 |
| P69797 | ManX | Mannose-specific PTS enzyme IIAB component | 0.708 |

| | | | |
|---|---|---|---|
| P0DSH7 | YsgD | Unannotated small protein | 0.707 |
| P0DPM7 | YadW | Unannotated small protein | 0.707 |
| P63204 | GadE | DNA-binding transcriptional activator | 0.706 |
| Q2EET0 | YpdG | Uncharacterized protein | 0.704 |
| P75675 | YkfJ | RNA ligase superfamily protein | 0.702 |
| P67601 | YobD | DUF986 domain-containing inner membrane protein | 0.701 |
| P24193 | HypE | Carbamoyl dehydratase | 0.697 |
| P0AEZ1 | MetF | 5,10-methylenetetrahydrofolate reductase | 0.692 |
| Q46948 | YajL | Protein/nucleic acid deglycase 3 | 0.692 |
| C1P601 | RzoQ | Putative lipoprotein | 0.690 |
| P0AGG4 | TrxC | Thioredoxin 2 | 0.689 |
| P0DPN2 | YkiD | Unannotated small protein | 0.685 |
| P15030 | FecC | Ferric citrate ABC transporter membrane subunit | 0.684 |
| P77184 | LomR | Putative Rac prophage | 0.682 |
| P65556 | YfcD | Putative Nudix hydrolase | 0.680 |
| P0DV23 | YtjE | Unannotated small protein | 0.674 |
| Q46897 | CasE | Pre-CRISPR RNA endonuclease | 0.674 |
| P75684 | YagP | Putative LysR family substrate binding domain-containing protein | 0.671 |
| P0DUW1 | YmgN | Unannotated small protein | 0.671 |
| Q46787 | YgeG | TPR repeat-containing putative chaperone | 0.670 |
| P64461 | LsrG | (4*S*)-4-hydroxy-5-phosphonooxypentane-2,3-dione isomerase | 0.666 |
| P75870 | YccS | Putative transporter | 0.664 |
| P0ABJ1 | CyoA | Cytochrome bo3 subunit 2 | 0.663 |
| P78061 | PuuA | Glutamate-putrescine ligase | 0.660 |
| P0DSH6 | YsdE | Unannotated small protein | 0.655 |

| | | | |
|---|---|---|---|
| P76010 | YcgR | Flagellar brake protein | 0.649 |
| P0DSF7 | YoaM | Unannotated small protein | 0.647 |
| P0AA25 | TrxA | Thioredoxin 1 | 0.642 |
| P0AFV4 | MepS | Peptidoglycan endopeptidase/peptidoglycan L,D-carboxypeptidase | 0.641 |
| P77188 | EcpB | Probable fimbrial chaperone | 0.638 |
| P02359 | uS7 | Small ribosomal subunit protein uS7 | 0.632 |
| P76539 | YpeA | Putative acetyltransferase | 0.630 |
| P07654 | PstA | Phosphate ABC transporter membrane subunit | 0.629 |
| P0DSI0 | YthB | Unannotated small protein | 0.629 |
| P66948 | BepA | Barrel assembly-enhancing protease | 0.627 |
| P76147 | DgcF | Putative diguanylate cyclase | 0.626 |
| P77806 | YbdL | Methionine transaminase | 0.625 |
| P77269 | YphF | Putative ABC transporter periplasmic binding protein | 0.624 |
| P22255 | CysQ | Bisphosphate nucleotidase | 0.623 |
| P0A8E5 | YacL | PF06062 family protein | 0.622 |
| P0A6V5 | GlpE | Thiosulfate sulfurtransferase | 0.617 |
| P64423 | ZntB | $Zn^{2+}$:$H^+$ symporter | 0.617 |
| P76577 | PbpC | Peptidoglycan glycosyltransferase | 0.616 |
| P24205 | LpxM | Lipid A biosynthesis myristoyltransferase | 0.615 |
| P77522 | SufB | Fe-S cluster scaffold complex subunit | 0.614 |
| C1P613 | YqeL | Uncharacterized small protein | 0.608 |
| P33927 | CcmF | Holocytochrome c synthase CcmF component | 0.605 |
| P37641 | YhjC | Putative DNA-binding transcriptional regulator | 0.603 |
| P21866 | KdpE | Putative DNA-binding transcriptional dual regulator | 0.602 |

**Table S2 Cryo-EM data collection, refinement, and validation statistics**

| | ΔYbeY 30S subunit (EMDB-78572) (PDB 37WT) |
|---|---|
| **Data collection and processing** | |
| Magnification | 136,200 |
| Voltage (kV) | 300 |
| Electron exposure (e–/Å$^2$) | 40 |
| Defocus range (μm) | -0.5/-1.5 |
| Pixel size (Å) | 0.6395 |
| Symmetry imposed | C1 |
| Initial particle images (no.) | 567,692 |
| Final particle images (no.) | 332,514 |
| Map resolution (Å) | 1.91 |
| FSC threshold | 0.143 |
| **Refinement** | |
| Initial model used (PDB code) | 8EMM |
| Model resolution (Å) | 2.1 |
| FSC threshold | 0.5 |
| Map sharpening B factor (Å$^2$) | -47.3 |
| Model composition | |
| Non-hydrogen atoms | 53,369 |
| Waters | 1,707 |
| $Mg^{2+}$ ions | 84 |
| B factors (Å$^2$) | |
| RNA | 3.09 |
| Protein | 4.46 |
| Waters | 2.43 |
| Ligand | 3.44 |
| R.m.s. deviations | |
| Bond lengths (Å) | 0.006 |

| | |
|---|---|
| Bond angles (°) | 0.852 |
| Validation | |
| MolProbity score | 2.29 |
| Clashscore | 9.58 |
| Poor rotamers (%) | 3.57 |
| Ramachandran plot | |
| Favored (%) | 94.66 |
| Allowed (%) | 5.21 |
| Disallowed (%) | 0.13 |
| RNA validation | |
| Angle outliers (%) | 0.006 |
| Sugar pucker outliers (%) | 0.729 |
| Average suiteness | 0.509 |

**Table S3 Oligonucleotide primers used in this study.** F, forward primer; R, reverse primer; Capitalized letters, mutagenized codon. *Ec*, *Escherichia coli*; *Tm*, *Thermotoga maritima*.

| **Name** | **Sequence (5′ to 3′)** |
|---|---|
| *Ec*uS11_F | catcaccacagccaggatccgatggcaaaggcaccaattcgtgc |
| *Ec*uS11_R | gcggtttctttaccagacttatacgcgacgttttttcggcggac |
| *Ec*YbeY_F | gaacctgtacttccaatccggatccatgagtcaggtgatcctcgatttac |
| *Ec*YbeY_R | gtgctcgagtgcggccgcttattctttctcggcaatgtacggatc |
| *Ec*YbeY_T42F_F | atcggaagtgTTCattcgcgtggtcgataccgccgaaag |
| *Ec*YbeY_T42F_R | ccacgcgaatGAAcacttccgattcttcctgaaactgc |
| *Ec*YbeY_R44D_F | agtgacgattGATgtggtcgataccgccgaaagccacag |
| *Ec*YbeY_R44D_R | tatcgaccacATCaatcgtcacttccgattcttcctg |
| *Ec*YbeY_N55A_F | ccacagtctgGCCctgacctatcgcggtaaggataagc |
| *Ec*YbeY_N55A_R | gataggtcagGGCcagactgtggctttcggcggtatcg |
| *Ec*YbeY_R59A_F | tctgacctatGCCggtaaggataagccgaccaacgtgc |
| *Ec*YbeY_R59A_R | tatccttaccGGCataggtcagattcagactgtggctttc |
| *Ec*YbeY_T65A_F | ggataagccgGCCaacgtgctctccttcccgtttgaag |
| *Ec*YbeY_T65A_R | agagcacgttGGCcggcttatccttaccgcgataggtcagattcag |
| *Ec*YbeY_N66A_F | taagccgaccGCCgtgctctccttcccgtttgaagtgc |
| *Ec*YbeY_N66A_R | aggagagcacGGCggtcggcttatccttaccgcgatag |
| *Ec*YbeY_D85R_F | gctactgggcCGTctggttatctgccgtcaggtggttg |
| *Ec*YbeY_D85R_R | agataaccagACGgcccagtagcgacatttccatgccag |
| *Ec*YbeY_H114A_F | tatggtggtgGCCggcagtctgcatttgttaggttacg |
| *Ec*YbeY_H114A_R | gcagactgccGGCcaccaccatatgcgcccagtgcgcc |
| *Ec*YbeY_H118A_F | cggcagtctgGCCttgttaggttacgatcacatcgaagatgac |
| *Ec*YbeY_H118A_R | aacctaacaaGGCcagactgccgtgcaccaccatatgc |
| *Ec*YbeY_H124A_F | aggttacgatGCCatcgaagatgacgaagcagaagaaatg |
| *Ec*YbeY_H124A_R | catcttcgatGGCatcgtaacctaacaaatgcagactgc |
| *Ec*uS11_P117A_F | gactccgatcGCCcataacggttgtcgtccgccgaaaaaac |

| | |
|---|---|
| *Ec*uS11_P117A_R | aaccgttatgGGCgatcggagtcacatcagtaatgttag |
| *Ec*uS11_H118A_F | tccgatccctGCCaacggttgtcgtccgccgaaaaaac |
| *Ec*uS11_H118A_R | gacaaccgttGGCagggatcggagtcacatcagtaatg |
| *Ec*uS11_N119A_F | gatccctcatGCCggttgtcgtccgccgaaaaaacgtc |
| *Ec*uS11_N119A_R | gacgacaaccGGCatgagggatcggagtcacatcagtaatg |
| *Ec*uS11_N119D_F | gatccctcatGATggttgtcgtccgccgaaaaaacgtc |
| *Ec*uS11_N119D_R | gacgacaaccATCatgagggatcggagtcacatcagtaatg |
| *Ec*uS11_N119Q_F | gatccctcatCAGggttgtcgtccgccgaaaaaacgtc |
| *Ec*uS11_N119Q_R | gacgacaaccCTGatgagggatcggagtcacatcagtaatg |
| *Ec*uS11_N119E_F | gatccctcatGAAggttgtcgtccgccgaaaaaacgtc |
| *Ec*uS11_N119E_R | gacgacaaccTTCatgagggatcggagtcacatcagtaatg |
| *Ec*uS11_G120A_F | ccctcataacGCCtgtcgtccgccgaaaaaacgtcgcg |
| *Ec*uS11_G120A_R | gcggacgacaGGCgttatgagggatcggagtcacatcag |
| *Ec*uS11_C121A_F | tcataacggtGCCcgtccgccgaaaaaacgtcgcgtataag |
| *Ec*uS11_C121A_R | tcggcggacgGGCaccgttatgagggatcggagtcacatc |
| *Ec*uS11_R122A_F | taacggttgtGCCccgccgaaaaaacgtcgcgtataag |
| *Ec*uS11_R122A_R | ttttcggcggGGCacaaccgttatgagggatcggagtc |
| *EcΔybeY:Kan*_F | gctggcagcagaacgcaagcgcgaagaacaggaacaaaaatgagtcaggtggatccgtcgacc tgcagttc |
| *EcΔybeY:Kan*_R | gttaatcaccaacggcggggacgtctgccagtcaaatgcctggcaaattagtgtaggctggagctgc ttc |
| *EcΔybeY*_Check_F | caagcgcgaagaacaggaac |
| *EcΔybeY*_Check_R | ctgccagtcaaatgcctggc |
| *Ec*uS11_F1 (pBAD24) | ggctagcaggaggaattcacatgggcagcagccatcacc |
| *Ec*uS11_R1 (pBAD24) | ctcatccgccaaaacagccattatacgcgacgtttttcggcg |
| *Ec*YbeY_F1 (pBAD24) | gggctagcaggaggaattcacatgagtcaggtgatcctcgatttacaac |
| *Ec*YbeY_R1 (pBAD24) | tctcatccgccaaaacagccattattctttctcggcaatgtacggatcc |
| *Tm*uS11_F | accatcatcaccacagccaggatccgatggcgcgcaaacgcggcg |
| *Tm*uS11_R | cgcagcagcggtttctttaccagacttacacgcgacggcgtttttc |

| | |
|---|---|
| *Tm*YbeY_F | catcatcaccacagccaggatccgatgattcgcattctgggcgaag |
| *Tm*YbeY_R | cgcagcagcggtttctttaccagacttagcgtttgcccggatcgc |

**Table S4 *E. coli* strains used in this study.**

| Name | Characteristics | Source |
|---|---|---|
| DH5a | Host for general cloning | NEB |
| BL21(DE3) | Host for general protein expression | NEB |
| MRE600 | Parental strain for ribosome assembly assays | Laboratory collection |
| K-12 BW25113 | Parental strain for gene deletion | Laboratory collection |
| K-12 BW25113 Δ*ybeY:Kan* | In-frame deletion of *ybeY* | This work |
| K-12 BW25113 Δ*ybeY:Kan* + pYbeY | Trans-complementation of YbeY using a pBAD24-derived plasmid | This work |
| K-12 BW25113 Δ*ybeY:Kan* + pBAD24 | Empty pBAD24 plasmid control | This work |
| K-12 BW25113 Δ*ybeY:Kan* + pYbeY-R59A | Trans-complementation of YbeY-R59A using a pBAD24-derived plasmid | This work |
| K-12 BW25113 Δ*ybeY:Kan* + pYbeY-D85A | Trans-complementation of YbeY-D85A using a pBAD24-derived plasmid | This work |
| K-12 BW25113 Δ*ybeY:Kan* + pYbeY-H114A | Trans-complementation of YbeY-H114A using a pBAD24-derived plasmid | This work |

**Supplementary References**.

1. Arnold, R. J. & Reilly, J. P. Observation of *Escherichia coli* Ribosomal Proteins and Their Posttranslational Modifications by Mass Spectrometry. *Anal. Biochem.* **269**, 105–112 (1999).
2. Zastrow, M. L. & Pecoraro, V. L. Designing Hydrolytic Zinc Metalloenzymes. Biochemistry ***53***, 957-978 (2014).
